# MOSHPIT: accessible, reproducible metagenome data science on the QIIME 2 framework

**DOI:** 10.1101/2025.01.27.635007

**Authors:** Michal Ziemski, Liz Gehret, Anthony Simard, Santiago Castro Dau, Vinzent Risch, Doriela Grabocka, Christos Matzoros, Colin Wood, Paula Momo Cabrera, Rodrigo Hernández-Velázquez, Chloe Herman, Keegan Evans, Michael S. Robeson, Evan Bolyen, J. Gregory Caporaso, Nicholas A. Bokulich

## Abstract

Metagenome sequencing has revolutionized functional microbiome analysis across diverse ecosystems, but is fraught with technical hurdles. We introduce MOSHPIT (https://moshpit.readthedocs.io), software built on the QIIME 2 framework (Q2F) that integrates best-in-class CAMI2-validated metagenome tools with robust provenance tracking and multiple user interfaces, enabling streamlined, reproducible metagenome analysis for all expertise levels. By building on Q2F, MOSHPIT enhances scalability, interoperability, and reproducibility in complex workflows, democratizing and accelerating discovery at the frontiers of metagenomics.

## Main text

Untargeted next-generation DNA sequencing has revolutionized microbiome research, yielding unparalleled insight into the composition, functionality, and interactions of microbial communities in diverse global ecosystems. The most common technique, marker-gene amplicon sequencing (e.g., of 16S rRNA genes), is powerful and widely used for its scalability and flexibility, but provides a view of the microbial community that is contingent on PCR primer choice, and enables only indirect inference of functional potential. Untargeted whole metagenome (a.k.a., shotgun metagenome) sequencing (WMS) provides several advantages: it facilitates comprehensive community profiling of both functional gene and taxonomic composition; supports sub-species-level taxonomic resolution; and enables assembly of metagenome-assembled genomes (MAGs) without the need for cultivation^1^. Numerous tools exist for different aspects of WMS analysis, but the complex workflows involved, lack of standardization, and issues with interoperability complicate analysis and hamper reproducibility^2^.

Since its official release in 2018^3^, the QIIME 2 microbiome bioinformatics platform has gained a wide user base for its accessibility, interoperability, and reproducibility. However, the Q2F ecosystem has historically lacked full support for untargeted WMS applications. Here, we introduce MOSHPIT (**MO**dular **SH**otgun metagenome **P**ipelines with **I**ntegrated provenance **T**racking), a metagenomics software suite built on the Q2F supporting flexible, customizable, end-to-end WMS analysis. MOSHPIT includes two core plugins and several optional accessory plugins that further enhance analysis. The q2-assembly plugin (https://github.com/bokulich-lab/q2-assembly) provides a set of actions for contig assembly, evaluation, and read mapping, while q2-annotate (https://github.com/bokulich-lab/q2-annotate) supports downstream analysis steps for WMS data, including contig binning, taxonomic classification, and functional annotation. Optional accessory plugins currently include q2-viromics (https://github.com/bokulich-lab/q2-viromics) and q2-virsorter2 (https://github.com/bokulich-lab/q2-virsorter2) for detection and identification of viral sequences in WMS data; and q2-amrfinderplus (https://github.com/bokulich-lab/q2-amrfinderplus) and q2-rgi (https://github.com/bokulich-lab/q2-rgi) for annotation of antimicrobial resistance genes in assembled metagenomes via the popular AMRFinderPlus^4^ and RGI/CARD tools^5^. To increase interoperability between these and other plugins, new data types, formats, and transformers have been added to Q2F to facilitate streamlined analysis workflows for WMS datasets (https://github.com/qiime2/q2-types).

MOSHPIT integrates state-of-the-art tools selected from the top-performing methods in the CAMI2 benchmarks^6^ and other methods to enable customizable fully reproducible end-to-end WMS analysis workflows. This includes : MEGAHIT^7^ and metaSPAdes^8^ for genome assembly; Bowtie2^9^ for read mapping; MetaBAT 2^10^ for contig binning; QUAST^11^ and BUSCO^12^ for quality control of contigs and genomes; and both read- and assembly-based taxonomic classification and functional annotation workflows via Kraken 2^13^, Kaiju^14^, and eggNOG-mapper^15^. (see Figure 1A for an overview of the available workflows and Supplementary Figures 1-3 for detailed sub-workflows).

**Figure 1.**
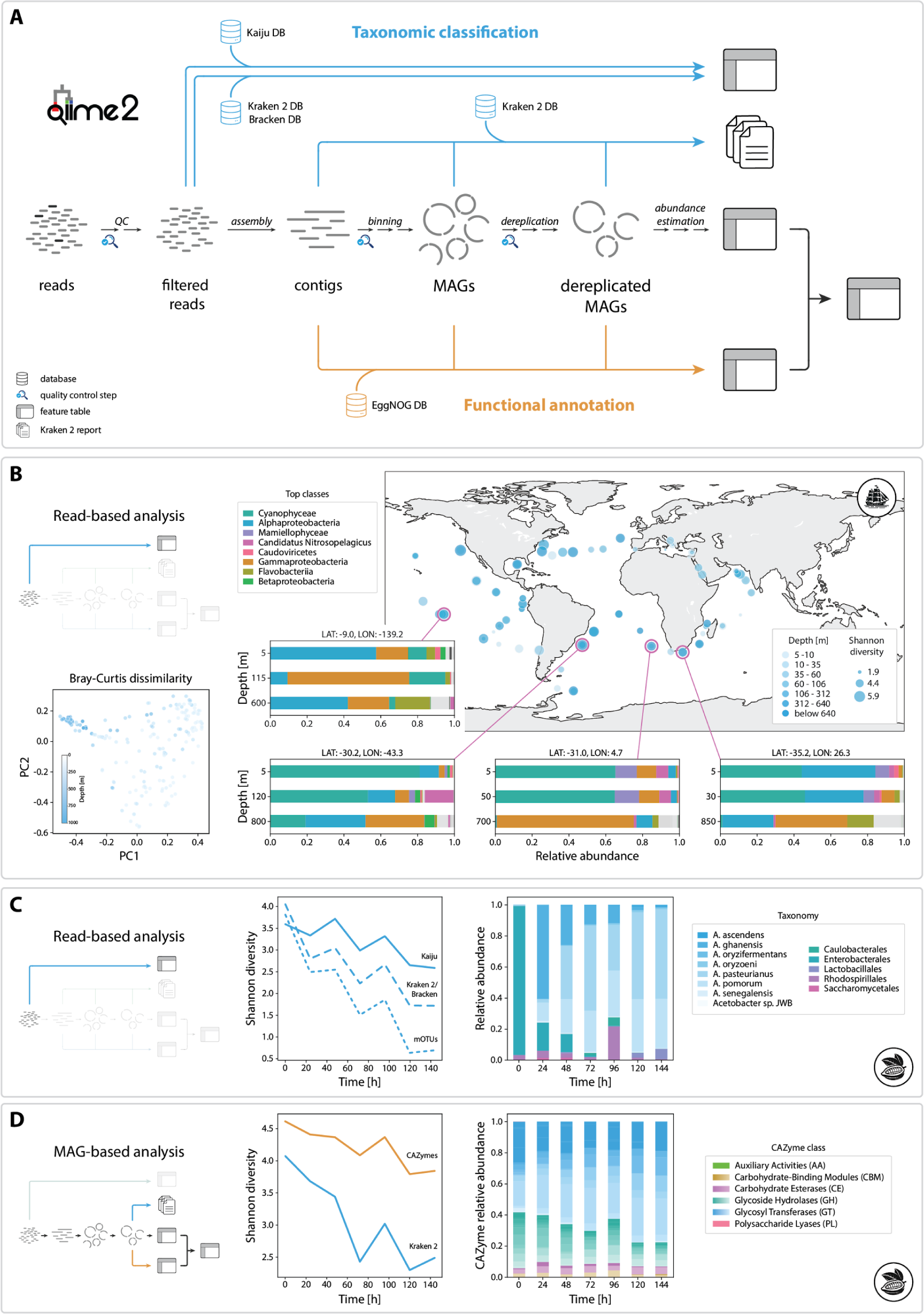
Overview of MOSHPIT and demonstration analysis. (A) Schematic of current analysis workflows available in MOSHPIT. Taxonomic annotation with Kaiju is supported for raw reads, and Kraken 2 can be applied to classify raw reads, contigs, or dereplicated MAGs. Functional annotation with EggNOG-mapper is available for contigs or (dereplicated) MAGs. (B) Reanalysis of the TARA Oceans dataset. The map depicts Shannon diversity of globally collected samples, with zoomed-in views of four locations showing taxonomic assignments across sample depths. Bray-Curtis principal coordinate scatterplot highlights compositional similarities among deep-sea samples. (C-D) Read-based (C) and MAG-based analysis (D) of cocoa demonstrates a consistent diversity decline over the fermentation process, accompanied by shifts in functional gene profile.

MOSHPIT includes built-in support for retrieving diverse pre-built databases as well as constructing custom databases, e.g., for Kraken2/Bracken, Kaiju, Diamond and EggNOG. This integration streamlines workflows and enables precise versioning of reference databases used throughout the analysis, significantly enhancing reproducibility as database retrieval date, query parameters, and version information are embedded in provenance information.

Beyond integration of existing tools, we have introduced custom actions to enhance MAG analysis, including: MAG dereplication with q2-sourmash^16^ (https://github.com/dib-lab/q2-sourmash) to produce a set of non-redundant genomes; estimating genome abundance through read mapping; integrating abundance data with functional annotation tables; and MAG quality filtering and human host read removal via mapping reads to the human pangenome^17^ and GRCh38 genome (Supplementary Figure 1). Customized host read filtering workflows, e.g., with other human or non-human reference genome collections, are also possible.

Each workflow in MOSHPIT incorporates checkpoint visualizations, allowing users to inspect results throughout their analysis. Notable examples include quality control of genome assembly (via the evaluate-contigs action incorporating metaQUAST^11^) and contig binning (via the evaluate-busco action incorporating BUSCO^12^), both accessible via a web browser interface for interactive exploration and quality inspection.

MOSHPIT leverages recent enhancements to the Q2F to maximize efficiency in the analysis process. The artifact cache eliminates repeated compression and decompression of large QIIME Zipped Artifacts (.qza files), conserving time and computational resources. Additionally, built-in parallelization support, powered by Parsl^18^, enables users to process multiple data partitions simultaneously on systems ranging from multi-core laptops to high-performance compute clusters. Integrated Pipeline resumption allows restarting interrupted or failed runs while reusing results that had already finished processing. Q2F’s Provenance Replay^19^ automates retrospective workflow documentation, facilitating transparency and replication.

To demonstrate MOSHPIT’s capability for handling large WMS datasets, we re-analyzed a subset of samples from the TARA Oceans Expedition^20^, specifically focusing on the archaea and bacteria. Leveraging MOSHPIT’s interoperability with existing QIIME 2 plugins, we used q2-fondue^21^ to retrieve the original reads from the NCBI Sequence Read Archive (SRA), followed by taxonomic classification using Kraken 2 within q2-annotate and downstream diversity analysis in Q2F (Fig. 1B).

To demonstrate an assembly-based workflow and highlight compatibility with other community-supported plugins, we re-analyzed 14 cocoa fermentations^22^. Results consistently recapitulate findings of the original study, via both read-based classification with three methods (Kraken 2, Kaiju, and mOTUs 3^23^, a community-developed plugin) (Fig. 1C) and MAG-based taxonomic classification and diversity analysis. Functional annotation of the recovered genomes in MOSHPIT yielded CAZyme gene counts for each sample (Fig. 1D), revealing a decrease in CAZyme diversity over the course of the fermentation process.

MOSHPIT provides a comprehensive platform built on the Q2F to seamlessly integrate marker-gene and metagenome analyses, bridging a critical gap in the microbiome research community. By incorporating state-of-the-art tools for read-based and assembly-based workflows alongside downstream tools for statistical analysis and visualization of diverse data types, MOSHPIT allows researchers to perform efficient, end-to-end analyses within a single, unified ecosystem, enabling flexible and fully reproducible integration of WMS analysis pipelines with the large variety of downstream analysis modules already available in QIIME 2. Using the newly developed plugin development tutorial in *Developing with QIIME 2* (https://develop.qiime2.org), the ecosystem is readily extensible. Leveraging Q2F’s built-in parallelization features, users can now scale their workflows with ease, optimizing resource use across desktop and high-performance computing environments alike. This expansion enhances accessibility, reproducibility, and scalability while providing best-in-class methods, capturing both taxonomic and functional dimensions of microbial communities for richer, more comprehensive community insights.

## Funding acknowledgement

This work was supported in part by grant 2022-309890 from the Chan Zuckerberg Initiative DAF, an advised fund of the Silicon Valley Community Foundation, to NAB and JGC; by NIH National Cancer Institute Informatics Technology for Cancer Research Award 1U24CA248454-01 to JGC; and by the Open Research Data Program of the ETH Board to NAB.

