## Supplementary Information for "MOSHPIT: accessible, reproducible metagenome data science on the QIIME 2 framework"

#### Analysis workflows

##### MAG reconstruction

MAG reconstruction workflow (Sup. Fig. 1) comprises all the steps which are required to recover Metagenome-Assembled Genomes from reads, including their dereplication into a non-redundant set of genomes. Initially, reads can be quality controlled/filtered to remove potential contaminants (e.g. host reads). On top of the *filter-reads* action (available through the existing q2-quality-control plugin), we developed the new *filter-reads-pangenome* action (as part of the q2-annotate plugin) which allows the users to automatically construct an index comprised of the GRCh38 reference human genome and the most recently published human pangenome[1] and use it to filter out human host reads.

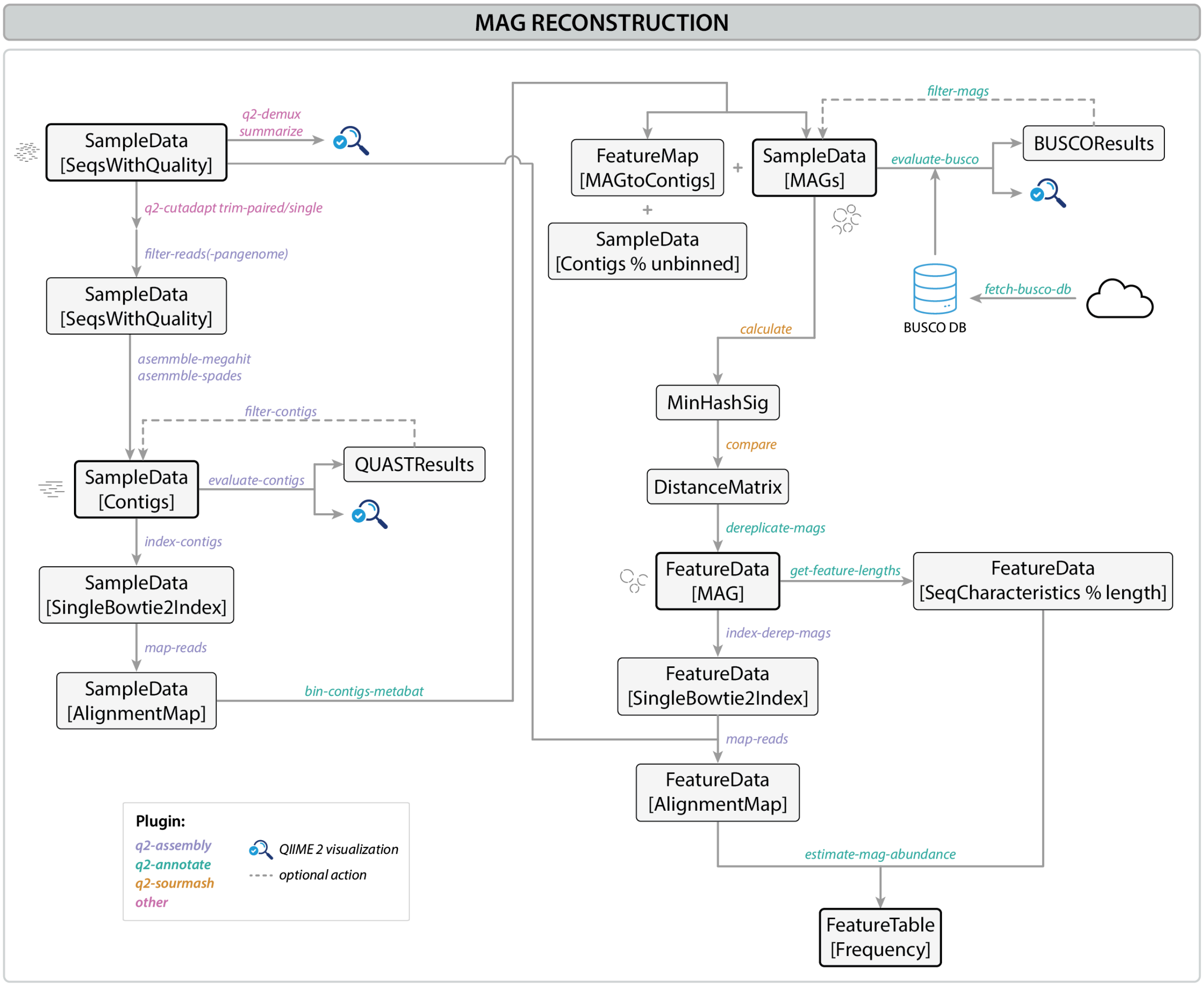

**Supplementary Figure 1.** MOSHPIT MAG reconstruction workflow.

Following the quality control steps, metagenome assembly can be performed using one of the two actions: *assemble-megahit* or *assemble-spades*, which will use the respective assembler (MEGAHIT[2] or SPAdes[3]) to reconstruct the contigs. Those can then be quality controlled using the *evaluate-contigs*[4] action and indexed with the *index-contigs* action. In preparation for the binning step, the original reads can then be mapped to the contigs using the index generated in the previous step (*map-reads* action) and used as an input to the *bin-contigs-metabat* action, which will use MetaBat 2[5] to form the bins. Those can then be quality controlled with the *evaluate-busco* action[6]. Following binning, construction of a non-redundant set of genomes may be performed by first generating a distance matrix for the given genome set and using it to cluster genomes together and select a representative one per cluster. Our workflow makes use of the per-MAG MinHash signatures generated through the *calculate* action (q2-sourmash plugin[7]), followed by the *compare* action (from the same plugin) to generate the distance matrix. The resulting table can then be input into our *dereplicate-mags* action which will find MAG clusters based on a distance threshold using hierarchical clustering with Ward linkage. It will then select the longest genome in each cluster to be its representative and return those non-redundant genomes together with a presence-absence table indicating which samples they were found in. Finally, to estimate the abundance of those genomes in the original samples, the dereplicated MAGs can be indexed using the *index-derep-mags* action (q2-assembly), followed by read mapping using the *map-reads* action. The resulting maps, together with genome lengths (obtained via the get-feature-lengths action), can be input into the *estimate-mag-abundance* action. This action will count the reads mapped to every MAG (while applying filtering parameters like minimal mapping quality, read length or base quality) and convert the counts using one of the available metrics like RPKM or TPM. Those metrics will then be returned as a feature table which can be used with any of the existing downstream QIIME 2 actions.

#### Taxonomic classification

Within the MOSHPIT plugin suite we provide two commonly used taxonomic classifiers (Sup. Fig. 2): Kraken 2[8] together with its Bracken companion[9] used for read abundance correction and Kaiju[10]. We include actions which will allow users to easily fetch any of the pre-built databases for both of those classifiers (*build-kraken-db* and *fetch-kaiju-db* actions) and build custom Kraken 2 databases (build-kraken-db). We are planning on including an action to build custom Kaiju databases at a later point. Kraken 2 can be used to classify reads, contigs and MAGs (before and after dereplication) by using the *classify-kraken2* action. The reports obtained for the input reads can be then used as an input to the *estimate-bracken* action which will generate a feature table and a corresponding taxonomy. Furthermore, taxonomy features may be retrieved for the results generated from dereplicated MAGs by using the *kraken2-to-mag-features* - those can later be used to analyze the feature table obtained through MAG abundance estimation described in the previous section. The Kaiju classifier can be used to classify reads directly using the *classify-kaiju* action which will produce a feature table and a corresponding taxonomy, similarly to how the *classify-kraken2* action does when reads are used as an input.

#### Functional annotation

The functional annotation workflow (Sup. Fig. 3) is provided mostly by the new actions included in the q2-annotate QIIME 2 plugin. The majority of them wrap different functionalities provided by the EggNOG-mapper[11] and allow a more intuitive approach to functional annotation use this widely used tool, depending on the user’s needs. We included a couple of action for retrieval/construction of all

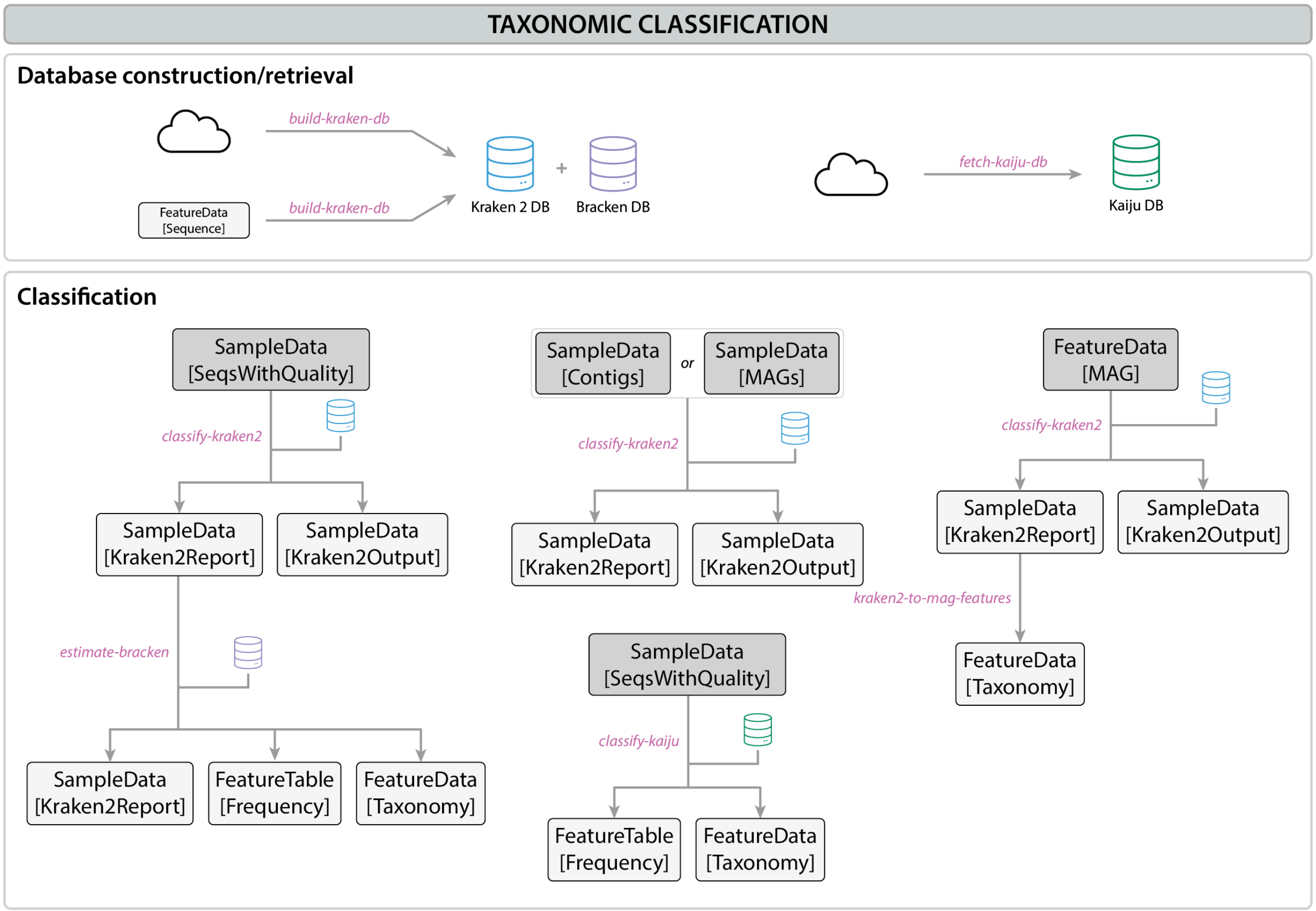

**Supplementary Figure 2.** MOSHPIT taxonomic classification workflow.

kinds of databases required throughout the functional annotation analysis: Diamond/HMMER databases for ortholog search as well as the EggNOG database used for ortholog annotation. Both, contigs and MAGs (before and after dereplication) can be first searched for protein orthologs using either *eggnog-diamond-search* or *eggnog-hmmer-search* actions. The resulting ortholog tables can then be annotated using the *eggnog-annotate* action and any specific annotation (e.g.: CAZymes or KEGG pathways) can be expanded into an independent feature table through the use of the *extract-annotations* action. This table can be then input into any other action accepting regular feature tables, e.g. to perform diversity analyses. Furthermore, when annotating the dereplicated MAGs (represented by the FeatureData[MAG] semantic type) this table can be combined with the feature table obtained from MAG abundance estimation step (see the “MAG reconstruction” workflow above) using the *multiply-tables* action. This will generate a feature table with annotation counts normalized by the MAG abundances in each sample.

#### MAGMock mock community analysis

To illustrate the new functionality of the MOSHPIT plugin suite we constructed three mock communities (further referred to as MAGMock) comprising seven different bacterial species with their reference genomes available in the RefSeq NCBI database: *Pseudomonas aeruginosa* PAO1, *Escherichia coli* str. K-12 substr. MG1655, *Escherichia coli* O157:H7 str. Sakai DNA, *Salmonella*

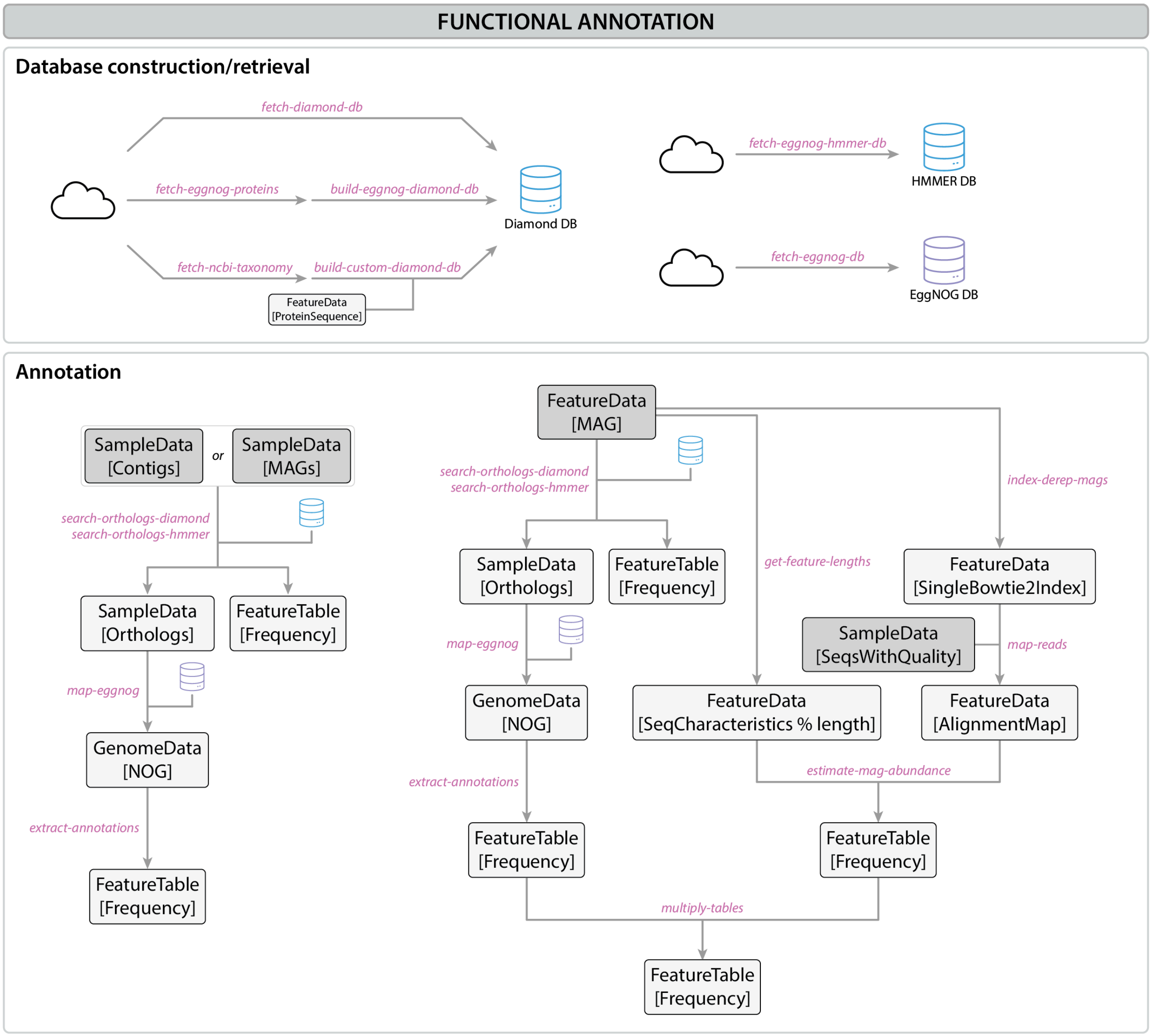

**Supplementary Figure 3.** MOSHPIT functional annotation workflow.

*enterica* subsp. *enterica* serovar Typhimurium str. LT2, *Staphylococcus aureus* subsp. *aureus* NCTC 8325, *Listeria monocytogenes* EGD-e, *Bacillus subtilis* subsp. subtilis str. 168 and *Mycobacterium tuberculosis* H37Rv. We used the Mason[12] read simulator to generate three samples with 20 million Illumina reads following three different abundance profiles: uniform, exponential and lognormal. All the samples underwent taxonomic classification using Kraken2, genome assembly using MEGAHIT, binning using MetaBAT 2 (followed by dereplication) and MAG abundance estimation. Moreover, we used Kraken 2 to classify the resulting dereplicated MAGs and performed functional annotation of those to illustrate the ability of our workflow to produce feature tables of selected gene annotations per sample. The fully re-runnable version of the Jupyter notebook containing the entire analysis can be found in the manuscript repository under https://github.com/bokulich-publications/moshpit-notebooks.

### Tara Oceans dataset analysis

To demonstrate the applicability of the MOSHPIT suite to analysis of large metagenomic datasets we chose to focus on a subset of samples from the TARA Oceans Expedition[13]. We selected the approx. 240 samples which correspond to the prokaryotic fraction of the ocean water. Moreover, we wanted to illustrate how the set of our new plugins can integrate with existing QIIME 2 plugins, including those developed by members of our community. We first pulled all the sequences from the SRA using q2-fondue[14] plugin developed in our lab and performed the taxonomic classification of the reads using three different classifiers: Kraken 2[8] and Kaiju[10] (both included in the MOSHPIT suite), as well as the community-developed mOTUs[15] QIIME 2 plugin. Inclusion of all those classifiers in the QIIME 2 ecosystem allowed us to easily compare and further analyze the TARA samples as feature tables received as a result of all the three actions are fully interoperable with the other QIIME 2 plugins.

#### Methods

##### Data retrieval and quality control

Sequencing data for 243 samples from the PRJEB1787 BioProject was fetched from the SRA archive using the q2-fondue plugin (https://github.com/bokulich-lab/q2-fondue) using the default parameters.

To ensure high-quality of read classification, reads were filtered using the q2-cutadapt plugin (**trim-paired** action) to remove all the reads shorter than 90 bp (*minimum_length*=90).

##### Read-based analysis

Read-based taxonomic classification was performed using the Kraken 2[8] tool built into the q2-annotate plugin. The Kraken 2 PlusPFP database was fetched using the **build-kraken-db** action and was used to classify the reads through the **classify-kraken2** action using the confidence score of 0.2 (while keeping all the other parameters at their default values). Following the classification, read counts were corrected for genome lengths using Bracken[9] through the **estimate-bracken** action (with *read_len*=100 and *threshold*=5 parameters) using the respective database fetched together with the Kraken 2 PlusPFP (see above). Subsequently, any unclassified reads or reads classified as *Homo sapiens* were removed from the resulting feature table by using the **filter-table** action from the feature-table plugin. Alpha- and beta-diversity metrics were calculated using the **core-metrics** action from the q2-diversity plugin using sampling depth of 617’000.

### Cocoa fermentation dataset analysis

To demonstrate both possible analysis pathways using our new MOSHPIT plugin suite we selected the cocoa fermentation dataset originally published by de Lima *et al.*[16]. The dataset comprises 14 time-resolved shotgun metagenomics samples obtained for two different varieties of cocoa beans (7 samples per variety) - in this study we focused on the Forasteiro variety. For the read-based analysis we chose to use two different classifiers: Kraken 2, which we integrated into our q2-annotate plugin, and mOTUs, which is a plugin developed by the QIIME 2 community and illustrates how our pipeline directly integrates with tools developed by other users. Using this approach we were able to reproduce the findings of the original study. Results obtained with both classifiers consistently showed the decrease in taxonomic diversity during the cocoa fermentation process for both seed varieties

(Fig. 1C). Moreover, we could detect the significant decrease in abundance of the Acetobacter and decrease of the Pantoea genera (Fig. 1C) as reported by the authors of the study.

Since the results of the original study were obtained using the read-based approach, we wanted to supplement this analysis by showcasing the assembly-based pathway of our pipeline. We assembled contigs and binned them into MAGs which were then subject to the dereplication procedure, as described in the methods section. We retained only MAGs which were flagged by BUSCO as at least 50% complete and annotated those with Kraken 2. The taxonomic diversity across the fermentation period was consistent with the results obtained from reads only - we saw a decrease in Shannon diversity for both of the seed types (Fig. 1D).

In addition to the taxonomic analysis, we wanted to explore the functional potential of the studied communities to demonstrate the potential of the MOSHPIT pipeline in performing functional annotation. We used the EggNOG tool, integrated into the q2-annotate plugin, to identify and annotate orthologs in the dereplicated set of MAGs. We were then able to extract the counts of CAZymes present in each MAG and generate a feature table containing those counts normalized per sample. We observed a decrease in the Shannon diversity of CAZymes during the fermentation process, determined by an increase in abundance of CAZymes from the glycosyltransferases (GT) class and decrease of enzymes classified as glycoside hydrolases (GH; Fig. 1D).

#### Methods

##### Data retrieval

Sequencing data for all the samples from the BioProject PRJNA552479 was fetched from the SRA archive using the q2-fondue plugin (https://github.com/bokulich-lab/q2-fondue) using the default parameters.

##### Read-based analysis

Read-based taxonomic classification was performed using two different classifiers available through QIIME 2 plugins. Kraken 2 classifier is available as part of the MOSHPIT suite (within the q2-annotate plugin; https://github.com/bokulich-lab/q2-annotate) and mOTUs is available as a community plugin (https://github.com/motu-tool/q2-mOTUs).

The Kraken 2 PlusPF database was fetched using the **build-kraken-db** action from the q2-annotate plugin. It was then used to classify the reads through the **classify-kraken2** action using the confidence score of 0.5 (while keeping all the other parameters at their default values). Following the classification, read counts were corrected for genome lengths using Bracken through the **estimate-bracken** action (with *read_len*=150 and *threshold*=5 parameters) using the respective database fetched together with the Kraken 2 PlusPF (see above).

The mOTUs classification was performed using the default parameters through the **profile** action from the mOTUs plugin.

The nr_euk Kaiju database was fetched using the **fetch-kaiju-db** action from the q2-annotate plugin and used to classify the reads with the **classify-kaiju** action (with *c*=0.1).

Unclassified reads were removed from the resulting feature tables by using the **filter-table** action from the feature-table plugin. Alpha- and beta-diversity metrics were calculated using the **core-metrics** action from the q2-diversity plugin using sampling depths of 2’320’000 (for Kraken 2/Bracken), 5’298’000 (for Kaiju) or 1’792 (for mOTUs).

##### Assembly-based analysis

Reads were assembled into contigs using the MEGAHIT[2] assembler through the **assemble-megahit** action from the q2-assembly plugin with the “meta-sensitive” preset. The quality of the generated assemblies was checked with metaQUAST[4] through the **evaluate-contigs** action. The contigs were indexed with Bowtie 2[17] (**index-contigs** action) and the reads were mapped to contigs using those indices as input to the **map-reads** action.

Contig binning was performed with MetaBAT 2[5] available through the **bin-contigs-metabat** action from the q2-annotate plugin. The resulting MAGs were subject to quality control with BUSCO[6]. The database of prokaryotic BUSCOs was fetched using the **fetch-busco-db** action and used as input (together with the generated MAGs) to the **evaluate-busco** action.

The resulting QC table was used to filter the MAGs based on the completeness score - only MAGs which were at least 50% complete were retained (**filter-mags** action from q2-annotate with *on*=mag and *where*=”complete>50” parameters).

To generate a non-redundant set of MAGs we developed a dereplication action (**dereplicate-mags**) which can use a distance matrix of the original MAGs, cluster similar MAGs given a similarity threshold and identify the single MAG representative of each cluster (based on the assembly length). The distance matrix was obtained using the SourMash[7] tool: MinHash signatures of each MAG were calculated using the **compute** action from the sourmash plugin (using *ksizes*=35 and *scaled*=10 parameters) and compared using the **compare** action from the same plugin. The resulting distance matrix was then used as input to the **dereplicate-mags** action which generated a non-redundant set of MAGs at the similarity threshold of 0.99.

The taxonomic identity of the dereplicated MAGs was determined using the **classify-kraken2** action as described in the section above. The taxonomic assignment of each MAG was then obtained through the **kraken2-to-mag-features** action which performs an LCA selection from all the classified contigs within each individual MAG.

To obtain the abundances of each MAG in the original community we developed a set of actions which map the original reads to the dereplicated MAGs and use read counts (expressed as RPKM or TPM values) as a proxy for abundance. We first indexed the dereplicated MAGs using the **index-derep-mags** action (q2-assembly), followed by mapping the reads to the generated indices using the **map-reads** action. The obtained alignment maps were then used as input to the **estimate-mag-abundance** action (together with MAG lengths extracted using the **get-feature-lengths** action). In order to only retain reads which mapped with high quality we set the *min_mapq* parameter to 42 and *min_base_quality* to 20. This action uses the *samtools coverage* tool to extract coverage of each MAG and uses the extracted counts to estimate the RPKM and TPM values.

The functional profile of each community was obtained using the EggNOG[11] annotation tool which we incorporated into the q2-annotate plugin. The required databases were retrieved using the

**fetch-diamond-db** and **fetch-eggnog-db** actions. The ortholog candidates were identified using the **search-orthologs-diamond** action using the above-mentioned database and the dereplicated set of MAGs. The resulting ortholog tables were then input to the **map-eggnog** action (together with the corresponding database) which generated tables of EggNOG annotations. We used the **extract-annotations** action to retrieve the CAZyme annotations from the tables obtained from EggNOG and convert them into a feature table (using *annotation*=caz and *max_evalue*=0.0001 parameters). This feature table was then input to the **multiply-tables** action (together with the MAG abundance feature table) to calculate their dot product representing abundances of each CAZyme in each sample.

#### Distribution and installation

The MOSHPIT software suite is provided as a self-contained QIIME 2 distribution named "metagenome", in the form of fully resolved conda environments and docker images. All the updates to any of the MOSHPIT plugins will follow the regular QIIME 2 release cycle and will be announced on the QIIME 2 forum (https://forum.qiime2.org). All the information about the exact plugin composition and installation instructions for all the supported operating systems can be found in the official QIIME 2 documentation on https://docs.qiime2.org.

#### MOSHPIT tutorials

We have developed a set of tutorials demonstrating all the functionality described in this manuscript using the cocoa fermentation dataset. The tutorials are divided into workflow-based sections and can be found under https://moshpit.readthedocs.io.

#### MOSHPIT Nextflow workflow

To enable end-to-end analyses of whole metagenome data using MOSHPIT we have also been actively working on a Nextflow workflow which allows deployment of all of the described functionality on a variety of clusters and computing environments using both, conda environments and containers. The workflow can be found under https://github.com/bokulich-lab/moshpit-nf.

### MOSHPIT tutorials

The Bokulich Lab

Jan 27, 2025

MOSHPIT (MOdular SHotgun metagenome Pipelines with Integrated provenance Tracking) is a toolkit of plugins for whole metagenome assembly, annotation, and analysis built on the microbiome multi-omics data science framework QIIME 2. MOSHPIT enables flexible, modular, fully reproducible workflows for read-based or assembly-based analysis of metagenome data.

The following main plugins comprise the core of the MOSHPIT toolkit:

- q2-assembly: contains actions for (meta)genome and quality control, genome indexing and red mapping
- q2-annotate: provides actions for contig binning and quality control, taxonomic and functional annotations of contigs and MAGs, human host removal.

Additionally, you may want to check out these other QIIME 2 plugins for antimicrobial resistance gene (ARG) detection and viromics applications. These plugins are not covered in this tutorial. They have their own installation instructions and tutorials (see the wiki page on the respective GitHub repositories). You can use these plugins with some of the artifacts produced by q2-assembly and q2-annotate:

- q2-rgi: antimicrobial resistance gene annotation of MAGs and metagenomic reads with RGI and CARD
- q2-amrfinderplus: ARG detection using the AMRFinderPlus tool
- q2-viromics: detection of viral sequences and their quality control.

This tutorial will guide you through the process of analyzing metagenomic data using QIIME 2 framework and MOSHPIT. The tutorial is divided into several chapters, each focusing on a different aspect of metagenomic data analysis. We will use a small published dataset to demonstrate the capabilities of most of the methods available in MOSHPIT.

We will begin by setting up our computational environment and fetching all the necessary data (see *Setup* and *Data retrieval*). Then, we will move to quality control and filtering of the raw reads (see *Quality control*). Once we have our clean dataset, we can start by recovering metagenome-assembled genomes (MAGs) (see *here*), followed by taxonomic classification of reads and MAGs themselves (see *Taxonomic classification*). Finally, we will estimate perform functional annotation of the dereplicated MAGs (see *Functional annotation*).

Let's dive in!

CHAPTER

ONE

### SETUP

Before we dive into the tutorial, let's make sure we have all the necessary components in place. Make sure you have a working MOSHPIT environment available - please follow the instructions from the developer QIIME 2 documentation to install all the required components. These installation instructions will also become available through the official QIIME 2 user documentation in the 2025.4 release later this year.

In this tutorial we will be storing all the data in the QIIME 2 cache. To learn more about how the cache works, you can refer to the *Using an Artifact Cache* tutorial. (Note that all calls to `qiime` in that document can be replaced by calls to `mosh`.) You should create a single cache directory in the current working directory by running the following command:

```
mosh tools cache-create --cache ./cache
```

#### 1.1 Note on parallelization

While we do not explicitly mention parallelization in the tutorial, many of the QIIME 2 actions can be parallelized by executing actions on smaller subsets of the data called **partitions**. To make use of parallelization, you will need a parsl config file which will define the resources available to the parallel execution. Here is an example of a config file which was used to assemble contigs through the `assemble-megahit` action:

```
[parsl]

[[parsl.executors]]
class = "HighThroughputExecutor"
label = "default"
max_workers = 1

[parsl.executors.provider]
class = "SlurmProvider"
scheduler_options = "#SBATCH --mem-per-cpu=4G"
exclusive = false
worker_init = "source ~/.bashrc && conda activate q2-metagenome-2024.5"
walltime = "24:00:00"
nodes_per_block = 1
cores_per_node = 24
max_blocks = 14
```

To learn more about how to configure parallelization in QIIME 2, please consult the official documentation.

**Note**

To examine your generated QIIME 2 visualizations, you can use *QIIME 2 View*.

CHAPTER

TWO

### DATA RETRIEVAL

The dataset used in this tutorial is available through the NCBI Sequence Read Archive (SRA). To retrieve it we will use the q2-fondue plugin for programmatic access to sequences and metadata from SRA; we only need to provide a list of accession IDs to download - q2-fondue will take care of the rest.

> **Note**
>
> You need to provide an e-mail address when running this command - this is required by the NCBI as a way to ensure they can contact you in case of any issues.

- download the files containing all the accession IDs and corresponding metadata:

```
wget -O ./ids.tsv https://raw.githubusercontent.com/bokulich-lab/moshpit-docs/main/
↪moshpit_docs/data/ids.tsv
```

```
wget -O ./metadata.tsv https://raw.githubusercontent.com/bokulich-lab/moshpit-docs/
↪main/moshpit_docs/data/metadata.tsv
```

- import the file into a QIIME 2 artifact:

```
mosh tools cache-import \
    --type 'NCBIAccessionIDs' \
    --input-path ./ids.tsv \
    --cache ./cache \
    --key ids
```

- run the `get-all` action from the `fondue` plugin:

```
mosh fondue get-all \
    --i-accession-ids ./cache:ids \
    --p- \
    --p-n-jobs 5 \
    --p-retries 5 \
    --o-paired-reads ./cache:reads_paired \
    --o-metadata ./cache:metadata \
    --o-single-reads ./cache:reads_single \
    --o-failed-runs ./cache:failed_runs \
    --verbose
```

CHAPTER

THREE

### QUALITY CONTROL

As with any other NGS experiment, metagenome data should be quality controlled before any downstream analysis. The filtering steps may include adapter removal, quality trimming, and filtering out low-quality reads. Moreover, depending on the sample type and preparation procedures, metagenomic reads may contain host DNA, which should be removed. Other QIIME 2 plugins already provide generalized functionality to address quality filtering/control of next-generation sequencing data - the MOSHPIT plugin suite expands on these by focusing more on host DNA removal from metagenome data. The next sections contain a brief overview of the filtering steps which can be done using QIIME 2 and MOSHPIT.

#### 3.1 Quality filtering

##### 3.1.1 Quality overview

We can get an overview of the read quality by using the `summarize` action from the `demux` QIIME 2 plugin. This command will generate a visualization of the quality scores at each position. You can learn more about this action in the QIIME 2 documentation.

```
mosh demux summarize \
    --i-data ./cache:reads \
    --o-visualization demux.qzv
```

To see an example of the visualization you can go here.

##### 3.1.2 Read trimming and quality filtering

In order to remove low quality bases from the reads, we can use one of the `trim` actions from the `cutadapt` QIIME 2 plugin. Here we are using the `trim-paired` action to remove all the reads shorter than 90 bp:

```
mosh cutadapt trim-paired \
    --i-demultiplexed-sequences ./cache:reads \
    --p-minimum-length 90 \
    --o-trimmed-sequences ./cache:reads_trimmed
```

#### 3.2 Host read removal

There are a few different options to perform host read removal in QIIME 2: a more generic one using the `filter-reads` action and a more specific one using the `filter-reads-pangenome` action. Below you can learn how to use both of them. In this tutorial we will use the `filter-reads-pangenome` action to remove human reads from the dataset.

##### 3.2.1 Removal of contaminating reads

Removal of contaminating reads can generally be done by mapping the reads to a reference database and filtering out the reads that map to it. In QIIME 2 this can be done by using the `filter-reads` action from the `quality-control` plugin. Before filtering we need to construct the index of the reference database that will be used by Bowtie 2:

- start with the FASTA files containing the reference sequences - we will import them into a QIIME 2 artifact:

```
mosh tools cache-import \
    --cache ./cache \
    --key reference_seqs \
    --type "FeatureData[Sequence]" \
    --input-path ./reference_seeqs.fasta
```

- build the Bowtie 2 index:

```
mosh quality-control bowtie2-build \
    --i-sequences ./cache:reference_seqs \
    --o-database ./cache:reference_index
```

- filter out the reads that map to the reference database:

```
mosh quality-control filter-reads \
    --i-demultiplexed-sequences ./cache:reads_trimmed \
    --i-database ./cache:reference_index \
    --o-filtered-sequences ./cache:reads_filtered
```

##### 3.2.2 Human host reads

Contaminating human reads can also be filtered out using the approach shown above by providing a human reference genome. Since a single human reference genome is not enough to cover all the human genetic diversity, it is recommended to use a collection of genomes represented by the human pangenome (__CIT). We have built a new QIIME 2 action `filter-reads-pangenome` which allows to first fetch the human pangenome sequence, combine it with the GRCh38 reference genome, build a combined Bowtie 2 index and, finally, filter the reads against it. Next to the filtered reads, the action will also return the generated index so that it can be used in any other experiments.

```
mosh moshpit filter-reads-pangenome \
    --i-reads ./cache:reads_trimmed \
    --o-filtered-reads ./cache:reads_filtered \
    --o-reference-index ./cache:human_reference_index
```

### CHAPTER FOUR

### RECOVERY OF METAGENOME-ASSEMBLED GENOMES

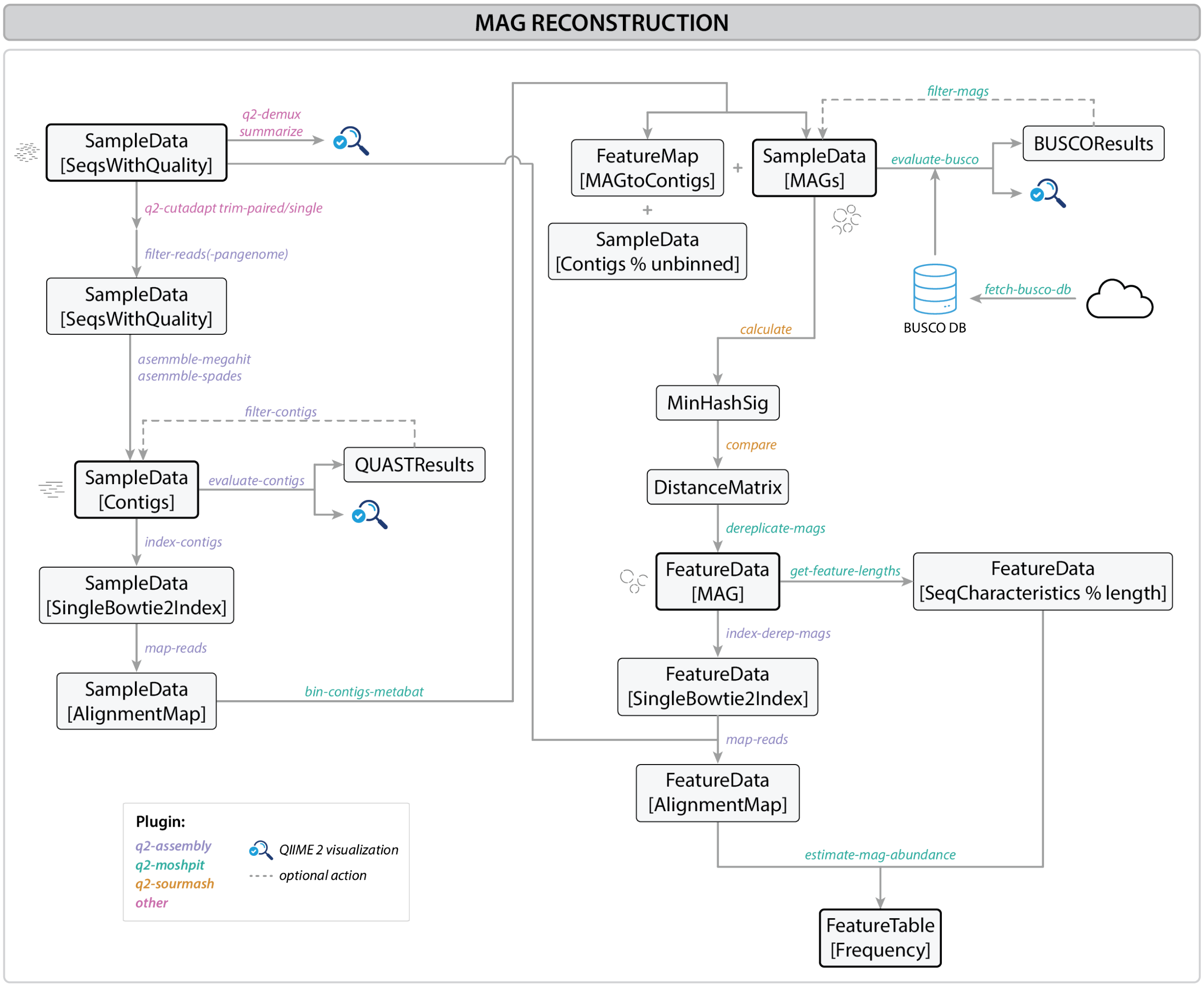

Fig. 4.1: MAG reconstruction workflow

**Metagenome-assembled genomes (MAGs)** are genomes reconstructed directly from DNA from complex microbial mixtures without the need for culturing organisms in the laboratory. This approach allows researchers to explore the genetic makeup of microbial communities in various environments, providing insights into the diversity, functions, and ecological roles of uncultured microorganisms. Recovering MAGs involves assembling sequencing reads into contigs, binning contigs into draft genomes, and evaluating their quality. Optionally, MAGs can be further dereplicated into a set of non-redundant genomes.

This workflow describes a step-by-step process for MAGs recovery using QIIME2. Each command includes explanations of the parameters used.

**For more information on the tools used in this workflow, refer to their official documentation:**

- MEGAHIT: https://github.com/voutcn/megahit
- SPAdes: https://github.com/ablab/spades
- QUAST: https://github.com/ablab/quast
- MetaBAT: https://bitbucket.org/berkeleylab/metabat/
- BUSCO: https://gitlab.com/ezlab/busco
- Sourmash: https://github.com/dib-lab/sourmash
- Kraken2: https://github.com/DerrickWood/kraken2
- QIIME 2: https://github.com/qiime2

#### 4.1 Recovery of MAGs

In this part of the tutorial we will go thorugh the steps required to recover metagenome-assembled genomes (MAGs) from metagenomic data. The workflow is divided into several steps, from contig assembly to binning and quality control.

**Warning**

Genome assembly and contig binning can be highly resource-intensive. Ensure that your system has sufficient CPU and memory resources before running these commands.

##### 4.1.1 Assemble contigs with MEGAHIT

The first step in recovering MAGs is genome assembly itself. There are many genome assemblers available, two of which you can use through the q2-assembly plugin - here, we will use MEGAHIT. MEGAHIT takes short DNA sequencing reads, constructs a simplified De Bruijn graph, and generates longer contiguous sequences called contigs, providing valuable genetic information for the next steps of our analysis.

- The `--p-presets` specifies the preset mode for MEGAHIT. In this case, it's set to "meta-sensitive" for metagenomic data.
- The `--p-cpu-threads` specifies the number of CPU threads to use during assembly.
- The `--p-min-contig-len` specifies the minimum length of contigs to keep.

**Warning**

`--p-coassemble` parameter can be set to "True" if you wish to co-assemble reads into contigs from all samples. **This parameter is still under development**: you will not be able to use the generated contigs for further analysis.

```
mosh assembly assemble-megahit \
    --i-seqs ./cache:reads_filtered \
    --p-presets "meta-sensitive" \
    --p-num-cpu-threads 24 \
    --p-min-contig-len 200 \
```

(continues on next page)

(continued from previous page)

```
    --p-coassemble False \     # Co-assembly is disabled for this example
    --o-contigs ./cache:contigs \
    --verbose
```

- Alternatively, you can also use `mosh assembly assemble-spades` to assemble contigs with SPAdes.

##### 4.1.2 Contig QC with QUAST

Once the reads are assembled into contigs, we can use QUAST to evaluate the quality of our assembly. There are many metrics that can be used for that purpose but here we will focus on the two most popular metrics:

- **N50**: represents the contiguity of a genome assembly. It's defined as the length of the contig (or scaffold) at which 50% of the entire genome is covered by contigs of that length or longer - the higher this number, the better.
- **L50**: represents the number of contigs required to cover 50% of the genome's total length - the smaller this number, the better.

In addition to calculating generic statistics like N50 and L50, QUAST will try to identify potential genomes from which the analyzed contigs originated. Alternatively, we can provide it with a set of reference genomes we would like it to run the analysis against using `--i-references`.

```
mosh assembly evaluate-contigs \
    --i-contigs ./cache:contigs  \
    --p-threads 128 \
    --p-memory-efficient \
    --o-visualization ./results/contigs.qzv \
    --verbose
```

Your visualization should look similar to this one.

##### 4.1.3 Index contigs

In this step, we generate an index for the assembled contigs. This index is required for mapping reads to the contigs later. Various parameters control the size and structure of the index, as well as resource usage.

```
mosh assembly index-contigs \
    --i-contigs ./cache:contigs \
    --p-threads 8 \
    --o-index ./cache:contigs_index \
    --verbose
```

##### 4.1.4 Map reads to contigs

Here we map the input paired-end reads to the indexed contigs created in the previous step. We use various alignment settings to ensure optimal mapping, including local alignment mode and sensitivity settings.

```
mosh assembly map-reads \
    --i-index ./cache:contigs_index \
    --i-reads ./cache:reads_filtered \
    --o-alignment-map ./cache:reads_to_contigs \
    --verbose
```

##### 4.1.5 Bin contigs with MetaBAT

Binning contigs involves grouping assembled contigs into MAGs. This step uses MetaBAT to assign contigs based on co-abundance and other features, producing MAG files that represent putative genomes.

```
mosh moshpit bin-contigs-metabat \
    --i-contigs ./cache:contigs \
    --i-alignment-maps ./cache:reads_to_contigs \
    --p-num-threads 64 \
    --p-seed 100 \
    --p-verbose \
    --o-mags ./cache:mags \
    --o-contig-map ./cache:contig_map \
    --o-unbinned-contigs ./cache:unbinned_contigs \
    --verbose
```

This step generated several artifacts:

- `mags`: these are our actual MAGS, per sample.
- `contig_map`: this is a mapping between MAG IDs and IDs of contigs that belong to a given MAG.
- `unbinned_contigs`: these are all the contigs that could not be assign to any particular MAG. From now on, we will focus on the `mags`.

##### 4.1.6 Evaluate bins with BUSCO

This step evaluates the completeness and quality of MAGs using the BUSCO tool, which checks for the presence of single-copy orthologs. The evaluation helps ensure the quality of the recovered MAGs.

First we will use `mosh moshpit fetch-busco-db` to download a specific lineage's BUSCO database. BUSCO databases are precompiled collections of orthologous genes, tailored to specific lineages such as viruses, prokaryotes (bacteria and archaea), or eukaryotes.

- The `--p-prok` True parameter specifies that we want to download the prokaryote dataset (for bacterial genomes, for example).

```
mosh moshpit fetch-busco-db \
    --p-prok True \
    --o-busco-db ./cache:busco_db
    --verbose
```

Once the appropriate BUSCO database is fetched, the next step is to evaluate the completeness and quality of the MAGs.

```
mosh moshpit evaluate-busco \
    --i-bins ./cache:mags \
    --i-busco-db ./cache:busco_db \
    --p-lineage-dataset bacteria_odb10 \
    --p-cpu 16 \
    --o-visualization ./results/mags.qzv \
    --o-results-table ./cache:busco_results \
    --verbose
```

The `--p-lineage-dataset bacteria_odb10` parameter specifies the particular lineage dataset to use, in this case, the bacteria_odb10 dataset. This is a standard database for bacterial genomes.

Your visualization should look similar to this one.

##### 4.1.7 Filter MAGs

This step filters MAGs based on completeness. In this example, we filter out any MAGs with completeness below 50%. The filtering process ensures only high-quality genomes are kept for downstream analysis.

**Tip**

We recommed that this step is done before dereplication (as in this example). Alternatively, we can also use the *dereplicated set* and filter this one using `qiime moshpit filter-derep-mags`.

```
mosh moshpit filter-mags \
    --i-mags ./cache:mags \
    --m-metadata-file ./cache:busco_results \
    --p-where 'complete>50' \
    --p-no-exclude-ids \
    --p-on mag \
    --o-filtered-mags ./cache:mags_filtered_50 \
    --verbose
```

#### 4.2 MAG set dereplication

Depending on the application, it may be necessary to dereplicate the set of MAGs to remove redundancy and retain only unique genome representatives. Our workflow includes a dereplication step that can use any genome distance matrix to find clusters of similar genomes (based on a specific similarity threshold) and identify the most representative MAG (in our case, it will be the longest genome in the cluster). Here we use Sourmash to generate the distance matrix but any other tool could also be used.

##### 4.2.1 Compute MinHash signatures with Sourmash

In this step, Sourmash is used to compute MinHash signatures for the filtered MAGs. MinHash is a method used to compare large datasets efficiently by creating compressed representations of genomes.

```
mosh sourmash compute \
    --i-sequence-file ./cache:mags_filtered_50 \
    --p-ksizes 35 \
    --p-scaled 10 \
    --o-min-hash-signature ./cache:mags_minhash_50 \
    --verbose
```

##### 4.2.2 Compare MinHash signatures

Here we compare the computed MinHash signatures to evaluate the similarity between the genomes. This step will allow for dereplication by identifying highly similar genomes.

```
mosh sourmash compare \
    --i-min-hash-signature ./cache:mags_minhash_50 \
    --p-ksize 35 \
    --o-compare-output ./cache:mags_dist_matrix_50 \
    --verbose
```

##### 4.2.3 Dereplicate MAGs

This step dereplicates the filtered MAGs, ensuring that only unique MAGs are retained. Dereplication reduces redundancy by merging similar genomes based on a similarity threshold.

```
mosh moshpit dereplicate-mags \
    --i-mags ./cache:mags_filtered_50 \
    --i-distance-matrix ./cache:mags_dist_matrix_50 \
    --p-threshold 0.99 \
    --o-dereplicated-mags ./cache:mags_derep_50 \
    --o-feature-table ./cache:mags_ft_50  \
    --verbose
```

#### 4.3 MAG abundance estimation

Once we recover MAGs from metagenomic data, we may be interested in estimating their abundance in the samples. We can do it by mapping the original reads to the dereplicated MAGs and calculating the abundance based on the read mapping results. There are a couple of ways to estimate MAG abundance, such as RPKM (Reads Per Kilobase per Million mapped reads) and TPM (Transcripts Per Million). Here we will use TPM to estimate the abundance of each MAG in all samples.

##### 4.3.1 Get MAG lengths

This step calculates the lengths of each dereplicated MAG, which will be used in the next step to estimate abundance.

```
mosh moshpit get-feature-lengths \
    --i-features ./cache:mags_derep \
    --o-lengths ./cache:mags_derep_length \
    --verbose
```

##### 4.3.2 Index dereplicated MAGs

This step indexes the dereplicated MAGs for read mapping. The index is necessary to efficiently map the input reads back to the MAGs.

```
mosh assembly index-derep-mags \
    --i-mags ./cache:mags_derep \
    --p-threads 8 \
    --p-seed 100 \
    --o-index ./cache:mags_derep_index \
    --verbose
```

##### 4.3.3 Map reads to dereplicated MAGs

In this step, we map the input paired-end reads back to the dereplicated MAGs. This helps in calculating the abundance of each MAG in the sample.

```
mosh assembly map-reads \
    --i-index ./cache:mags_derep_index \
    --i-reads ./cache:reads_filtered \
    --p-threads 8 \
    --p-seed 100 \
    --o-alignment-map ./cache:reads_to_derep_mags \
    --verbose
```

##### 4.3.4 Estimate MAG abundance

This step estimates the abundance of each MAG in the sample based on the read mapping results.

- `metric` : currently, we support RPKM and TPM
- `min-mapq` : indicates the minimum required read mapping quality - for Bowtie2, 42 will allow only perfect matches to be retained – `min-base-quality` : only keep alignments with this minimal Phred quality score

For more options, see –help.

```
mosh moshpit estimate-mag-abundance \
    --i-mag-lengths ./cache:mags_derep_length \
    --i-maps ./cache:reads_to_derep_mags \
    --p-threads 10 \
    --p-metric tpm \
    --p-min-mapq 42 \
    --o-abundances ./cache:mags_derep_ft \
    --verbose
```

##### 4.3.5 Let's have a look at our estimated MAG abundance!

First we will use Kraken 2 to classify provided MAGs into taxonomic groups.

**Note**

Refer to *Taxonomic classification of reads* section for more details on taxonomic classification with Kraken 2.

The database used here is the `PlusPF` database, defined here.

```
mosh moshpit classify-kraken2 \
    --i-seqs ./cache:mags_derep \
    --i-kraken2-db ./cache:kraken2_db \
    --p-threads 40 \
    --p-confidence 0.5 \
    --p-report-minimizer-data \
    --o-reports ./cache:kraken_reports_mags_derep \
    --o-hits ./cache:kraken_hits_mags_derep \
    --verbose
```

Then we will convert a Kraken 2 report into a generic taxonomy artifact for downstream analyses.

```
mosh moshpit kraken2-to-mag-features \
    --i-reports ./cache:kraken_reports_mags_derep  \
    --i-hits ./cache:kraken_hits_mags_derep  \
    --o-taxonomy ./cache:mags_derep_taxonomy \
    --verbose
```

Now we are ready to generate a taxa bar plot.

```
mosh taxa barplot \
    --i-table ./cache:mags_derep_ft \
    --i-taxonomy ./cache:mags_derep_taxonomy \
    --m-metadata-file ./cocoa-metadata.tsv \
    --o-visualization ./results/mags-derep-taxa-bar-plot.qzv \
    --verbose
```

Your visualization should look similar to this one.

CHAPTER

FIVE

### TAXONOMIC CLASSIFICATION

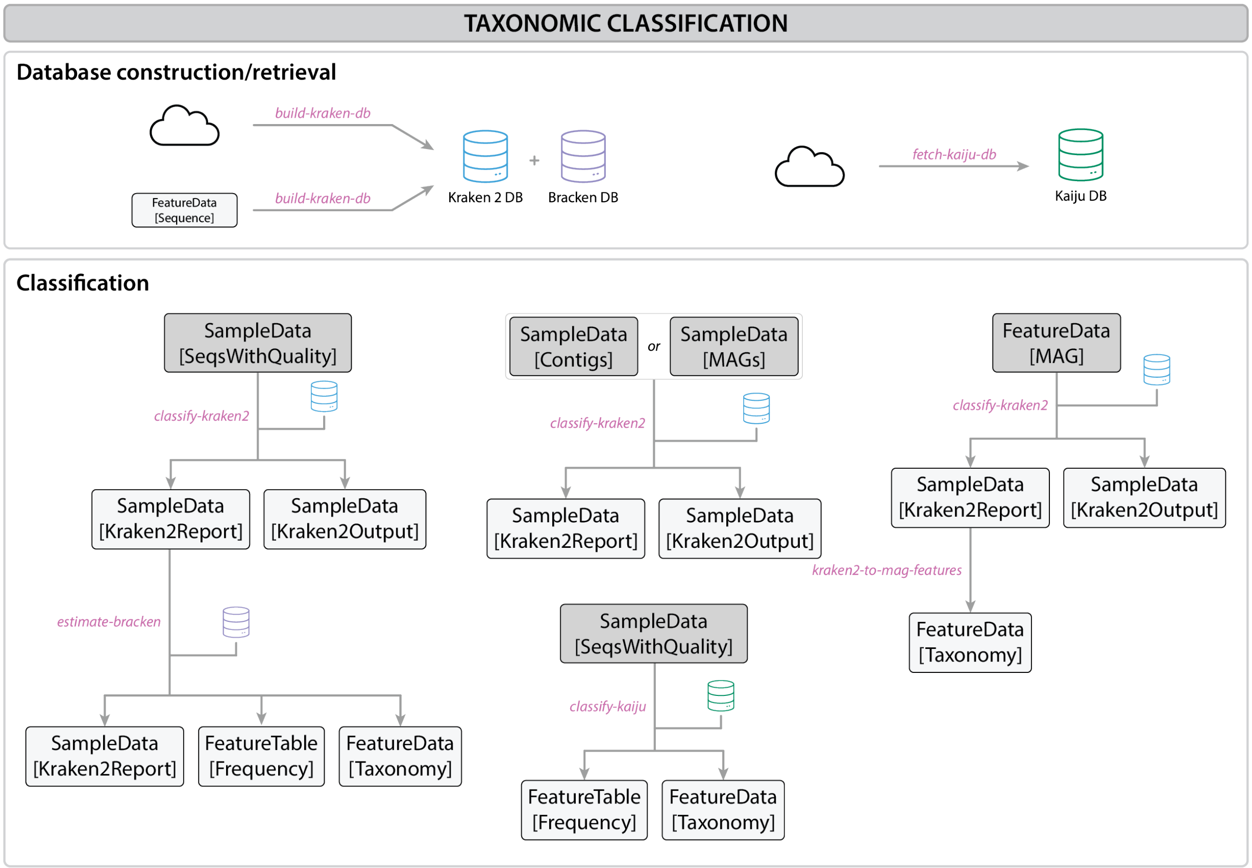

Fig. 5.1: Taxonomic classification workflow

#### 5.1 Read-based classification overview

Read-based classification is commonly used to determine the taxonomic groups present within a given sample. This technique is useful for assessing the biodiversity or the composition of microbial communities by assigning DNA reads to known organisms. One significant advantage of read-based classification is that it allows for the classification of all reads, including those that may not be directly involved in downstream analyses (e.g. assemblies or MAGs).

**Key Factors Influencing Read Classification:** The outcome of read classification is heavily influenced by the selection of reference databases. These databases vary in size, quality, and scope, which means that the more comprehensive and accurate the reference database is, the more accurate the classification of reads will be. Some databases might be specific to certain taxonomic groups, while others could provide a broader reference, potentially affecting the results depending on the sample type and research goals.

**See also**

For more information and a benchmark, consult Ye et al., 2019.

#### 5.2 Kraken 2: DNA-to-DNA classification

Kraken 2 is a DNA-to-DNA classification tool that assigns taxonomic labels to reads by directly comparing k-mers (short DNA sequence fragments of a fixed length, typically 31 base pairs) from the query read to a database of known sequences. Kraken 2 classifies the read based on the majority of k-mer matches within the read, providing fast and accurate taxonomic classification.

**See also**

For more information on Kraken 2, consult Wood et al., 2019.

#### 5.3 Kaiju: protein-based classification

Kaiju compares reads by translating DNA sequences into protein sequences (similar to BLASTx). This allows Kaiju to identify organisms accurately when nucleotide sequences are too divergent to be identified with DNA-based methods. Kaiju uses a fast exact matching algorithm based on Burrows-Wheeler Transform (BWT) and FM-index to align translated DNA reads against a reference database of protein sequences.

**See also**

For more information on Kaiju, consult Menzel et al., 2016.

**Warning**

Taxonomic classification can be highly resource-intensive. Ensure that your system has sufficient CPU and memory resources before running these commands.

**For more information on the tools used in this workflow, refer to their official documentation:**

- Kraken 2: https://github.com/DerrickWood/kraken2
- Kaiju: https://github.com/bioinformatics-centre/kaiju

#### 5.4 Taxonomic classification of reads

In this section we will focus on the taxonomic classification of shotgun metagenomic reads using two different tools: Kraken 2 and Kaiju. We will use the data obtained in the *data retrieval section*.

##### 5.4.1 Approach 1: Kraken 2

Before we can use Kraken 2, we need to build or download a database. We will use the `build-kraken-db` action to fetch the PlusPF database from here - this database covers RefSeq sequences for archaea, bacteria, viral, plasmid, human, UniVec_Core, protozoa and fungi.

```
mosh moshpit build-kraken-db \
    --p-collection pluspf \
    --o-kraken2-database ./cache:kraken2_db \
    --o-bracken-database ./cache:bracken_db \
```

We can now use the `classify-kraken2` command to run Kraken2 using the paired-end reads as a query and the PlusPF database retrieved in the previous step:

```
mosh moshpit classify-kraken2 \
    --i-seqs ./cache:reads_filtered \
    --i-kraken2-db ./cache:kraken2_db \
    --p-threads 72 \
    --p-confidence 0.5 \
    --p-memory-mapping False \
    --p-report-minimizer-data \
    --o-reports ./cache:kraken_reports_reads \
    --o-hits ./cache:kraken_hits_reads
    --verbose
```

**See also**

Bracken is a related tool that additionally estimates relative abundances of species or genera to adjust for the genome size the organisms from which each read originated. In order to use this tool we need the Bracken database that was fetched in the first step.

```
mosh moshpit estimate-bracken \
    --i-kraken-reports ./cache:kraken_reports_reads \
    --i-bracken-db ./cache:bracken_db \
    --p-threshold 5 \
    --p-read-len 150 \
    --o-taxonomy ./cache:bracken_taxonomy \
    --o-table ./cache:bracken_ft \
    --o-reports ./cache:bracken_reports
```

To remove the unclassified read fraction we can use the `filter-table` action from the `q2-taxa` QIIME 2 plugin:

```
mosh taxa filter-table \
    --i-table ./cache:bracken_ft \
    --i-taxonomy ./cache:bracken_taxonomy \
    --p-exclude Unclassified \
    --o-filtered-table ./cache:bracken_ft_filtered
```

##### 5.4.2 Approach 2: Kaiju

Similarly to Kraken 2, Kaiju requires a reference database to perform taxonomic classification. We will use the `fetch-kaiju-db` action to download the nr_euk database that includes both prokaryotes and eukaryotes (more info on the taxa here).

```
mosh moshpit fetch-kaiju-db \
    --p-database-type nr_euk \
    --o-database ./cache:kaiju_nr_euk
```

We run Kaiju with the confidence of 0.1 using the paired-end reads as a query and the database artifact that was generated in the previous step:

```
mosh moshpit classify-kaiju \
    --i-seqs ./cache:reads_paired \
    --i-db ./cache:kaiju_nr_euk \
    --p-z 16 \
    --p-c 0.1 \
    --o-taxonomy ./cache:kaiju_taxonomy \
    --o-abundances ./cache:kaiju_ft
```

Finally, we filter the table to remove the unclassified reads:

```
mosh taxa filter-table \
    --i-table ./cache:kaiju_ft \
    --i-taxonomy ./cache:kaiju_taxonomy \
    --p-exclude unclassified,belong,cannot \
    --o-filtered-table ./cache:kaiju_ft_filtered
```

##### 5.4.3 Visualization

You can try to generate a taxa bar plot with either of these results now! We will continue with the Kaiju results - to generate a taxa bar plot, you can run:

```
mosh taxa barplot \
    --i-table ./cache:kaiju_ft_filtered \
    --i-taxonomy ./cache:kaiju_taxonomy \
    --m-metadata-file ./metadata.tsv \
    --o-visualization ./results/kaiju_barplot.qzv
```

Your visualization should look similar to this one.

#### 5.5 Taxonomic classification of MAGs

Kraken 2 can also be used to taxonomically classify metagenome-assembled genomes (MAGs). In this tutorial we use this tool to classify a subset of dereplicated MAGs but the same approach can be used for the entire set of MAGs contained in the `SampleData[MAGs]` or `SampleData[Contigs]` artifacts.

```
mosh moshpit classify-kraken2 \
    --i-seqs ./cache:mags_derep_50 \
    --i-kraken2-db ./cache:kraken2_db \
    --p-threads 72 \
    --p-confidence 0.5 \
    --p-memory-mapping False \
    --p-report-minimizer-data \
    --o-reports ./cache:kraken_reports_mags_derep_50 \
    --o-hits ./cache:kraken_hits_derep_50 \
    --verbose
```

We can now extract the taxonomy from the Kraken 2 reports using the `kraken2-to-mag-features` command:

```
mosh moshpit kraken2-to-mag-features \
    --i-reports ./cache:kraken_reports_mags_derep_50 \
    --i-hits ./cache:kraken_hits_derep_50 \
    --p-coverage-threshold 0.1 \
    --o-taxonomy ./cache:mags_derep_taxonomy_50
```

CHAPTER

SIX

### FUNCTIONAL ANNOTATION

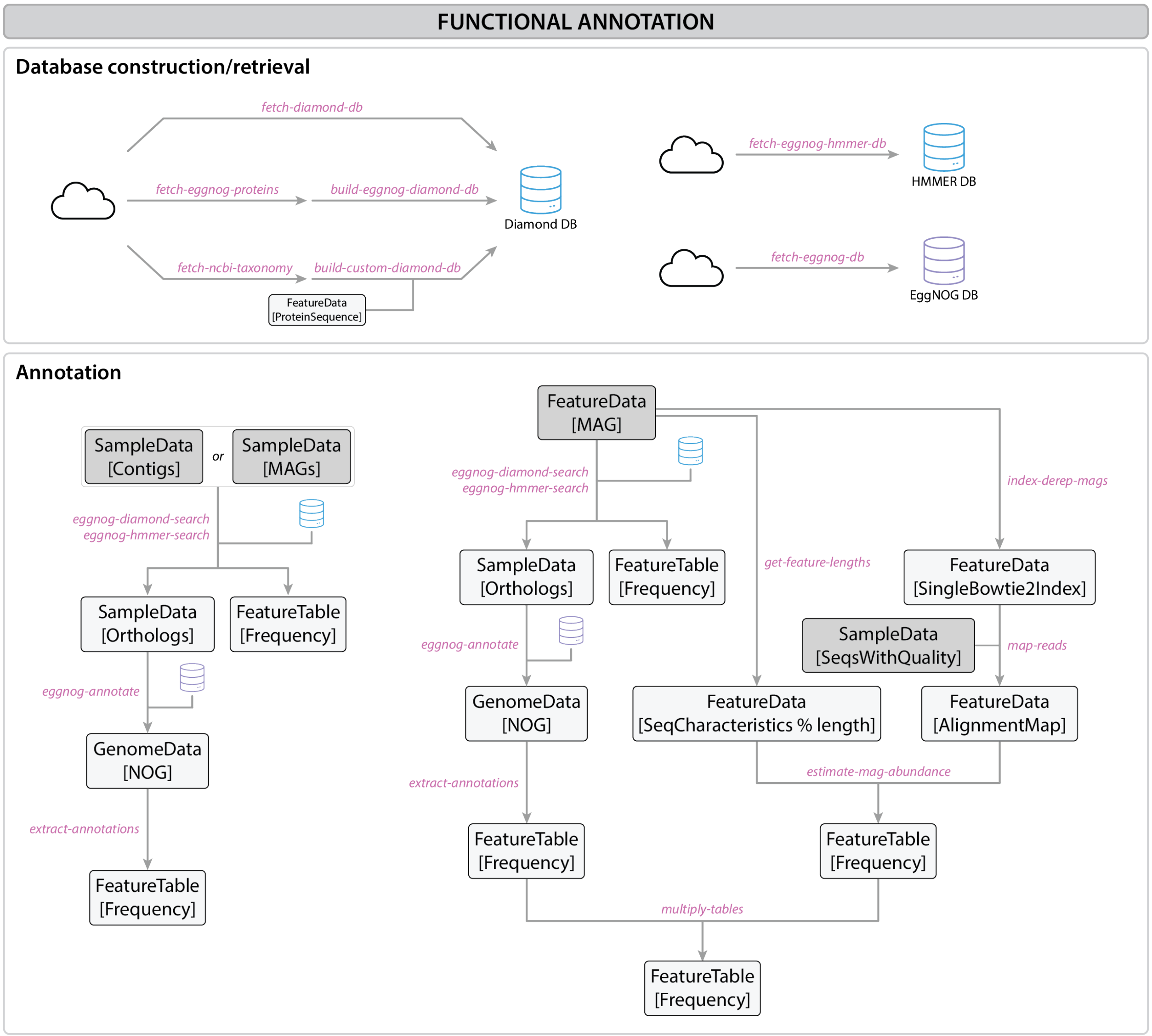

Fig. 6.1: Functional annotation workflow

Functional annotation of **metagenome-assembled genomes (MAGs)** involves the identification and classification of genes within these reconstructed genomes to understand their roles and potential functions. MAGs are recovered from

DNA directly extracted from complex microbial communities, bypassing the need to culture the organisms.

This process provides insights into the genes that code for enzymes, transporters, and other proteins critical to the survival and function of the microbes in various ecosystems. Annotating these genomes allows for the study of their contributions to nutrient cycles, disease processes, or specialized ecological functions, to name only a few examples.

This workflow outlines the step-by-step process for functional annotation of MAGs or contigs using tools like EggNOG and the Diamond aligner in QIIME2.

> **Note**
>
> Functional annotation can be performed on fully reconstructed **MAGs** or directly on **contigs** (the contiguous sequences assembled from sequencing reads). Annotating **contigs** can provide early insights into important functional genes even before complete genomes are assembled. Annotating **MAGs** has the added benefit of seeing how these annotated genes are connected and organized in a single genome.
>
> In this tutorial, we will focus on functional annotation of our previously reconstructed MAGs (see **Recovery of MAGs section**)

> **Warning**
>
> Functional annotation can be highly resource-intensive. Ensure that your system has sufficient CPU and memory resources before running these commands.

**For more information on the tools used in this workflow, refer to their official documentation:**

- EggNOG-mapper: https://github.com/eggnogdb/eggnog-mapper
- DIAMOND: https://github.com/bbuchfink/diamond
- QIIME 2: https://github.com/qiime2

#### 6.1 Functional annotation

##### 6.1.1 Required databases

In order to perform the functional annotation, we will need a couple of different reference databases. Below you will find instructions on how to download these databases using MOSHPIT.

```
mosh moshpit fetch-diamond-db \
    --o-diamond-db ./cache:diamond_db \
    --verbose
```

```
mosh moshpit fetch-eggnog-db \
    --o-eggnog-db ./cache:eggnog_db \
    --verbose
```

Alternatively, you can use:

- `mosh moshpit build-eggnog-diamond-db` to create a DIAMOND formatted reference database for the specified taxon.
- `mosh moshpit build-custom-diamond-db` to create a DIAMOND formatted reference database from a FASTA input file.

##### 6.1.2 EggNOG search using Diamond aligner

We will search the dereplicated MAGs against the EggNOG database using the Diamond aligner to identify functional annotations.

```
mosh moshpit search-orthologs-diamond \
    --i-sequences ./cache:mags_derep \
    --i-diamond-db ./cache:diamond_db \
    --p-num-cpus 16 \
    --p-db-in-memory \
    --o-eggnog-hits ./cache:eggnog_hits \
    --o-table ./cache:eggnog_ft  \
    --verbose
```

##### 6.1.3 Annotate orthologs against eggNOG database

Orthologs from dereplicated MAGs are annotated against the EggNOG database, providing functional insights into the genes and gene products present in the MAGs.

```
mosh moshpit map-eggnog \
    --i-eggnog-hits ./cache:eggnog_hits \
    --i-eggnog-db ./cache:eggnog_db \
    --p-num-cpus 16 \
    --p-db-in-memory \
    --o-ortholog-annotations ./cache:eggnog_annotations \
    --verbose
```

##### 6.1.4 Extract annotations

This method extract a specific annotation from the table generated by EggNOG and calculates its frequencies across all MAGs.

**Note**

The `qiime moshpit extract-annotations` method allows us to extract specific types of functional annotations, such as **CAZymes**, **KEGG pathways**, **COG categories**, or other functional elements, and calculate their frequency across all dereplicated MAGs.

In this tutorial, we focus on demonstrating the extraction of **CAZymes**.

```
mosh moshpit extract-annotations \
    --i-ortholog-annotations ./cache:eggnog_annotations \
    --p-annotation caz \
    --p-max-evalue 0.0001 \
    --o-annotation-frequency ./cache:caz_annot_ft \
    --verbose
```

##### 6.1.5 Multiply tables

This steps simply calculates the dot product of the `mags_derep_ft` and `caz_annot_ft` feature tables. This is useful for combining the annotation data (e.g., **CAZymes**) with MAG abundance to determine how specific functional annotations are distributed across MAGs, and use this information to estimate the total frequency of each annotation in each sample.

```
mosh moshpit multiply-tables \
    --i-table1 ./cache:mags_derep_ft \
    --i-table2 ./cache:caz_annot_ft \
    --o-result-table ./cache:caz_ft \
    --verbose
```

##### 6.1.6 Let's have a look at our CAZymes functional diversity!

We will start by calculating a Bray-curtis dissimilarity matrix to measure the dissimilarity between each sample, based on observed frequency of different CAZyme annotations in each sample.

```
mosh diversity beta \
    --i-table ./cache:caz_ft \
    --p-metric braycurtis \
    --o-distance-matrix ./cache:caz_braycurtis_dist
```

Next, we will perform principal coordinate analysis (PCoA) from the obtained Bray-curtis matrix.

```
mosh diversity pcoa \
    --i-distance-matrix ./cache:caz_braycurtis_dist  \
    --o-pcoa ./cache:caz_braycurtis_pcoa
```

Visualization time! Let's plot the PCoA results.

```
mosh emperor plot \
    --i-pcoa ./cache:caz_braycurtis_dist \
    --m-metadata-file ./metadata.tsv \
    --o-visualization caz-pcoa.qzv
```

Your visualization should look similar to this one.

**Tip**

Once your visualization is ready, click on the `Color` tab at the top right and select `scatter:seed` on the first tab to color your samples by seed type. Then click on the `Animations` tab and choose `timepoint` as gradient and `seed` as trajectory. Now, press play! You should see the progression of samples over time.

CHAPTER

SEVEN

### INTEROPERABILITY WITH OTHER TOOLS

While most of the typical steps in a metagenomic analysis can be performed within QIIME 2, there are cases where you might want to use other tools to perform certain tasks. In this chapter, we will show you how you can get some data in and out of the QIIME 2 artifacts to continue your analysis workflow elsewhere.

#### 7.1 Importing data from other tools

The MOSHPIT pipeline allows you to start working directly with the NGS reads, which you can take through various analysis, like contig assembly, binning, and annotation. However, if you have already performed some of these steps outside of QIIME 2, you can import the results into an appropriate QIIME 2 artifact and continue from there. Below you can see some examples and use cases where this may be relevant.

##### 7.1.1 Working with exisiting contigs

In case you already have contigs assembled from your metagenomic data, you can import them into a `SampleData[Contigs]` artifact. This should not differ much from the typical import process (see here for more details on importing data), but the command may look like:

```
mosh tools cache-import \
    --cache ./cache \
    --key contigs \
    --type "SampleData[Contigs]" \
    --input-path ./<directory with contig FASTA files>
```

Some actions in the MOSHPIT pipeline assume that contig IDs are unique across your entire sample set. If this is not the case, you may use the `qiime assembly rename-contigs` action to rename contigs with unique identifiers:

```
mosh assembly rename-contigs \
    --i-contigs ./cache:contigs \
    --p-uuid-type shortuuid \
    --o-renamed-contigs ./cache:contigs_renamed
```

From here, you should be able to continue with the rest of the MOSHPIT pipeline as described in our tutorials.

##### 7.1.2 Working with existing MAGs

You may also be interested in continuing your analysis with MAGs that you have already recovered using other tools. In this case, you can import the MAGs into a `SampleData[MAGs]` (non-dereplicated) or `FeatureData[MAG]` (dereplicated) artifact. Before you do that, you will need to rename each MAG's FASTA file using the UUID4 format: this is required to ensure that MAG IDs are unique across your entire sample set. Here is a sample Python script which could be used for that purpose:

```
import os
from uuid import uuid4
path = 'path/to/your/mag/directory/'

for file in os.listdir(path):
    os.rename(os.path.join(path, file), os.path.join(path, f'{uuid4()}.fa'))
```

Once you have renamed the MAGs, you can import them into a QIIME 2 artifact:

```
mosh tools cache-import \
    --cache ./cache \
    --key mags \
    --type "SampleData[MAGs]" \
    --input-path ./<directory with MAG FASTA files per sample>
```

for MAGs-per-sample, or:

```
mosh tools cache-import \
    --cache ./cache \
    --key mags \
    --type "FeatureData[MAG]" \
    --input-path ./<directory with MAG FASTA files>
```

for dereplicated MAGs. From here, you should be able to continue with the rest of the MOSHPIT pipeline as described in our tutorials.

##### 7.1.3 Importing other data

If you have other data that you would like to import into QIIME 2, you can use the `qiime tools cache-import` command - no additional steps should be required. For example, you can import a set of Kraken 2 reports into a `SampleData[Kraken2Report % Properties('reads')]` like this:

```
mosh tools cache-import \
    --cache ./cache \
    --key kraken2_reports_reads \
    --type "SampleData[Kraken2Report % reads]" \
    --input-path ./<directory with Kraken 2 reports>
```

> **Note**
>
> Remember: you can import any existing data into QIIME 2 artifacts, as long as it matches the format required by the respective QIIME 2 semantic type.

#### 7.2 Exporting data and connecting with other tools

QIIME 2 offers various ways of visualizing and processing your data further, but sometimes you may want to use other tools that are not (yet) available through QIIME 2. This is, of course, possible and very easy to do: you can export your data from any QIIME 2 artifact and use it with any of your other favourite tools, as long as the underlying format is compatible. The formats that QIIME 2 supports are common and should be readable by most bioinformatics tools - most of the time, the artifacts will contain data in the original format that the underlying tool uses. Below are some examples of how you can export data from QIIME 2 and connect it with other tools.

**Warning**

QIIME 2 does not yet support exporting data from the cache (see below). This means that you will need to manually copy the data from the cache directory to a location where you can access it with other tools. In our examples, the cache directory is located directly in the working directory and that is where we will copy the data from. Keep in mind that you should never temper with the files in the cache directory directly, as this may lead to broken artifacts and failed analyses.

##### 7.2.1 Visualizing Kraken 2 reports with Pavian

If you have used Kraken 2 to *classify your reads*, you can export the resulting reports from the corresponding QIIME 2 artifact and visualize them with Pavian which will allow you to explore the taxonomic composition of your samples in an interactive way. To export the Kraken 2 reports, you can use the following commands:

###### Direct export

Support for extracting data out of the cache is not yet available but is coming soon! You can track the progress of the issue here.

###### Workaround 1: without using QIIME

```
UUID=$(cat ./cache/keys/kraken_reports_reads | grep 'data' | awk '{print $2}')
mkdir ./exported_reports
cp -r ./cache/data/$UUID/data/* ./exported_reports/
```

This will find the UUID of the reports artifact, use it to locate the data within the cache directory, create a directory for the exported data and copy the files from the cache into it.

###### Workaround 2: with QIIME

```
mosh tools cache-fetch \
    --cache ./cache \
    --key kraken_reports_reads \
    --output-path ./kraken_reports_reads.qza

mosh tools export \
    --input-path ./kraken_reports_reads.qza \
    --output-path ./exported_reports
```

This will first fetch the reports artifact from the cache and then export it to the `exported_reports` directory.

Once you got the data into the `exported_reports` directory, you can then use those with Pavian. To give it a quick try, navigate to Pavian's demo site and upload the exported files.

##### 7.2.2 Microbial pangenomics with Anvi'o

Another suite of tools you may be familiar with is the Anvi'o platform. One of the workflows that Anvi'o provides is the microbial pangenomics analysis, which can be used to explore the gene clusters within your samples. You could export the MAGs obtained from the *binning step* and use them as input to the `anvi-pan-genome` workflow, as described here. To export the MAGs, you can use the following command:

###### Direct export

Support for extracting data out of the cache is not yet available but is coming soon! You can track the progress of the issue here.

###### Workaround 1: without using QIIME

```
UUID=$(cat ./cache/keys/mags | grep 'data' | awk '{print $2}')
mkdir ./exported_mags
cp -r ./cache/data/$UUID/data/* ./exported_mags/
```

This will find the UUID of the MAGs artifact, use it to locate the data within the cache directory, create a directory for the exported data and copy the files from the cache into it.

###### Workaround 2: with QIIME

```
mosh tools cache-fetch \
    --cache ./cache \
    --key mags \
    --output-path ./mags.qza

mosh tools export \
    --input-path ./mags.qza \
    --output-path ./exported_mags
```

This will first fetch the MAGs artifact from the cache and then export it to the `exported_mags` directory.

#### TARA Oceans: provenance replay script

```
#!/usr/bin/env bash
##########################################################################################
# Auto-generated by qiime2 v.2024.10.1 at 01:11:43 PM on 29 Nov, 2024
# This document is a representation of the scholarly work of the creator of the
# QIIME 2 Results provided as input to this software, and may be protected by
# intellectual property law. Please respect all copyright restrictions and
# licenses governing the use, modification, and redistribution of this work.

# For User Support, post to the QIIME2 Forum at https://forum.qiime2.org.

# Instructions for use:
# 1. Open this script in a text editor or IDE. Support for BASH
#    syntax highlighting can be helpful.
# 2. Search or scan visually for '<' or '>' characters to find places where
#    user input (e.g. a filepath or column name) is required. These must be
#    replaced with your own values. E.g. <column name> -> 'patient_id'.
#    Failure to remove '<' or '>' may result in `No such File ...` errors
# 3. Search for 'FIXME' comments in the script, and respond as directed.
# 4. Remove all 'FIXME' comments from the script completely. Failure to do so
#    may result in 'Missing Option' errors
# 5. Adjust the arguments to the commands below to suit your data and metadata.
#    If your data is not identical to that in the replayed analysis,
#    changes may be required. (e.g. sample ids or rarefaction depth)
# 6. Optional: replace any filenames in this script that begin with 'XX' with
#    unique file names to ensure they are preserved. QIIME 2 saves all outputs
#    from all actions in this script to disk regardless of whether those
#    outputs were in the original collection of replayed results. The filenames
#    of "un-replayed" artifacts are prefixed with 'XX' so they may be easily
#    located. These names are not guaranteed to be unique, so 'XX_table.qza'
#    may be overwritten by another 'XX_table.qza' later in the script.
# 7. Activate your replay conda environment, and confirm you have installed all
#    plugins used by the script.
# 8. Run this script with `bash <path to this script>`, or copy-paste commands
#    into the terminal for a more interactive analysis.
# 9. Optional: to delete all results not required to produce the figures and
#    data used to generate this script, navigate to the directory in which you
#    ran the script and `rm XX*.qz*`
##########################################################################################
## function to create result collections ##
construct_result_collection () {
	mkdir $rc_name
	touch $rc_name.order
	for key in "${keys[@]}"; do
		echo $key >> $rc_name.order
	done
	for i in "${!keys[@]}"; do
		ln -s ../"${names[i]}" $rc_name"${keys[i]}"$ext
	done
}
##

# This tells bash to -e exit immediately if a command fails
```

```
# and -x show all commands in stdout so you can track progress
set -e -x

qiime tools import \
  --type 'NCBIAccessionIDs' \
  --input-path <your data here> \
  --output-path ncbi-accession-i-ds-0.qza

qiime tools import \
  --type 'NCBIAccessionIDs' \
  --input-path <your data here> \
  --output-path ncbi-accession-i-ds-1.qza

qiime tools import \
  --type 'NCBIAccessionIDs' \
  --input-path <your data here> \
  --output-path ncbi-accession-i-ds-2.qza

qiime tools import \
  --type 'NCBIAccessionIDs' \
  --input-path <your data here> \
  --output-path ncbi-accession-i-ds-3.qza

qiime tools import \
  --type 'NCBIAccessionIDs' \
  --input-path <your data here> \
  --output-path ncbi-accession-i-ds-4.qza

qiime tools import \
  --type 'NCBIAccessionIDs' \
  --input-path <your data here> \
  --output-path ncbi-accession-i-ds-5.qza

qiime tools import \
  --type 'NCBIAccessionIDs' \
  --input-path <your data here> \
  --output-path ncbi-accession-i-ds-6.qza

qiime tools import \
  --type 'NCBIAccessionIDs' \
  --input-path <your data here> \
  --output-path ncbi-accession-i-ds-7.qza

qiime tools import \
  --type 'NCBIAccessionIDs' \
  --input-path <your data here> \
  --output-path ncbi-accession-i-ds-8.qza

qiime tools import \
  --type 'NCBIAccessionIDs' \
  --input-path <your data here> \
  --output-path ncbi-accession-i-ds-9.qza
```

```
qiime tools import \
  --type 'NCBIAccessionIDs' \
  --input-path <your data here> \
  --output-path ncbi-accession-i-ds-10.qza

qiime tools import \
  --type 'NCBIAccessionIDs' \
  --input-path <your data here> \
  --output-path ncbi-accession-i-ds-11.qza

qiime tools import \
  --type 'NCBIAccessionIDs' \
  --input-path <your data here> \
  --output-path ncbi-accession-i-ds-12.qza

qiime tools import \
  --type 'NCBIAccessionIDs' \
  --input-path <your data here> \
  --output-path ncbi-accession-i-ds-13.qza

qiime tools import \
  --type 'NCBIAccessionIDs' \
  --input-path <your data here> \
  --output-path ncbi-accession-i-ds-14.qza

qiime tools import \
  --type 'NCBIAccessionIDs' \
  --input-path <your data here> \
  --output-path ncbi-accession-i-ds-15.qza

qiime tools import \
  --type 'NCBIAccessionIDs' \
  --input-path <your data here> \
  --output-path ncbi-accession-i-ds-16.qza

qiime tools import \
  --type 'NCBIAccessionIDs' \
  --input-path <your data here> \
  --output-path ncbi-accession-i-ds-17.qza

qiime tools import \
  --type 'NCBIAccessionIDs' \
  --input-path <your data here> \
  --output-path ncbi-accession-i-ds-18.qza

qiime tools import \
  --type 'NCBIAccessionIDs' \
  --input-path <your data here> \
  --output-path ncbi-accession-i-ds-19.qza

qiime tools import \
```

```
  --type 'NCBIAccessionIDs' \
  --input-path <your data here> \
  --output-path ncbi-accession-i-ds-20.qza

qiime tools import \
  --type 'NCBIAccessionIDs' \
  --input-path <your data here> \
  --output-path ncbi-accession-i-ds-21.qza

qiime tools import \
  --type 'NCBIAccessionIDs' \
  --input-path <your data here> \
  --output-path ncbi-accession-i-ds-22.qza

qiime tools import \
  --type 'NCBIAccessionIDs' \
  --input-path <your data here> \
  --output-path ncbi-accession-i-ds-23.qza

qiime tools import \
  --type 'NCBIAccessionIDs' \
  --input-path <your data here> \
  --output-path ncbi-accession-i-ds-24.qza

qiime tools import \
  --type 'NCBIAccessionIDs' \
  --input-path <your data here> \
  --output-path ncbi-accession-i-ds-25.qza

qiime tools import \
  --type 'NCBIAccessionIDs' \
  --input-path <your data here> \
  --output-path ncbi-accession-i-ds-26.qza

qiime tools import \
  --type 'NCBIAccessionIDs' \
  --input-path <your data here> \
  --output-path ncbi-accession-i-ds-27.qza

qiime tools import \
  --type 'NCBIAccessionIDs' \
  --input-path <your data here> \
  --output-path ncbi-accession-i-ds-28.qza

qiime tools import \
  --type 'NCBIAccessionIDs' \
  --input-path <your data here> \
  --output-path ncbi-accession-i-ds-29.qza

qiime tools import \
  --type 'NCBIAccessionIDs' \
  --input-path <your data here> \
```

```
  --output-path ncbi-accession-i-ds-30.qza

qiime tools import \
  --type 'NCBIAccessionIDs' \
  --input-path <your data here> \
  --output-path ncbi-accession-i-ds-31.qza

qiime tools import \
  --type 'NCBIAccessionIDs' \
  --input-path <your data here> \
  --output-path ncbi-accession-i-ds-32.qza

qiime tools import \
  --type 'NCBIAccessionIDs' \
  --input-path <your data here> \
  --output-path ncbi-accession-i-ds-33.qza

qiime tools import \
  --type 'NCBIAccessionIDs' \
  --input-path <your data here> \
  --output-path ncbi-accession-i-ds-34.qza

qiime tools import \
  --type 'NCBIAccessionIDs' \
  --input-path <your data here> \
  --output-path ncbi-accession-i-ds-35.qza

qiime tools import \
  --type 'NCBIAccessionIDs' \
  --input-path <your data here> \
  --output-path ncbi-accession-i-ds-36.qza

qiime tools import \
  --type 'NCBIAccessionIDs' \
  --input-path <your data here> \
  --output-path ncbi-accession-i-ds-37.qza

qiime tools import \
  --type 'NCBIAccessionIDs' \
  --input-path <your data here> \
  --output-path ncbi-accession-i-ds-38.qza

qiime tools import \
  --type 'NCBIAccessionIDs' \
  --input-path <your data here> \
  --output-path ncbi-accession-i-ds-39.qza

qiime tools import \
  --type 'NCBIAccessionIDs' \
  --input-path <your data here> \
  --output-path ncbi-accession-i-ds-40.qza
```

```
qiime tools import \
  --type 'NCBIAccessionIDs' \
  --input-path <your data here> \
  --output-path ncbi-accession-i-ds-41.qza

qiime tools import \
  --type 'NCBIAccessionIDs' \
  --input-path <your data here> \
  --output-path ncbi-accession-i-ds-42.qza

qiime tools import \
  --type 'NCBIAccessionIDs' \
  --input-path <your data here> \
  --output-path ncbi-accession-i-ds-43.qza

qiime tools import \
  --type 'NCBIAccessionIDs' \
  --input-path <your data here> \
  --output-path ncbi-accession-i-ds-44.qza

qiime tools import \
  --type 'NCBIAccessionIDs' \
  --input-path <your data here> \
  --output-path ncbi-accession-i-ds-45.qza

qiime tools import \
  --type 'NCBIAccessionIDs' \
  --input-path <your data here> \
  --output-path ncbi-accession-i-ds-46.qza

qiime tools import \
  --type 'NCBIAccessionIDs' \
  --input-path <your data here> \
  --output-path ncbi-accession-i-ds-47.qza

qiime tools import \
  --type 'NCBIAccessionIDs' \
  --input-path <your data here> \
  --output-path ncbi-accession-i-ds-48.qza

qiime tools import \
  --type 'NCBIAccessionIDs' \
  --input-path <your data here> \
  --output-path ncbi-accession-i-ds-49.qza

qiime tools import \
  --type 'NCBIAccessionIDs' \
  --input-path <your data here> \
  --output-path ncbi-accession-i-ds-50.qza

qiime tools import \
  --type 'NCBIAccessionIDs' \
```

```
  --input-path <your data here> \
  --output-path ncbi-accession-i-ds-51.qza

qiime tools import \
  --type 'NCBIAccessionIDs' \
  --input-path <your data here> \
  --output-path ncbi-accession-i-ds-52.qza

qiime tools import \
  --type 'NCBIAccessionIDs' \
  --input-path <your data here> \
  --output-path ncbi-accession-i-ds-53.qza

qiime tools import \
  --type 'NCBIAccessionIDs' \
  --input-path <your data here> \
  --output-path ncbi-accession-i-ds-54.qza

qiime tools import \
  --type 'NCBIAccessionIDs' \
  --input-path <your data here> \
  --output-path ncbi-accession-i-ds-55.qza

qiime tools import \
  --type 'NCBIAccessionIDs' \
  --input-path <your data here> \
  --output-path ncbi-accession-i-ds-56.qza

qiime tools import \
  --type 'NCBIAccessionIDs' \
  --input-path <your data here> \
  --output-path ncbi-accession-i-ds-57.qza

qiime tools import \
  --type 'NCBIAccessionIDs' \
  --input-path <your data here> \
  --output-path ncbi-accession-i-ds-58.qza

qiime tools import \
  --type 'NCBIAccessionIDs' \
  --input-path <your data here> \
  --output-path ncbi-accession-i-ds-59.qza

qiime tools import \
  --type 'NCBIAccessionIDs' \
  --input-path <your data here> \
  --output-path ncbi-accession-i-ds-60.qza

qiime tools import \
  --type 'NCBIAccessionIDs' \
  --input-path <your data here> \
  --output-path ncbi-accession-i-ds-61.qza
```

```
qiime tools import \
  --type 'NCBIAccessionIDs' \
  --input-path <your data here> \
  --output-path ncbi-accession-i-ds-62.qza

qiime tools import \
  --type 'NCBIAccessionIDs' \
  --input-path <your data here> \
  --output-path ncbi-accession-i-ds-63.qza

qiime tools import \
  --type 'NCBIAccessionIDs' \
  --input-path <your data here> \
  --output-path ncbi-accession-i-ds-64.qza

qiime tools import \
  --type 'NCBIAccessionIDs' \
  --input-path <your data here> \
  --output-path ncbi-accession-i-ds-65.qza

qiime tools import \
  --type 'NCBIAccessionIDs' \
  --input-path <your data here> \
  --output-path ncbi-accession-i-ds-66.qza

qiime tools import \
  --type 'NCBIAccessionIDs' \
  --input-path <your data here> \
  --output-path ncbi-accession-i-ds-67.qza

qiime tools import \
  --type 'NCBIAccessionIDs' \
  --input-path <your data here> \
  --output-path ncbi-accession-i-ds-68.qza

qiime tools import \
  --type 'NCBIAccessionIDs' \
  --input-path <your data here> \
  --output-path ncbi-accession-i-ds-69.qza

qiime tools import \
  --type 'NCBIAccessionIDs' \
  --input-path <your data here> \
  --output-path ncbi-accession-i-ds-70.qza

qiime tools import \
  --type 'NCBIAccessionIDs' \
  --input-path <your data here> \
  --output-path ncbi-accession-i-ds-71.qza

qiime tools import \
```

```
  --type 'NCBIAccessionIDs' \
  --input-path <your data here> \
  --output-path ncbi-accession-i-ds-72.qza

qiime tools import \
  --type 'NCBIAccessionIDs' \
  --input-path <your data here> \
  --output-path ncbi-accession-i-ds-73.qza

qiime tools import \
  --type 'NCBIAccessionIDs' \
  --input-path <your data here> \
  --output-path ncbi-accession-i-ds-74.qza

qiime tools import \
  --type 'NCBIAccessionIDs' \
  --input-path <your data here> \
  --output-path ncbi-accession-i-ds-75.qza

qiime tools import \
  --type 'NCBIAccessionIDs' \
  --input-path <your data here> \
  --output-path ncbi-accession-i-ds-76.qza

qiime tools import \
  --type 'NCBIAccessionIDs' \
  --input-path <your data here> \
  --output-path ncbi-accession-i-ds-77.qza

qiime tools import \
  --type 'NCBIAccessionIDs' \
  --input-path <your data here> \
  --output-path ncbi-accession-i-ds-78.qza

qiime tools import \
  --type 'NCBIAccessionIDs' \
  --input-path <your data here> \
  --output-path ncbi-accession-i-ds-79.qza

qiime tools import \
  --type 'NCBIAccessionIDs' \
  --input-path <your data here> \
  --output-path ncbi-accession-i-ds-80.qza

qiime tools import \
  --type 'NCBIAccessionIDs' \
  --input-path <your data here> \
  --output-path ncbi-accession-i-ds-81.qza

qiime tools import \
  --type 'NCBIAccessionIDs' \
  --input-path <your data here> \
```

```
  --output-path ncbi-accession-i-ds-82.qza

qiime tools import \
  --type 'NCBIAccessionIDs' \
  --input-path <your data here> \
  --output-path ncbi-accession-i-ds-83.qza

qiime tools import \
  --type 'NCBIAccessionIDs' \
  --input-path <your data here> \
  --output-path ncbi-accession-i-ds-84.qza

qiime tools import \
  --type 'NCBIAccessionIDs' \
  --input-path <your data here> \
  --output-path ncbi-accession-i-ds-85.qza

qiime tools import \
  --type 'NCBIAccessionIDs' \
  --input-path <your data here> \
  --output-path ncbi-accession-i-ds-86.qza

qiime moshpit build-kraken-db \
  --p-collection pluspfp \
  --p-threads 1 \
  --p-kmer-len 35 \
  --p-minimizer-len 31 \
  --p-minimizer-spaces 7 \
  --p-no-no-masking \
  --p-max-db-size 0 \
  --p-no-use-ftp \
  --p-load-factor 0.7 \
  --p-no-fast-build \
  --o-bracken-database bracken-database-0.qza \
  --o-kraken2-database kraken2-database-0.qza

qiime tools import \
  --type 'NCBIAccessionIDs' \
  --input-path <your data here> \
  --output-path ncbi-accession-i-ds-87.qza

qiime tools import \
  --type 'NCBIAccessionIDs' \
  --input-path <your data here> \
  --output-path ncbi-accession-i-ds-88.qza

qiime tools import \
  --type 'NCBIAccessionIDs' \
  --input-path <your data here> \
  --output-path ncbi-accession-i-ds-89.qza

qiime tools import \
```

```
  --type 'NCBIAccessionIDs' \
  --input-path <your data here> \
  --output-path ncbi-accession-i-ds-90.qza

qiime tools import \
  --type 'NCBIAccessionIDs' \
  --input-path <your data here> \
  --output-path ncbi-accession-i-ds-91.qza

qiime tools import \
  --type 'NCBIAccessionIDs' \
  --input-path <your data here> \
  --output-path ncbi-accession-i-ds-92.qza

qiime tools import \
  --type 'NCBIAccessionIDs' \
  --input-path <your data here> \
  --output-path ncbi-accession-i-ds-93.qza

qiime tools import \
  --type 'NCBIAccessionIDs' \
  --input-path <your data here> \
  --output-path ncbi-accession-i-ds-94.qza

qiime tools import \
  --type 'NCBIAccessionIDs' \
  --input-path <your data here> \
  --output-path ncbi-accession-i-ds-95.qza

qiime tools import \
  --type 'NCBIAccessionIDs' \
  --input-path <your data here> \
  --output-path ncbi-accession-i-ds-96.qza

qiime tools import \
  --type 'NCBIAccessionIDs' \
  --input-path <your data here> \
  --output-path ncbi-accession-i-ds-97.qza

qiime tools import \
  --type 'NCBIAccessionIDs' \
  --input-path <your data here> \
  --output-path ncbi-accession-i-ds-98.qza

qiime tools import \
  --type 'NCBIAccessionIDs' \
  --input-path <your data here> \
  --output-path ncbi-accession-i-ds-99.qza

qiime tools import \
  --type 'NCBIAccessionIDs' \
  --input-path <your data here> \
```

```
  --output-path ncbi-accession-i-ds-100.qza

qiime tools import \
  --type 'NCBIAccessionIDs' \
  --input-path <your data here> \
  --output-path ncbi-accession-i-ds-101.qza

qiime tools import \
  --type 'NCBIAccessionIDs' \
  --input-path <your data here> \
  --output-path ncbi-accession-i-ds-102.qza

qiime tools import \
  --type 'NCBIAccessionIDs' \
  --input-path <your data here> \
  --output-path ncbi-accession-i-ds-103.qza

qiime tools import \
  --type 'NCBIAccessionIDs' \
  --input-path <your data here> \
  --output-path ncbi-accession-i-ds-104.qza

qiime tools import \
  --type 'NCBIAccessionIDs' \
  --input-path <your data here> \
  --output-path ncbi-accession-i-ds-105.qza

qiime tools import \
  --type 'NCBIAccessionIDs' \
  --input-path <your data here> \
  --output-path ncbi-accession-i-ds-106.qza

qiime tools import \
  --type 'NCBIAccessionIDs' \
  --input-path <your data here> \
  --output-path ncbi-accession-i-ds-107.qza

qiime tools import \
  --type 'NCBIAccessionIDs' \
  --input-path <your data here> \
  --output-path ncbi-accession-i-ds-108.qza

qiime tools import \
  --type 'NCBIAccessionIDs' \
  --input-path <your data here> \
  --output-path ncbi-accession-i-ds-109.qza

qiime tools import \
  --type 'NCBIAccessionIDs' \
  --input-path <your data here> \
  --output-path ncbi-accession-i-ds-110.qza
```

```
qiime tools import \
  --type 'NCBIAccessionIDs' \
  --input-path <your data here> \
  --output-path ncbi-accession-i-ds-111.qza

qiime tools import \
  --type 'NCBIAccessionIDs' \
  --input-path <your data here> \
  --output-path ncbi-accession-i-ds-112.qza

qiime tools import \
  --type 'NCBIAccessionIDs' \
  --input-path <your data here> \
  --output-path ncbi-accession-i-ds-113.qza

qiime tools import \
  --type 'NCBIAccessionIDs' \
  --input-path <your data here> \
  --output-path ncbi-accession-i-ds-114.qza

qiime tools import \
  --type 'NCBIAccessionIDs' \
  --input-path <your data here> \
  --output-path ncbi-accession-i-ds-115.qza

qiime tools import \
  --type 'NCBIAccessionIDs' \
  --input-path <your data here> \
  --output-path ncbi-accession-i-ds-116.qza

qiime tools import \
  --type 'NCBIAccessionIDs' \
  --input-path <your data here> \
  --output-path ncbi-accession-i-ds-117.qza

qiime tools import \
  --type 'NCBIAccessionIDs' \
  --input-path <your data here> \
  --output-path ncbi-accession-i-ds-118.qza

qiime tools import \
  --type 'NCBIAccessionIDs' \
  --input-path <your data here> \
  --output-path ncbi-accession-i-ds-119.qza

qiime tools import \
  --type 'NCBIAccessionIDs' \
  --input-path <your data here> \
  --output-path ncbi-accession-i-ds-120.qza

qiime tools import \
  --type 'NCBIAccessionIDs' \
```

```
  --input-path <your data here> \
  --output-path ncbi-accession-i-ds-121.qza

qiime tools import \
  --type 'NCBIAccessionIDs' \
  --input-path <your data here> \
  --output-path ncbi-accession-i-ds-122.qza

qiime tools import \
  --type 'NCBIAccessionIDs' \
  --input-path <your data here> \
  --output-path ncbi-accession-i-ds-123.qza

qiime tools import \
  --type 'NCBIAccessionIDs' \
  --input-path <your data here> \
  --output-path ncbi-accession-i-ds-124.qza

qiime tools import \
  --type 'NCBIAccessionIDs' \
  --input-path <your data here> \
  --output-path ncbi-accession-i-ds-125.qza

qiime tools import \
  --type 'NCBIAccessionIDs' \
  --input-path <your data here> \
  --output-path ncbi-accession-i-ds-126.qza

qiime tools import \
  --type 'NCBIAccessionIDs' \
  --input-path <your data here> \
  --output-path ncbi-accession-i-ds-127.qza

qiime tools import \
  --type 'NCBIAccessionIDs' \
  --input-path <your data here> \
  --output-path ncbi-accession-i-ds-128.qza

qiime tools import \
  --type 'NCBIAccessionIDs' \
  --input-path <your data here> \
  --output-path ncbi-accession-i-ds-129.qza

qiime tools import \
  --type 'NCBIAccessionIDs' \
  --input-path <your data here> \
  --output-path ncbi-accession-i-ds-130.qza

qiime tools import \
  --type 'NCBIAccessionIDs' \
  --input-path <your data here> \
  --output-path ncbi-accession-i-ds-131.qza
```

```
qiime tools import \
  --type 'NCBIAccessionIDs' \
  --input-path <your data here> \
  --output-path ncbi-accession-i-ds-132.qza

qiime tools import \
  --type 'NCBIAccessionIDs' \
  --input-path <your data here> \
  --output-path ncbi-accession-i-ds-133.qza

qiime tools import \
  --type 'NCBIAccessionIDs' \
  --input-path <your data here> \
  --output-path ncbi-accession-i-ds-134.qza

qiime tools import \
  --type 'NCBIAccessionIDs' \
  --input-path <your data here> \
  --output-path ncbi-accession-i-ds-135.qza

qiime tools import \
  --type 'NCBIAccessionIDs' \
  --input-path <your data here> \
  --output-path ncbi-accession-i-ds-136.qza

qiime tools import \
  --type 'NCBIAccessionIDs' \
  --input-path <your data here> \
  --output-path ncbi-accession-i-ds-137.qza

qiime tools import \
  --type 'NCBIAccessionIDs' \
  --input-path <your data here> \
  --output-path ncbi-accession-i-ds-138.qza

qiime tools import \
  --type 'NCBIAccessionIDs' \
  --input-path <your data here> \
  --output-path ncbi-accession-i-ds-139.qza

qiime tools import \
  --type 'NCBIAccessionIDs' \
  --input-path <your data here> \
  --output-path ncbi-accession-i-ds-140.qza

qiime tools import \
  --type 'NCBIAccessionIDs' \
  --input-path <your data here> \
  --output-path ncbi-accession-i-ds-141.qza

qiime tools import \
```

```
  --type 'NCBIAccessionIDs' \
  --input-path <your data here> \
  --output-path ncbi-accession-i-ds-142.qza

qiime tools import \
  --type 'NCBIAccessionIDs' \
  --input-path <your data here> \
  --output-path ncbi-accession-i-ds-143.qza

qiime tools import \
  --type 'NCBIAccessionIDs' \
  --input-path <your data here> \
  --output-path ncbi-accession-i-ds-144.qza

qiime tools import \
  --type 'NCBIAccessionIDs' \
  --input-path <your data here> \
  --output-path ncbi-accession-i-ds-145.qza

qiime tools import \
  --type 'NCBIAccessionIDs' \
  --input-path <your data here> \
  --output-path ncbi-accession-i-ds-146.qza

qiime tools import \
  --type 'NCBIAccessionIDs' \
  --input-path <your data here> \
  --output-path ncbi-accession-i-ds-147.qza

qiime tools import \
  --type 'NCBIAccessionIDs' \
  --input-path <your data here> \
  --output-path ncbi-accession-i-ds-148.qza

qiime tools import \
  --type 'NCBIAccessionIDs' \
  --input-path <your data here> \
  --output-path ncbi-accession-i-ds-149.qza

qiime tools import \
  --type 'NCBIAccessionIDs' \
  --input-path <your data here> \
  --output-path ncbi-accession-i-ds-150.qza

qiime tools import \
  --type 'NCBIAccessionIDs' \
  --input-path <your data here> \
  --output-path ncbi-accession-i-ds-151.qza

qiime tools import \
  --type 'NCBIAccessionIDs' \
  --input-path <your data here> \
```

```
  --output-path ncbi-accession-i-ds-152.qza

qiime tools import \
  --type 'NCBIAccessionIDs' \
  --input-path <your data here> \
  --output-path ncbi-accession-i-ds-153.qza

qiime tools import \
  --type 'NCBIAccessionIDs' \
  --input-path <your data here> \
  --output-path ncbi-accession-i-ds-154.qza

qiime tools import \
  --type 'NCBIAccessionIDs' \
  --input-path <your data here> \
  --output-path ncbi-accession-i-ds-155.qza

qiime tools import \
  --type 'NCBIAccessionIDs' \
  --input-path <your data here> \
  --output-path ncbi-accession-i-ds-156.qza

qiime tools import \
  --type 'NCBIAccessionIDs' \
  --input-path <your data here> \
  --output-path ncbi-accession-i-ds-157.qza

qiime tools import \
  --type 'NCBIAccessionIDs' \
  --input-path <your data here> \
  --output-path ncbi-accession-i-ds-158.qza

qiime tools import \
  --type 'NCBIAccessionIDs' \
  --input-path <your data here> \
  --output-path ncbi-accession-i-ds-159.qza

qiime tools import \
  --type 'NCBIAccessionIDs' \
  --input-path <your data here> \
  --output-path ncbi-accession-i-ds-160.qza

qiime tools import \
  --type 'NCBIAccessionIDs' \
  --input-path <your data here> \
  --output-path ncbi-accession-i-ds-161.qza

qiime tools import \
  --type 'NCBIAccessionIDs' \
  --input-path <your data here> \
  --output-path ncbi-accession-i-ds-162.qza
```

```
qiime tools import \
  --type 'NCBIAccessionIDs' \
  --input-path <your data here> \
  --output-path ncbi-accession-i-ds-163.qza

qiime tools import \
  --type 'NCBIAccessionIDs' \
  --input-path <your data here> \
  --output-path ncbi-accession-i-ds-164.qza

qiime tools import \
  --type 'NCBIAccessionIDs' \
  --input-path <your data here> \
  --output-path ncbi-accession-i-ds-165.qza

qiime tools import \
  --type 'NCBIAccessionIDs' \
  --input-path <your data here> \
  --output-path ncbi-accession-i-ds-166.qza

qiime tools import \
  --type 'NCBIAccessionIDs' \
  --input-path <your data here> \
  --output-path ncbi-accession-i-ds-167.qza

qiime tools import \
  --type 'NCBIAccessionIDs' \
  --input-path <your data here> \
  --output-path ncbi-accession-i-ds-168.qza

qiime tools import \
  --type 'NCBIAccessionIDs' \
  --input-path <your data here> \
  --output-path ncbi-accession-i-ds-169.qza

qiime tools import \
  --type 'NCBIAccessionIDs' \
  --input-path <your data here> \
  --output-path ncbi-accession-i-ds-170.qza

qiime tools import \
  --type 'NCBIAccessionIDs' \
  --input-path <your data here> \
  --output-path ncbi-accession-i-ds-171.qza

qiime tools import \
  --type 'NCBIAccessionIDs' \
  --input-path <your data here> \
  --output-path ncbi-accession-i-ds-172.qza

qiime tools import \
  --type 'NCBIAccessionIDs' \
```

```
  --input-path <your data here> \
  --output-path ncbi-accession-i-ds-173.qza

qiime tools import \
  --type 'NCBIAccessionIDs' \
  --input-path <your data here> \
  --output-path ncbi-accession-i-ds-174.qza

qiime tools import \
  --type 'NCBIAccessionIDs' \
  --input-path <your data here> \
  --output-path ncbi-accession-i-ds-175.qza

qiime tools import \
  --type 'NCBIAccessionIDs' \
  --input-path <your data here> \
  --output-path ncbi-accession-i-ds-176.qza

qiime tools import \
  --type 'NCBIAccessionIDs' \
  --input-path <your data here> \
  --output-path ncbi-accession-i-ds-177.qza

qiime tools import \
  --type 'NCBIAccessionIDs' \
  --input-path <your data here> \
  --output-path ncbi-accession-i-ds-178.qza

qiime tools import \
  --type 'NCBIAccessionIDs' \
  --input-path <your data here> \
  --output-path ncbi-accession-i-ds-179.qza

qiime tools import \
  --type 'NCBIAccessionIDs' \
  --input-path <your data here> \
  --output-path ncbi-accession-i-ds-180.qza

qiime tools import \
  --type 'NCBIAccessionIDs' \
  --input-path <your data here> \
  --output-path ncbi-accession-i-ds-181.qza

qiime tools import \
  --type 'NCBIAccessionIDs' \
  --input-path <your data here> \
  --output-path ncbi-accession-i-ds-182.qza

qiime tools import \
  --type 'NCBIAccessionIDs' \
  --input-path <your data here> \
  --output-path ncbi-accession-i-ds-183.qza
```

```
qiime tools import \
  --type 'NCBIAccessionIDs' \
  --input-path <your data here> \
  --output-path ncbi-accession-i-ds-184.qza

qiime tools import \
  --type 'NCBIAccessionIDs' \
  --input-path <your data here> \
  --output-path ncbi-accession-i-ds-185.qza

qiime tools import \
  --type 'NCBIAccessionIDs' \
  --input-path <your data here> \
  --output-path ncbi-accession-i-ds-186.qza

qiime tools import \
  --type 'NCBIAccessionIDs' \
  --input-path <your data here> \
  --output-path ncbi-accession-i-ds-187.qza

qiime tools import \
  --type 'NCBIAccessionIDs' \
  --input-path <your data here> \
  --output-path ncbi-accession-i-ds-188.qza

qiime tools import \
  --type 'NCBIAccessionIDs' \
  --input-path <your data here> \
  --output-path ncbi-accession-i-ds-189.qza

qiime tools import \
  --type 'NCBIAccessionIDs' \
  --input-path <your data here> \
  --output-path ncbi-accession-i-ds-190.qza

qiime tools import \
  --type 'NCBIAccessionIDs' \
  --input-path <your data here> \
  --output-path ncbi-accession-i-ds-191.qza

qiime tools import \
  --type 'NCBIAccessionIDs' \
  --input-path <your data here> \
  --output-path ncbi-accession-i-ds-192.qza

qiime tools import \
  --type 'NCBIAccessionIDs' \
  --input-path <your data here> \
  --output-path ncbi-accession-i-ds-193.qza

qiime tools import \
```

```
  --type 'NCBIAccessionIDs' \
  --input-path <your data here> \
  --output-path ncbi-accession-i-ds-194.qza

qiime tools import \
  --type 'NCBIAccessionIDs' \
  --input-path <your data here> \
  --output-path ncbi-accession-i-ds-195.qza

qiime tools import \
  --type 'NCBIAccessionIDs' \
  --input-path <your data here> \
  --output-path ncbi-accession-i-ds-196.qza

qiime tools import \
  --type 'NCBIAccessionIDs' \
  --input-path <your data here> \
  --output-path ncbi-accession-i-ds-197.qza

qiime tools import \
  --type 'NCBIAccessionIDs' \
  --input-path <your data here> \
  --output-path ncbi-accession-i-ds-198.qza

qiime tools import \
  --type 'NCBIAccessionIDs' \
  --input-path <your data here> \
  --output-path ncbi-accession-i-ds-199.qza

qiime tools import \
  --type 'NCBIAccessionIDs' \
  --input-path <your data here> \
  --output-path ncbi-accession-i-ds-200.qza

qiime tools import \
  --type 'NCBIAccessionIDs' \
  --input-path <your data here> \
  --output-path ncbi-accession-i-ds-201.qza

qiime tools import \
  --type 'NCBIAccessionIDs' \
  --input-path <your data here> \
  --output-path ncbi-accession-i-ds-202.qza

qiime tools import \
  --type 'NCBIAccessionIDs' \
  --input-path <your data here> \
  --output-path ncbi-accession-i-ds-203.qza

qiime tools import \
  --type 'NCBIAccessionIDs' \
  --input-path <your data here> \
```

```
  --output-path ncbi-accession-i-ds-204.qza

qiime tools import \
  --type 'NCBIAccessionIDs' \
  --input-path <your data here> \
  --output-path ncbi-accession-i-ds-205.qza

qiime tools import \
  --type 'NCBIAccessionIDs' \
  --input-path <your data here> \
  --output-path ncbi-accession-i-ds-206.qza

qiime tools import \
  --type 'NCBIAccessionIDs' \
  --input-path <your data here> \
  --output-path ncbi-accession-i-ds-207.qza

qiime tools import \
  --type 'NCBIAccessionIDs' \
  --input-path <your data here> \
  --output-path ncbi-accession-i-ds-208.qza

qiime tools import \
  --type 'NCBIAccessionIDs' \
  --input-path <your data here> \
  --output-path ncbi-accession-i-ds-209.qza

qiime tools import \
  --type 'NCBIAccessionIDs' \
  --input-path <your data here> \
  --output-path ncbi-accession-i-ds-210.qza

qiime tools import \
  --type 'NCBIAccessionIDs' \
  --input-path <your data here> \
  --output-path ncbi-accession-i-ds-211.qza

qiime tools import \
  --type 'NCBIAccessionIDs' \
  --input-path <your data here> \
  --output-path ncbi-accession-i-ds-212.qza

qiime tools import \
  --type 'NCBIAccessionIDs' \
  --input-path <your data here> \
  --output-path ncbi-accession-i-ds-213.qza

qiime tools import \
  --type 'NCBIAccessionIDs' \
  --input-path <your data here> \
  --output-path ncbi-accession-i-ds-214.qza
```

```
qiime tools import \
  --type 'NCBIAccessionIDs' \
  --input-path <your data here> \
  --output-path ncbi-accession-i-ds-215.qza

qiime tools import \
  --type 'NCBIAccessionIDs' \
  --input-path <your data here> \
  --output-path ncbi-accession-i-ds-216.qza

qiime tools import \
  --type 'NCBIAccessionIDs' \
  --input-path <your data here> \
  --output-path ncbi-accession-i-ds-217.qza

qiime tools import \
  --type 'NCBIAccessionIDs' \
  --input-path <your data here> \
  --output-path ncbi-accession-i-ds-218.qza

qiime tools import \
  --type 'NCBIAccessionIDs' \
  --input-path <your data here> \
  --output-path ncbi-accession-i-ds-219.qza

qiime tools import \
  --type 'NCBIAccessionIDs' \
  --input-path <your data here> \
  --output-path ncbi-accession-i-ds-220.qza

qiime tools import \
  --type 'NCBIAccessionIDs' \
  --input-path <your data here> \
  --output-path ncbi-accession-i-ds-221.qza

qiime tools import \
  --type 'NCBIAccessionIDs' \
  --input-path <your data here> \
  --output-path ncbi-accession-i-ds-222.qza

qiime tools import \
  --type 'NCBIAccessionIDs' \
  --input-path <your data here> \
  --output-path ncbi-accession-i-ds-223.qza

qiime tools import \
  --type 'NCBIAccessionIDs' \
  --input-path <your data here> \
  --output-path ncbi-accession-i-ds-224.qza

qiime tools import \
  --type 'NCBIAccessionIDs' \
```

```
  --input-path <your data here> \
  --output-path ncbi-accession-i-ds-225.qza

qiime tools import \
  --type 'NCBIAccessionIDs' \
  --input-path <your data here> \
  --output-path ncbi-accession-i-ds-226.qza

qiime tools import \
  --type 'NCBIAccessionIDs' \
  --input-path <your data here> \
  --output-path ncbi-accession-i-ds-227.qza

qiime tools import \
  --type 'NCBIAccessionIDs' \
  --input-path <your data here> \
  --output-path ncbi-accession-i-ds-228.qza

qiime tools import \
  --type 'NCBIAccessionIDs' \
  --input-path <your data here> \
  --output-path ncbi-accession-i-ds-229.qza

qiime tools import \
  --type 'NCBIAccessionIDs' \
  --input-path <your data here> \
  --output-path ncbi-accession-i-ds-230.qza

qiime tools import \
  --type 'NCBIAccessionIDs' \
  --input-path <your data here> \
  --output-path ncbi-accession-i-ds-231.qza

qiime tools import \
  --type 'NCBIAccessionIDs' \
  --input-path <your data here> \
  --output-path ncbi-accession-i-ds-232.qza

qiime tools import \
  --type 'NCBIAccessionIDs' \
  --input-path <your data here> \
  --output-path ncbi-accession-i-ds-233.qza

qiime tools import \
  --type 'NCBIAccessionIDs' \
  --input-path <your data here> \
  --output-path ncbi-accession-i-ds-234.qza

qiime tools import \
  --type 'NCBIAccessionIDs' \
  --input-path <your data here> \
  --output-path ncbi-accession-i-ds-235.qza
```

```
qiime tools import \
  --type 'NCBIAccessionIDs' \
  --input-path <your data here> \
  --output-path ncbi-accession-i-ds-236.qza

qiime tools import \
  --type 'NCBIAccessionIDs' \
  --input-path <your data here> \
  --output-path ncbi-accession-i-ds-237.qza

qiime tools import \
  --type 'NCBIAccessionIDs' \
  --input-path <your data here> \
  --output-path ncbi-accession-i-ds-238.qza

qiime tools import \
  --type 'NCBIAccessionIDs' \
  --input-path <your data here> \
  --output-path ncbi-accession-i-ds-239.qza

qiime tools import \
  --type 'NCBIAccessionIDs' \
  --input-path <your data here> \
  --output-path ncbi-accession-i-ds-240.qza

qiime tools import \
  --type 'NCBIAccessionIDs' \
  --input-path <your data here> \
  --output-path ncbi-accession-i-ds-241.qza

qiime tools import \
  --type 'NCBIAccessionIDs' \
  --input-path <your data here> \
  --output-path ncbi-accession-i-ds-242.qza

qiime fondue get-sequences \
  --i-accession-ids ncbi-accession-i-ds-0.qza \
  --p- \
  --p-retries 5 \
  --p-n-jobs 1 \
  --p-log-level DEBUG \
  --p-no-restricted-access \
  --o-paired-reads paired-reads-0.qza \
  --o-single-reads XX_single_reads \
  --o-failed-runs XX_failed_runs

qiime fondue get-sequences \
  --i-accession-ids ncbi-accession-i-ds-1.qza \
  --p- \
  --p-retries 5 \
  --p-n-jobs 1 \
```

```
  --p-log-level DEBUG \
  --p-no-restricted-access \
  --o-paired-reads paired-reads-1.qza \
  --o-single-reads XX_single_reads \
  --o-failed-runs XX_failed_runs

qiime fondue get-sequences \
  --i-accession-ids ncbi-accession-i-ds-2.qza \
  --p- \
  --p-retries 5 \
  --p-n-jobs 1 \
  --p-log-level DEBUG \
  --p-no-restricted-access \
  --o-paired-reads paired-reads-2.qza \
  --o-single-reads XX_single_reads \
  --o-failed-runs XX_failed_runs

qiime fondue get-sequences \
  --i-accession-ids ncbi-accession-i-ds-3.qza \
  --p- \
  --p-retries 5 \
  --p-n-jobs 1 \
  --p-log-level DEBUG \
  --p-no-restricted-access \
  --o-paired-reads paired-reads-3.qza \
  --o-single-reads XX_single_reads \
  --o-failed-runs XX_failed_runs

qiime fondue get-sequences \
  --i-accession-ids ncbi-accession-i-ds-4.qza \
  --p- \
  --p-retries 5 \
  --p-n-jobs 1 \
  --p-log-level DEBUG \
  --p-no-restricted-access \
  --o-paired-reads paired-reads-4.qza \
  --o-single-reads XX_single_reads \
  --o-failed-runs XX_failed_runs

qiime fondue get-sequences \
  --i-accession-ids ncbi-accession-i-ds-5.qza \
  --p- \
  --p-retries 5 \
  --p-n-jobs 1 \
  --p-log-level DEBUG \
  --p-no-restricted-access \
  --o-paired-reads paired-reads-5.qza \
  --o-single-reads XX_single_reads \
  --o-failed-runs XX_failed_runs

qiime fondue get-sequences \
  --i-accession-ids ncbi-accession-i-ds-6.qza \
```

```
  --p- \
  --p-retries 5 \
  --p-n-jobs 1 \
  --p-log-level DEBUG \
  --p-no-restricted-access \
  --o-paired-reads paired-reads-6.qza \
  --o-single-reads XX_single_reads \
  --o-failed-runs XX_failed_runs

qiime fondue get-sequences \
  --i-accession-ids ncbi-accession-i-ds-7.qza \
  --p- \
  --p-retries 5 \
  --p-n-jobs 1 \
  --p-log-level DEBUG \
  --p-no-restricted-access \
  --o-paired-reads paired-reads-7.qza \
  --o-single-reads XX_single_reads \
  --o-failed-runs XX_failed_runs

qiime fondue get-sequences \
  --i-accession-ids ncbi-accession-i-ds-8.qza \
  --p- \
  --p-retries 5 \
  --p-n-jobs 1 \
  --p-log-level DEBUG \
  --p-no-restricted-access \
  --o-paired-reads paired-reads-8.qza \
  --o-single-reads XX_single_reads \
  --o-failed-runs XX_failed_runs

qiime fondue get-sequences \
  --i-accession-ids ncbi-accession-i-ds-9.qza \
  --p- \
  --p-retries 5 \
  --p-n-jobs 1 \
  --p-log-level DEBUG \
  --p-no-restricted-access \
  --o-paired-reads paired-reads-9.qza \
  --o-single-reads XX_single_reads \
  --o-failed-runs XX_failed_runs

qiime fondue get-sequences \
  --i-accession-ids ncbi-accession-i-ds-10.qza \
  --p- \
  --p-retries 5 \
  --p-n-jobs 1 \
  --p-log-level DEBUG \
  --p-no-restricted-access \
  --o-paired-reads paired-reads-10.qza \
  --o-single-reads XX_single_reads \
  --o-failed-runs XX_failed_runs
```

```
qiime fondue get-sequences \
  --i-accession-ids ncbi-accession-i-ds-11.qza \
  --p- \
  --p-retries 5 \
  --p-n-jobs 1 \
  --p-log-level DEBUG \
  --p-no-restricted-access \
  --o-paired-reads paired-reads-11.qza \
  --o-single-reads XX_single_reads \
  --o-failed-runs XX_failed_runs

qiime fondue get-sequences \
  --i-accession-ids ncbi-accession-i-ds-12.qza \
  --p- \
  --p-retries 5 \
  --p-n-jobs 1 \
  --p-log-level DEBUG \
  --p-no-restricted-access \
  --o-paired-reads paired-reads-12.qza \
  --o-single-reads XX_single_reads \
  --o-failed-runs XX_failed_runs

qiime fondue get-sequences \
  --i-accession-ids ncbi-accession-i-ds-13.qza \
  --p- \
  --p-retries 5 \
  --p-n-jobs 1 \
  --p-log-level DEBUG \
  --p-no-restricted-access \
  --o-paired-reads paired-reads-13.qza \
  --o-single-reads XX_single_reads \
  --o-failed-runs XX_failed_runs

qiime fondue get-sequences \
  --i-accession-ids ncbi-accession-i-ds-14.qza \
  --p- \
  --p-retries 5 \
  --p-n-jobs 1 \
  --p-log-level DEBUG \
  --p-no-restricted-access \
  --o-paired-reads paired-reads-14.qza \
  --o-single-reads XX_single_reads \
  --o-failed-runs XX_failed_runs

qiime fondue get-sequences \
  --i-accession-ids ncbi-accession-i-ds-15.qza \
  --p- \
  --p-retries 5 \
  --p-n-jobs 1 \
  --p-log-level DEBUG \
  --p-no-restricted-access \
```

```
  --o-paired-reads paired-reads-15.qza \
  --o-single-reads XX_single_reads \
  --o-failed-runs XX_failed_runs

qiime fondue get-sequences \
  --i-accession-ids ncbi-accession-i-ds-16.qza \
  --p- \
  --p-retries 5 \
  --p-n-jobs 1 \
  --p-log-level DEBUG \
  --p-no-restricted-access \
  --o-paired-reads paired-reads-16.qza \
  --o-single-reads XX_single_reads \
  --o-failed-runs XX_failed_runs

qiime fondue get-sequences \
  --i-accession-ids ncbi-accession-i-ds-17.qza \
  --p- \
  --p-retries 5 \
  --p-n-jobs 1 \
  --p-log-level DEBUG \
  --p-no-restricted-access \
  --o-paired-reads paired-reads-17.qza \
  --o-single-reads XX_single_reads \
  --o-failed-runs XX_failed_runs

qiime fondue get-sequences \
  --i-accession-ids ncbi-accession-i-ds-18.qza \
  --p- \
  --p-retries 5 \
  --p-n-jobs 1 \
  --p-log-level DEBUG \
  --p-no-restricted-access \
  --o-paired-reads paired-reads-18.qza \
  --o-single-reads XX_single_reads \
  --o-failed-runs XX_failed_runs

qiime fondue get-sequences \
  --i-accession-ids ncbi-accession-i-ds-19.qza \
  --p- \
  --p-retries 5 \
  --p-n-jobs 1 \
  --p-log-level DEBUG \
  --p-no-restricted-access \
  --o-paired-reads paired-reads-19.qza \
  --o-single-reads XX_single_reads \
  --o-failed-runs XX_failed_runs

qiime fondue get-sequences \
  --i-accession-ids ncbi-accession-i-ds-20.qza \
  --p- \
  --p-retries 5 \
```

```
  --p-n-jobs 1 \
  --p-log-level DEBUG \
  --p-no-restricted-access \
  --o-paired-reads paired-reads-20.qza \
  --o-single-reads XX_single_reads \
  --o-failed-runs XX_failed_runs

qiime fondue get-sequences \
  --i-accession-ids ncbi-accession-i-ds-21.qza \
  --p- \
  --p-retries 5 \
  --p-n-jobs 1 \
  --p-log-level DEBUG \
  --p-no-restricted-access \
  --o-paired-reads paired-reads-21.qza \
  --o-single-reads XX_single_reads \
  --o-failed-runs XX_failed_runs

qiime fondue get-sequences \
  --i-accession-ids ncbi-accession-i-ds-22.qza \
  --p- \
  --p-retries 5 \
  --p-n-jobs 1 \
  --p-log-level DEBUG \
  --p-no-restricted-access \
  --o-paired-reads paired-reads-22.qza \
  --o-single-reads XX_single_reads \
  --o-failed-runs XX_failed_runs

qiime fondue get-sequences \
  --i-accession-ids ncbi-accession-i-ds-23.qza \
  --p- \
  --p-retries 5 \
  --p-n-jobs 1 \
  --p-log-level DEBUG \
  --p-no-restricted-access \
  --o-paired-reads paired-reads-23.qza \
  --o-single-reads XX_single_reads \
  --o-failed-runs XX_failed_runs

qiime fondue get-sequences \
  --i-accession-ids ncbi-accession-i-ds-24.qza \
  --p- \
  --p-retries 5 \
  --p-n-jobs 1 \
  --p-log-level DEBUG \
  --p-no-restricted-access \
  --o-paired-reads paired-reads-24.qza \
  --o-single-reads XX_single_reads \
  --o-failed-runs XX_failed_runs

qiime fondue get-sequences \
```

```
  --i-accession-ids ncbi-accession-i-ds-25.qza \
  --p- \
  --p-retries 5 \
  --p-n-jobs 1 \
  --p-log-level DEBUG \
  --p-no-restricted-access \
  --o-paired-reads paired-reads-25.qza \
  --o-single-reads XX_single_reads \
  --o-failed-runs XX_failed_runs

qiime fondue get-sequences \
  --i-accession-ids ncbi-accession-i-ds-26.qza \
  --p- \
  --p-retries 5 \
  --p-n-jobs 1 \
  --p-log-level DEBUG \
  --p-no-restricted-access \
  --o-paired-reads paired-reads-26.qza \
  --o-single-reads XX_single_reads \
  --o-failed-runs XX_failed_runs

qiime fondue get-sequences \
  --i-accession-ids ncbi-accession-i-ds-27.qza \
  --p- \
  --p-retries 5 \
  --p-n-jobs 1 \
  --p-log-level DEBUG \
  --p-no-restricted-access \
  --o-paired-reads paired-reads-27.qza \
  --o-single-reads XX_single_reads \
  --o-failed-runs XX_failed_runs

qiime fondue get-sequences \
  --i-accession-ids ncbi-accession-i-ds-28.qza \
  --p- \
  --p-retries 5 \
  --p-n-jobs 1 \
  --p-log-level DEBUG \
  --p-no-restricted-access \
  --o-paired-reads paired-reads-28.qza \
  --o-single-reads XX_single_reads \
  --o-failed-runs XX_failed_runs

qiime fondue get-sequences \
  --i-accession-ids ncbi-accession-i-ds-29.qza \
  --p- \
  --p-retries 5 \
  --p-n-jobs 1 \
  --p-log-level DEBUG \
  --p-no-restricted-access \
  --o-paired-reads paired-reads-29.qza \
  --o-single-reads XX_single_reads \
```

```
  --o-failed-runs XX_failed_runs

qiime fondue get-sequences \
  --i-accession-ids ncbi-accession-i-ds-30.qza \
  --p- \
  --p-retries 5 \
  --p-n-jobs 1 \
  --p-log-level DEBUG \
  --p-no-restricted-access \
  --o-paired-reads paired-reads-30.qza \
  --o-single-reads XX_single_reads \
  --o-failed-runs XX_failed_runs

qiime fondue get-sequences \
  --i-accession-ids ncbi-accession-i-ds-31.qza \
  --p- \
  --p-retries 5 \
  --p-n-jobs 1 \
  --p-log-level DEBUG \
  --p-no-restricted-access \
  --o-paired-reads paired-reads-31.qza \
  --o-single-reads XX_single_reads \
  --o-failed-runs XX_failed_runs

qiime fondue get-sequences \
  --i-accession-ids ncbi-accession-i-ds-32.qza \
  --p- \
  --p-retries 5 \
  --p-n-jobs 1 \
  --p-log-level DEBUG \
  --p-no-restricted-access \
  --o-paired-reads paired-reads-32.qza \
  --o-single-reads XX_single_reads \
  --o-failed-runs XX_failed_runs

qiime fondue get-sequences \
  --i-accession-ids ncbi-accession-i-ds-33.qza \
  --p- \
  --p-retries 5 \
  --p-n-jobs 1 \
  --p-log-level DEBUG \
  --p-no-restricted-access \
  --o-paired-reads paired-reads-33.qza \
  --o-single-reads XX_single_reads \
  --o-failed-runs XX_failed_runs

qiime fondue get-sequences \
  --i-accession-ids ncbi-accession-i-ds-34.qza \
  --p- \
  --p-retries 5 \
  --p-n-jobs 1 \
  --p-log-level DEBUG \
```

```
  --p-no-restricted-access \
  --o-paired-reads paired-reads-34.qza \
  --o-single-reads XX_single_reads \
  --o-failed-runs XX_failed_runs

qiime fondue get-sequences \
  --i-accession-ids ncbi-accession-i-ds-35.qza \
  --p- \
  --p-retries 5 \
  --p-n-jobs 1 \
  --p-log-level DEBUG \
  --p-no-restricted-access \
  --o-paired-reads paired-reads-35.qza \
  --o-single-reads XX_single_reads \
  --o-failed-runs XX_failed_runs

qiime fondue get-sequences \
  --i-accession-ids ncbi-accession-i-ds-36.qza \
  --p- \
  --p-retries 5 \
  --p-n-jobs 1 \
  --p-log-level DEBUG \
  --p-no-restricted-access \
  --o-paired-reads paired-reads-36.qza \
  --o-single-reads XX_single_reads \
  --o-failed-runs XX_failed_runs

qiime fondue get-sequences \
  --i-accession-ids ncbi-accession-i-ds-37.qza \
  --p- \
  --p-retries 5 \
  --p-n-jobs 1 \
  --p-log-level DEBUG \
  --p-no-restricted-access \
  --o-paired-reads paired-reads-37.qza \
  --o-single-reads XX_single_reads \
  --o-failed-runs XX_failed_runs

qiime fondue get-sequences \
  --i-accession-ids ncbi-accession-i-ds-38.qza \
  --p- \
  --p-retries 5 \
  --p-n-jobs 1 \
  --p-log-level DEBUG \
  --p-no-restricted-access \
  --o-paired-reads paired-reads-38.qza \
  --o-single-reads XX_single_reads \
  --o-failed-runs XX_failed_runs

qiime fondue get-sequences \
  --i-accession-ids ncbi-accession-i-ds-39.qza \
  --p- \
```

```
  --p-retries 5 \
  --p-n-jobs 1 \
  --p-log-level DEBUG \
  --p-no-restricted-access \
  --o-paired-reads paired-reads-39.qza \
  --o-single-reads XX_single_reads \
  --o-failed-runs XX_failed_runs

qiime fondue get-sequences \
  --i-accession-ids ncbi-accession-i-ds-40.qza \
  --p- \
  --p-retries 5 \
  --p-n-jobs 1 \
  --p-log-level DEBUG \
  --p-no-restricted-access \
  --o-paired-reads paired-reads-40.qza \
  --o-single-reads XX_single_reads \
  --o-failed-runs XX_failed_runs

qiime fondue get-sequences \
  --i-accession-ids ncbi-accession-i-ds-41.qza \
  --p- \
  --p-retries 5 \
  --p-n-jobs 1 \
  --p-log-level DEBUG \
  --p-no-restricted-access \
  --o-paired-reads paired-reads-41.qza \
  --o-single-reads XX_single_reads \
  --o-failed-runs XX_failed_runs

qiime fondue get-sequences \
  --i-accession-ids ncbi-accession-i-ds-42.qza \
  --p- \
  --p-retries 5 \
  --p-n-jobs 1 \
  --p-log-level DEBUG \
  --p-no-restricted-access \
  --o-paired-reads paired-reads-42.qza \
  --o-single-reads XX_single_reads \
  --o-failed-runs XX_failed_runs

qiime fondue get-sequences \
  --i-accession-ids ncbi-accession-i-ds-43.qza \
  --p- \
  --p-retries 5 \
  --p-n-jobs 1 \
  --p-log-level DEBUG \
  --p-no-restricted-access \
  --o-paired-reads paired-reads-43.qza \
  --o-single-reads XX_single_reads \
  --o-failed-runs XX_failed_runs
```

```
qiime fondue get-sequences \
  --i-accession-ids ncbi-accession-i-ds-44.qza \
  --p- \
  --p-retries 5 \
  --p-n-jobs 1 \
  --p-log-level DEBUG \
  --p-no-restricted-access \
  --o-paired-reads paired-reads-44.qza \
  --o-single-reads XX_single_reads \
  --o-failed-runs XX_failed_runs

qiime fondue get-sequences \
  --i-accession-ids ncbi-accession-i-ds-45.qza \
  --p- \
  --p-retries 5 \
  --p-n-jobs 1 \
  --p-log-level DEBUG \
  --p-no-restricted-access \
  --o-paired-reads paired-reads-45.qza \
  --o-single-reads XX_single_reads \
  --o-failed-runs XX_failed_runs

qiime fondue get-sequences \
  --i-accession-ids ncbi-accession-i-ds-46.qza \
  --p- \
  --p-retries 5 \
  --p-n-jobs 1 \
  --p-log-level DEBUG \
  --p-no-restricted-access \
  --o-paired-reads paired-reads-46.qza \
  --o-single-reads XX_single_reads \
  --o-failed-runs XX_failed_runs

qiime fondue get-sequences \
  --i-accession-ids ncbi-accession-i-ds-47.qza \
  --p- \
  --p-retries 5 \
  --p-n-jobs 1 \
  --p-log-level DEBUG \
  --p-no-restricted-access \
  --o-paired-reads paired-reads-47.qza \
  --o-single-reads XX_single_reads \
  --o-failed-runs XX_failed_runs

qiime fondue get-sequences \
  --i-accession-ids ncbi-accession-i-ds-48.qza \
  --p- \
  --p-retries 5 \
  --p-n-jobs 1 \
  --p-log-level DEBUG \
  --p-no-restricted-access \
  --o-paired-reads paired-reads-48.qza \
```

```
  --o-single-reads XX_single_reads \
  --o-failed-runs XX_failed_runs

qiime fondue get-sequences \
  --i-accession-ids ncbi-accession-i-ds-49.qza \
  --p- \
  --p-retries 5 \
  --p-n-jobs 1 \
  --p-log-level DEBUG \
  --p-no-restricted-access \
  --o-paired-reads paired-reads-49.qza \
  --o-single-reads XX_single_reads \
  --o-failed-runs XX_failed_runs

qiime fondue get-sequences \
  --i-accession-ids ncbi-accession-i-ds-50.qza \
  --p- \
  --p-retries 5 \
  --p-n-jobs 1 \
  --p-log-level DEBUG \
  --p-no-restricted-access \
  --o-paired-reads paired-reads-50.qza \
  --o-single-reads XX_single_reads \
  --o-failed-runs XX_failed_runs

qiime fondue get-sequences \
  --i-accession-ids ncbi-accession-i-ds-51.qza \
  --p- \
  --p-retries 5 \
  --p-n-jobs 1 \
  --p-log-level DEBUG \
  --p-no-restricted-access \
  --o-paired-reads paired-reads-51.qza \
  --o-single-reads XX_single_reads \
  --o-failed-runs XX_failed_runs

qiime fondue get-sequences \
  --i-accession-ids ncbi-accession-i-ds-52.qza \
  --p- \
  --p-retries 5 \
  --p-n-jobs 1 \
  --p-log-level DEBUG \
  --p-no-restricted-access \
  --o-paired-reads paired-reads-52.qza \
  --o-single-reads XX_single_reads \
  --o-failed-runs XX_failed_runs

qiime fondue get-sequences \
  --i-accession-ids ncbi-accession-i-ds-53.qza \
  --p- \
  --p-retries 5 \
  --p-n-jobs 1 \
```

```
  --p-log-level DEBUG \
  --p-no-restricted-access \
  --o-paired-reads paired-reads-53.qza \
  --o-single-reads XX_single_reads \
  --o-failed-runs XX_failed_runs

qiime fondue get-sequences \
  --i-accession-ids ncbi-accession-i-ds-54.qza \
  --p- \
  --p-retries 5 \
  --p-n-jobs 1 \
  --p-log-level DEBUG \
  --p-no-restricted-access \
  --o-paired-reads paired-reads-54.qza \
  --o-single-reads XX_single_reads \
  --o-failed-runs XX_failed_runs

qiime fondue get-sequences \
  --i-accession-ids ncbi-accession-i-ds-55.qza \
  --p- \
  --p-retries 5 \
  --p-n-jobs 1 \
  --p-log-level DEBUG \
  --p-no-restricted-access \
  --o-paired-reads paired-reads-55.qza \
  --o-single-reads XX_single_reads \
  --o-failed-runs XX_failed_runs

qiime fondue get-sequences \
  --i-accession-ids ncbi-accession-i-ds-56.qza \
  --p- \
  --p-retries 5 \
  --p-n-jobs 1 \
  --p-log-level DEBUG \
  --p-no-restricted-access \
  --o-paired-reads paired-reads-56.qza \
  --o-single-reads XX_single_reads \
  --o-failed-runs XX_failed_runs

qiime fondue get-sequences \
  --i-accession-ids ncbi-accession-i-ds-57.qza \
  --p- \
  --p-retries 5 \
  --p-n-jobs 1 \
  --p-log-level DEBUG \
  --p-no-restricted-access \
  --o-paired-reads paired-reads-57.qza \
  --o-single-reads XX_single_reads \
  --o-failed-runs XX_failed_runs

qiime fondue get-sequences \
  --i-accession-ids ncbi-accession-i-ds-58.qza \
```

```
  --p- \
  --p-retries 5 \
  --p-n-jobs 1 \
  --p-log-level DEBUG \
  --p-no-restricted-access \
  --o-paired-reads paired-reads-58.qza \
  --o-single-reads XX_single_reads \
  --o-failed-runs XX_failed_runs

qiime fondue get-sequences \
  --i-accession-ids ncbi-accession-i-ds-59.qza \
  --p- \
  --p-retries 5 \
  --p-n-jobs 1 \
  --p-log-level DEBUG \
  --p-no-restricted-access \
  --o-paired-reads paired-reads-59.qza \
  --o-single-reads XX_single_reads \
  --o-failed-runs XX_failed_runs

qiime fondue get-sequences \
  --i-accession-ids ncbi-accession-i-ds-60.qza \
  --p- \
  --p-retries 5 \
  --p-n-jobs 1 \
  --p-log-level DEBUG \
  --p-no-restricted-access \
  --o-paired-reads paired-reads-60.qza \
  --o-single-reads XX_single_reads \
  --o-failed-runs XX_failed_runs

qiime fondue get-sequences \
  --i-accession-ids ncbi-accession-i-ds-61.qza \
  --p- \
  --p-retries 5 \
  --p-n-jobs 1 \
  --p-log-level DEBUG \
  --p-no-restricted-access \
  --o-paired-reads paired-reads-61.qza \
  --o-single-reads XX_single_reads \
  --o-failed-runs XX_failed_runs

qiime fondue get-sequences \
  --i-accession-ids ncbi-accession-i-ds-62.qza \
  --p- \
  --p-retries 5 \
  --p-n-jobs 1 \
  --p-log-level DEBUG \
  --p-no-restricted-access \
  --o-paired-reads paired-reads-62.qza \
  --o-single-reads XX_single_reads \
  --o-failed-runs XX_failed_runs
```

```
qiime fondue get-sequences \
  --i-accession-ids ncbi-accession-i-ds-63.qza \
  --p- \
  --p-retries 5 \
  --p-n-jobs 1 \
  --p-log-level DEBUG \
  --p-no-restricted-access \
  --o-paired-reads paired-reads-63.qza \
  --o-single-reads XX_single_reads \
  --o-failed-runs XX_failed_runs

qiime fondue get-sequences \
  --i-accession-ids ncbi-accession-i-ds-64.qza \
  --p- \
  --p-retries 5 \
  --p-n-jobs 1 \
  --p-log-level DEBUG \
  --p-no-restricted-access \
  --o-paired-reads paired-reads-64.qza \
  --o-single-reads XX_single_reads \
  --o-failed-runs XX_failed_runs

qiime fondue get-sequences \
  --i-accession-ids ncbi-accession-i-ds-65.qza \
  --p- \
  --p-retries 5 \
  --p-n-jobs 1 \
  --p-log-level DEBUG \
  --p-no-restricted-access \
  --o-paired-reads paired-reads-65.qza \
  --o-single-reads XX_single_reads \
  --o-failed-runs XX_failed_runs

qiime fondue get-sequences \
  --i-accession-ids ncbi-accession-i-ds-66.qza \
  --p- \
  --p-retries 5 \
  --p-n-jobs 1 \
  --p-log-level DEBUG \
  --p-no-restricted-access \
  --o-paired-reads paired-reads-66.qza \
  --o-single-reads XX_single_reads \
  --o-failed-runs XX_failed_runs

qiime fondue get-sequences \
  --i-accession-ids ncbi-accession-i-ds-67.qza \
  --p- \
  --p-retries 5 \
  --p-n-jobs 1 \
  --p-log-level DEBUG \
  --p-no-restricted-access \
```

```
  --o-paired-reads paired-reads-67.qza \
  --o-single-reads XX_single_reads \
  --o-failed-runs XX_failed_runs

qiime fondue get-sequences \
  --i-accession-ids ncbi-accession-i-ds-68.qza \
  --p- \
  --p-retries 5 \
  --p-n-jobs 1 \
  --p-log-level DEBUG \
  --p-no-restricted-access \
  --o-paired-reads paired-reads-68.qza \
  --o-single-reads XX_single_reads \
  --o-failed-runs XX_failed_runs

qiime fondue get-sequences \
  --i-accession-ids ncbi-accession-i-ds-69.qza \
  --p- \
  --p-retries 5 \
  --p-n-jobs 1 \
  --p-log-level DEBUG \
  --p-no-restricted-access \
  --o-paired-reads paired-reads-69.qza \
  --o-single-reads XX_single_reads \
  --o-failed-runs XX_failed_runs

qiime fondue get-sequences \
  --i-accession-ids ncbi-accession-i-ds-70.qza \
  --p- \
  --p-retries 5 \
  --p-n-jobs 1 \
  --p-log-level DEBUG \
  --p-no-restricted-access \
  --o-paired-reads paired-reads-70.qza \
  --o-single-reads XX_single_reads \
  --o-failed-runs XX_failed_runs

qiime fondue get-sequences \
  --i-accession-ids ncbi-accession-i-ds-71.qza \
  --p- \
  --p-retries 5 \
  --p-n-jobs 1 \
  --p-log-level DEBUG \
  --p-no-restricted-access \
  --o-paired-reads paired-reads-71.qza \
  --o-single-reads XX_single_reads \
  --o-failed-runs XX_failed_runs

qiime fondue get-sequences \
  --i-accession-ids ncbi-accession-i-ds-72.qza \
  --p- \
  --p-retries 5 \
```

```
  --p-n-jobs 1 \
  --p-log-level DEBUG \
  --p-no-restricted-access \
  --o-paired-reads paired-reads-72.qza \
  --o-single-reads XX_single_reads \
  --o-failed-runs XX_failed_runs

qiime fondue get-sequences \
  --i-accession-ids ncbi-accession-i-ds-73.qza \
  --p- \
  --p-retries 5 \
  --p-n-jobs 1 \
  --p-log-level DEBUG \
  --p-no-restricted-access \
  --o-paired-reads paired-reads-73.qza \
  --o-single-reads XX_single_reads \
  --o-failed-runs XX_failed_runs

qiime fondue get-sequences \
  --i-accession-ids ncbi-accession-i-ds-74.qza \
  --p- \
  --p-retries 5 \
  --p-n-jobs 1 \
  --p-log-level DEBUG \
  --p-no-restricted-access \
  --o-paired-reads paired-reads-74.qza \
  --o-single-reads XX_single_reads \
  --o-failed-runs XX_failed_runs

qiime fondue get-sequences \
  --i-accession-ids ncbi-accession-i-ds-75.qza \
  --p- \
  --p-retries 5 \
  --p-n-jobs 1 \
  --p-log-level DEBUG \
  --p-no-restricted-access \
  --o-paired-reads paired-reads-75.qza \
  --o-single-reads XX_single_reads \
  --o-failed-runs XX_failed_runs

qiime fondue get-sequences \
  --i-accession-ids ncbi-accession-i-ds-76.qza \
  --p- \
  --p-retries 5 \
  --p-n-jobs 1 \
  --p-log-level DEBUG \
  --p-no-restricted-access \
  --o-paired-reads paired-reads-76.qza \
  --o-single-reads XX_single_reads \
  --o-failed-runs XX_failed_runs

qiime fondue get-sequences \
```

```
  --i-accession-ids ncbi-accession-i-ds-77.qza \
  --p- \
  --p-retries 5 \
  --p-n-jobs 1 \
  --p-log-level DEBUG \
  --p-no-restricted-access \
  --o-paired-reads paired-reads-77.qza \
  --o-single-reads XX_single_reads \
  --o-failed-runs XX_failed_runs

qiime fondue get-sequences \
  --i-accession-ids ncbi-accession-i-ds-78.qza \
  --p- \
  --p-retries 5 \
  --p-n-jobs 1 \
  --p-log-level DEBUG \
  --p-no-restricted-access \
  --o-paired-reads paired-reads-78.qza \
  --o-single-reads XX_single_reads \
  --o-failed-runs XX_failed_runs

qiime fondue get-sequences \
  --i-accession-ids ncbi-accession-i-ds-79.qza \
  --p- \
  --p-retries 5 \
  --p-n-jobs 1 \
  --p-log-level DEBUG \
  --p-no-restricted-access \
  --o-paired-reads paired-reads-79.qza \
  --o-single-reads XX_single_reads \
  --o-failed-runs XX_failed_runs

qiime fondue get-sequences \
  --i-accession-ids ncbi-accession-i-ds-80.qza \
  --p- \
  --p-retries 5 \
  --p-n-jobs 1 \
  --p-log-level DEBUG \
  --p-no-restricted-access \
  --o-paired-reads paired-reads-80.qza \
  --o-single-reads XX_single_reads \
  --o-failed-runs XX_failed_runs

qiime fondue get-sequences \
  --i-accession-ids ncbi-accession-i-ds-81.qza \
  --p- \
  --p-retries 5 \
  --p-n-jobs 1 \
  --p-log-level DEBUG \
  --p-no-restricted-access \
  --o-paired-reads paired-reads-81.qza \
  --o-single-reads XX_single_reads \
```

```
  --o-failed-runs XX_failed_runs

qiime fondue get-sequences \
  --i-accession-ids ncbi-accession-i-ds-82.qza \
  --p- \
  --p-retries 5 \
  --p-n-jobs 1 \
  --p-log-level DEBUG \
  --p-no-restricted-access \
  --o-paired-reads paired-reads-82.qza \
  --o-single-reads XX_single_reads \
  --o-failed-runs XX_failed_runs

qiime fondue get-sequences \
  --i-accession-ids ncbi-accession-i-ds-83.qza \
  --p- \
  --p-retries 5 \
  --p-n-jobs 1 \
  --p-log-level DEBUG \
  --p-no-restricted-access \
  --o-paired-reads paired-reads-83.qza \
  --o-single-reads XX_single_reads \
  --o-failed-runs XX_failed_runs

qiime fondue get-sequences \
  --i-accession-ids ncbi-accession-i-ds-84.qza \
  --p- \
  --p-retries 5 \
  --p-n-jobs 1 \
  --p-log-level DEBUG \
  --p-no-restricted-access \
  --o-paired-reads paired-reads-84.qza \
  --o-single-reads XX_single_reads \
  --o-failed-runs XX_failed_runs

qiime fondue get-sequences \
  --i-accession-ids ncbi-accession-i-ds-85.qza \
  --p- \
  --p-retries 5 \
  --p-n-jobs 1 \
  --p-log-level DEBUG \
  --p-no-restricted-access \
  --o-paired-reads paired-reads-85.qza \
  --o-single-reads XX_single_reads \
  --o-failed-runs XX_failed_runs

qiime fondue get-sequences \
  --i-accession-ids ncbi-accession-i-ds-86.qza \
  --p- \
  --p-retries 5 \
  --p-n-jobs 1 \
  --p-log-level DEBUG \
```

```
  --p-no-restricted-access \
  --o-paired-reads paired-reads-86.qza \
  --o-single-reads XX_single_reads \
  --o-failed-runs XX_failed_runs

qiime fondue get-sequences \
  --i-accession-ids ncbi-accession-i-ds-87.qza \
  --p- \
  --p-retries 5 \
  --p-n-jobs 1 \
  --p-log-level DEBUG \
  --p-no-restricted-access \
  --o-paired-reads paired-reads-87.qza \
  --o-single-reads XX_single_reads \
  --o-failed-runs XX_failed_runs

qiime fondue get-sequences \
  --i-accession-ids ncbi-accession-i-ds-88.qza \
  --p- \
  --p-retries 5 \
  --p-n-jobs 1 \
  --p-log-level DEBUG \
  --p-no-restricted-access \
  --o-paired-reads paired-reads-88.qza \
  --o-single-reads XX_single_reads \
  --o-failed-runs XX_failed_runs

qiime fondue get-sequences \
  --i-accession-ids ncbi-accession-i-ds-89.qza \
  --p- \
  --p-retries 5 \
  --p-n-jobs 1 \
  --p-log-level DEBUG \
  --p-no-restricted-access \
  --o-paired-reads paired-reads-89.qza \
  --o-single-reads XX_single_reads \
  --o-failed-runs XX_failed_runs

qiime fondue get-sequences \
  --i-accession-ids ncbi-accession-i-ds-90.qza \
  --p- \
  --p-retries 5 \
  --p-n-jobs 1 \
  --p-log-level DEBUG \
  --p-no-restricted-access \
  --o-paired-reads paired-reads-90.qza \
  --o-single-reads XX_single_reads \
  --o-failed-runs XX_failed_runs

qiime fondue get-sequences \
  --i-accession-ids ncbi-accession-i-ds-91.qza \
  --p- \
```

```
  --p-retries 5 \
  --p-n-jobs 1 \
  --p-log-level DEBUG \
  --p-no-restricted-access \
  --o-paired-reads paired-reads-91.qza \
  --o-single-reads XX_single_reads \
  --o-failed-runs XX_failed_runs

qiime fondue get-sequences \
  --i-accession-ids ncbi-accession-i-ds-92.qza \
  --p- \
  --p-retries 5 \
  --p-n-jobs 1 \
  --p-log-level DEBUG \
  --p-no-restricted-access \
  --o-paired-reads paired-reads-92.qza \
  --o-single-reads XX_single_reads \
  --o-failed-runs XX_failed_runs

qiime fondue get-sequences \
  --i-accession-ids ncbi-accession-i-ds-93.qza \
  --p- \
  --p-retries 5 \
  --p-n-jobs 1 \
  --p-log-level DEBUG \
  --p-no-restricted-access \
  --o-paired-reads paired-reads-93.qza \
  --o-single-reads XX_single_reads \
  --o-failed-runs XX_failed_runs

qiime fondue get-sequences \
  --i-accession-ids ncbi-accession-i-ds-94.qza \
  --p- \
  --p-retries 5 \
  --p-n-jobs 1 \
  --p-log-level DEBUG \
  --p-no-restricted-access \
  --o-paired-reads paired-reads-94.qza \
  --o-single-reads XX_single_reads \
  --o-failed-runs XX_failed_runs

qiime fondue get-sequences \
  --i-accession-ids ncbi-accession-i-ds-95.qza \
  --p- \
  --p-retries 5 \
  --p-n-jobs 1 \
  --p-log-level DEBUG \
  --p-no-restricted-access \
  --o-paired-reads paired-reads-95.qza \
  --o-single-reads XX_single_reads \
  --o-failed-runs XX_failed_runs
```

```
qiime fondue get-sequences \
  --i-accession-ids ncbi-accession-i-ds-96.qza \
  --p- \
  --p-retries 5 \
  --p-n-jobs 1 \
  --p-log-level DEBUG \
  --p-no-restricted-access \
  --o-paired-reads paired-reads-96.qza \
  --o-single-reads XX_single_reads \
  --o-failed-runs XX_failed_runs

qiime fondue get-sequences \
  --i-accession-ids ncbi-accession-i-ds-97.qza \
  --p- \
  --p-retries 5 \
  --p-n-jobs 1 \
  --p-log-level DEBUG \
  --p-no-restricted-access \
  --o-paired-reads paired-reads-97.qza \
  --o-single-reads XX_single_reads \
  --o-failed-runs XX_failed_runs

qiime fondue get-sequences \
  --i-accession-ids ncbi-accession-i-ds-98.qza \
  --p- \
  --p-retries 5 \
  --p-n-jobs 1 \
  --p-log-level DEBUG \
  --p-no-restricted-access \
  --o-paired-reads paired-reads-98.qza \
  --o-single-reads XX_single_reads \
  --o-failed-runs XX_failed_runs

qiime fondue get-sequences \
  --i-accession-ids ncbi-accession-i-ds-99.qza \
  --p- \
  --p-retries 5 \
  --p-n-jobs 1 \
  --p-log-level DEBUG \
  --p-no-restricted-access \
  --o-paired-reads paired-reads-99.qza \
  --o-single-reads XX_single_reads \
  --o-failed-runs XX_failed_runs

qiime fondue get-sequences \
  --i-accession-ids ncbi-accession-i-ds-100.qza \
  --p- \
  --p-retries 5 \
  --p-n-jobs 1 \
  --p-log-level DEBUG \
  --p-no-restricted-access \
  --o-paired-reads paired-reads-100.qza \
```

```
 --o-single-reads XX_single_reads \
 --o-failed-runs XX_failed_runs

qiime fondue get-sequences \
 --i-accession-ids ncbi-accession-i-ds-101.qza \
 --p- \
 --p-retries 5 \
 --p-n-jobs 1 \
 --p-log-level DEBUG \
 --p-no-restricted-access \
 --o-paired-reads paired-reads-101.qza \
 --o-single-reads XX_single_reads \
 --o-failed-runs XX_failed_runs

qiime fondue get-sequences \
 --i-accession-ids ncbi-accession-i-ds-102.qza \
 --p- \
 --p-retries 5 \
 --p-n-jobs 1 \
 --p-log-level DEBUG \
 --p-no-restricted-access \
 --o-paired-reads paired-reads-102.qza \
 --o-single-reads XX_single_reads \
 --o-failed-runs XX_failed_runs

qiime fondue get-sequences \
 --i-accession-ids ncbi-accession-i-ds-103.qza \
 --p- \
 --p-retries 5 \
 --p-n-jobs 1 \
 --p-log-level DEBUG \
 --p-no-restricted-access \
 --o-paired-reads paired-reads-103.qza \
 --o-single-reads XX_single_reads \
 --o-failed-runs XX_failed_runs

qiime fondue get-sequences \
 --i-accession-ids ncbi-accession-i-ds-104.qza \
 --p- \
 --p-retries 5 \
 --p-n-jobs 1 \
 --p-log-level DEBUG \
 --p-no-restricted-access \
 --o-paired-reads paired-reads-104.qza \
 --o-single-reads XX_single_reads \
 --o-failed-runs XX_failed_runs

qiime fondue get-sequences \
 --i-accession-ids ncbi-accession-i-ds-105.qza \
 --p- \
 --p-retries 5 \
 --p-n-jobs 1 \
```

```
  --p-log-level DEBUG \
  --p-no-restricted-access \
  --o-paired-reads paired-reads-105.qza \
  --o-single-reads XX_single_reads \
  --o-failed-runs XX_failed_runs

qiime fondue get-sequences \
  --i-accession-ids ncbi-accession-i-ds-106.qza \
  --p- \
  --p-retries 5 \
  --p-n-jobs 1 \
  --p-log-level DEBUG \
  --p-no-restricted-access \
  --o-paired-reads paired-reads-106.qza \
  --o-single-reads XX_single_reads \
  --o-failed-runs XX_failed_runs

qiime fondue get-sequences \
  --i-accession-ids ncbi-accession-i-ds-107.qza \
  --p- \
  --p-retries 5 \
  --p-n-jobs 1 \
  --p-log-level DEBUG \
  --p-no-restricted-access \
  --o-paired-reads paired-reads-107.qza \
  --o-single-reads XX_single_reads \
  --o-failed-runs XX_failed_runs

qiime fondue get-sequences \
  --i-accession-ids ncbi-accession-i-ds-108.qza \
  --p- \
  --p-retries 5 \
  --p-n-jobs 1 \
  --p-log-level DEBUG \
  --p-no-restricted-access \
  --o-paired-reads paired-reads-108.qza \
  --o-single-reads XX_single_reads \
  --o-failed-runs XX_failed_runs

qiime fondue get-sequences \
  --i-accession-ids ncbi-accession-i-ds-109.qza \
  --p- \
  --p-retries 5 \
  --p-n-jobs 1 \
  --p-log-level DEBUG \
  --p-no-restricted-access \
  --o-paired-reads paired-reads-109.qza \
  --o-single-reads XX_single_reads \
  --o-failed-runs XX_failed_runs

qiime fondue get-sequences \
  --i-accession-ids ncbi-accession-i-ds-110.qza \
```

```
  --p- \
  --p-retries 5 \
  --p-n-jobs 1 \
  --p-log-level DEBUG \
  --p-no-restricted-access \
  --o-paired-reads paired-reads-110.qza \
  --o-single-reads XX_single_reads \
  --o-failed-runs XX_failed_runs

qiime fondue get-sequences \
  --i-accession-ids ncbi-accession-i-ds-111.qza \
  --p- \
  --p-retries 5 \
  --p-n-jobs 1 \
  --p-log-level DEBUG \
  --p-no-restricted-access \
  --o-paired-reads paired-reads-111.qza \
  --o-single-reads XX_single_reads \
  --o-failed-runs XX_failed_runs

qiime fondue get-sequences \
  --i-accession-ids ncbi-accession-i-ds-112.qza \
  --p- \
  --p-retries 5 \
  --p-n-jobs 1 \
  --p-log-level DEBUG \
  --p-no-restricted-access \
  --o-paired-reads paired-reads-112.qza \
  --o-single-reads XX_single_reads \
  --o-failed-runs XX_failed_runs

qiime fondue get-sequences \
  --i-accession-ids ncbi-accession-i-ds-113.qza \
  --p- \
  --p-retries 5 \
  --p-n-jobs 1 \
  --p-log-level DEBUG \
  --p-no-restricted-access \
  --o-paired-reads paired-reads-113.qza \
  --o-single-reads XX_single_reads \
  --o-failed-runs XX_failed_runs

qiime fondue get-sequences \
  --i-accession-ids ncbi-accession-i-ds-114.qza \
  --p- \
  --p-retries 5 \
  --p-n-jobs 1 \
  --p-log-level DEBUG \
  --p-no-restricted-access \
  --o-paired-reads paired-reads-114.qza \
  --o-single-reads XX_single_reads \
  --o-failed-runs XX_failed_runs
```

```
qiime fondue get-sequences \
  --i-accession-ids ncbi-accession-i-ds-115.qza \
  --p- \
  --p-retries 5 \
  --p-n-jobs 1 \
  --p-log-level DEBUG \
  --p-no-restricted-access \
  --o-paired-reads paired-reads-115.qza \
  --o-single-reads XX_single_reads \
  --o-failed-runs XX_failed_runs

qiime fondue get-sequences \
  --i-accession-ids ncbi-accession-i-ds-116.qza \
  --p- \
  --p-retries 5 \
  --p-n-jobs 1 \
  --p-log-level DEBUG \
  --p-no-restricted-access \
  --o-paired-reads paired-reads-116.qza \
  --o-single-reads XX_single_reads \
  --o-failed-runs XX_failed_runs

qiime fondue get-sequences \
  --i-accession-ids ncbi-accession-i-ds-117.qza \
  --p- \
  --p-retries 5 \
  --p-n-jobs 1 \
  --p-log-level DEBUG \
  --p-no-restricted-access \
  --o-paired-reads paired-reads-117.qza \
  --o-single-reads XX_single_reads \
  --o-failed-runs XX_failed_runs

qiime fondue get-sequences \
  --i-accession-ids ncbi-accession-i-ds-118.qza \
  --p- \
  --p-retries 5 \
  --p-n-jobs 1 \
  --p-log-level DEBUG \
  --p-no-restricted-access \
  --o-paired-reads paired-reads-118.qza \
  --o-single-reads XX_single_reads \
  --o-failed-runs XX_failed_runs

qiime fondue get-sequences \
  --i-accession-ids ncbi-accession-i-ds-119.qza \
  --p- \
  --p-retries 5 \
  --p-n-jobs 1 \
  --p-log-level DEBUG \
  --p-no-restricted-access \
```

```
  --o-paired-reads paired-reads-119.qza \
  --o-single-reads XX_single_reads \
  --o-failed-runs XX_failed_runs

qiime fondue get-sequences \
  --i-accession-ids ncbi-accession-i-ds-120.qza \
  --p- \
  --p-retries 5 \
  --p-n-jobs 1 \
  --p-log-level DEBUG \
  --p-no-restricted-access \
  --o-paired-reads paired-reads-120.qza \
  --o-single-reads XX_single_reads \
  --o-failed-runs XX_failed_runs

qiime fondue get-sequences \
  --i-accession-ids ncbi-accession-i-ds-121.qza \
  --p- \
  --p-retries 5 \
  --p-n-jobs 1 \
  --p-log-level DEBUG \
  --p-no-restricted-access \
  --o-paired-reads paired-reads-121.qza \
  --o-single-reads XX_single_reads \
  --o-failed-runs XX_failed_runs

qiime fondue get-sequences \
  --i-accession-ids ncbi-accession-i-ds-122.qza \
  --p- \
  --p-retries 5 \
  --p-n-jobs 1 \
  --p-log-level DEBUG \
  --p-no-restricted-access \
  --o-paired-reads paired-reads-122.qza \
  --o-single-reads XX_single_reads \
  --o-failed-runs XX_failed_runs

qiime fondue get-sequences \
  --i-accession-ids ncbi-accession-i-ds-123.qza \
  --p- \
  --p-retries 5 \
  --p-n-jobs 1 \
  --p-log-level DEBUG \
  --p-no-restricted-access \
  --o-paired-reads paired-reads-123.qza \
  --o-single-reads XX_single_reads \
  --o-failed-runs XX_failed_runs

qiime fondue get-sequences \
  --i-accession-ids ncbi-accession-i-ds-124.qza \
  --p- \
  --p-retries 5 \
```

```
  --p-n-jobs 1 \
  --p-log-level DEBUG \
  --p-no-restricted-access \
  --o-paired-reads paired-reads-124.qza \
  --o-single-reads XX_single_reads \
  --o-failed-runs XX_failed_runs

qiime fondue get-sequences \
  --i-accession-ids ncbi-accession-i-ds-125.qza \
  --p- \
  --p-retries 5 \
  --p-n-jobs 1 \
  --p-log-level DEBUG \
  --p-no-restricted-access \
  --o-paired-reads paired-reads-125.qza \
  --o-single-reads XX_single_reads \
  --o-failed-runs XX_failed_runs

qiime fondue get-sequences \
  --i-accession-ids ncbi-accession-i-ds-126.qza \
  --p- \
  --p-retries 5 \
  --p-n-jobs 1 \
  --p-log-level DEBUG \
  --p-no-restricted-access \
  --o-paired-reads paired-reads-126.qza \
  --o-single-reads XX_single_reads \
  --o-failed-runs XX_failed_runs

qiime fondue get-sequences \
  --i-accession-ids ncbi-accession-i-ds-127.qza \
  --p- \
  --p-retries 5 \
  --p-n-jobs 1 \
  --p-log-level DEBUG \
  --p-no-restricted-access \
  --o-paired-reads paired-reads-127.qza \
  --o-single-reads XX_single_reads \
  --o-failed-runs XX_failed_runs

qiime fondue get-sequences \
  --i-accession-ids ncbi-accession-i-ds-128.qza \
  --p- \
  --p-retries 5 \
  --p-n-jobs 1 \
  --p-log-level DEBUG \
  --p-no-restricted-access \
  --o-paired-reads paired-reads-128.qza \
  --o-single-reads XX_single_reads \
  --o-failed-runs XX_failed_runs

qiime fondue get-sequences \
```

```
  --i-accession-ids ncbi-accession-i-ds-129.qza \
  --p- \
  --p-retries 5 \
  --p-n-jobs 1 \
  --p-log-level DEBUG \
  --p-no-restricted-access \
  --o-paired-reads paired-reads-129.qza \
  --o-single-reads XX_single_reads \
  --o-failed-runs XX_failed_runs

qiime fondue get-sequences \
  --i-accession-ids ncbi-accession-i-ds-130.qza \
  --p- \
  --p-retries 5 \
  --p-n-jobs 1 \
  --p-log-level DEBUG \
  --p-no-restricted-access \
  --o-paired-reads paired-reads-130.qza \
  --o-single-reads XX_single_reads \
  --o-failed-runs XX_failed_runs

qiime fondue get-sequences \
  --i-accession-ids ncbi-accession-i-ds-131.qza \
  --p- \
  --p-retries 5 \
  --p-n-jobs 1 \
  --p-log-level DEBUG \
  --p-no-restricted-access \
  --o-paired-reads paired-reads-131.qza \
  --o-single-reads XX_single_reads \
  --o-failed-runs XX_failed_runs

qiime fondue get-sequences \
  --i-accession-ids ncbi-accession-i-ds-132.qza \
  --p- \
  --p-retries 5 \
  --p-n-jobs 1 \
  --p-log-level DEBUG \
  --p-no-restricted-access \
  --o-paired-reads paired-reads-132.qza \
  --o-single-reads XX_single_reads \
  --o-failed-runs XX_failed_runs

qiime fondue get-sequences \
  --i-accession-ids ncbi-accession-i-ds-133.qza \
  --p- \
  --p-retries 5 \
  --p-n-jobs 1 \
  --p-log-level DEBUG \
  --p-no-restricted-access \
  --o-paired-reads paired-reads-133.qza \
  --o-single-reads XX_single_reads \
```

```
  --o-failed-runs XX_failed_runs

qiime fondue get-sequences \
  --i-accession-ids ncbi-accession-i-ds-134.qza \
  --p- \
  --p-retries 5 \
  --p-n-jobs 1 \
  --p-log-level DEBUG \
  --p-no-restricted-access \
  --o-paired-reads paired-reads-134.qza \
  --o-single-reads XX_single_reads \
  --o-failed-runs XX_failed_runs

qiime fondue get-sequences \
  --i-accession-ids ncbi-accession-i-ds-135.qza \
  --p- \
  --p-retries 5 \
  --p-n-jobs 1 \
  --p-log-level DEBUG \
  --p-no-restricted-access \
  --o-paired-reads paired-reads-135.qza \
  --o-single-reads XX_single_reads \
  --o-failed-runs XX_failed_runs

qiime fondue get-sequences \
  --i-accession-ids ncbi-accession-i-ds-136.qza \
  --p- \
  --p-retries 5 \
  --p-n-jobs 1 \
  --p-log-level DEBUG \
  --p-no-restricted-access \
  --o-paired-reads paired-reads-136.qza \
  --o-single-reads XX_single_reads \
  --o-failed-runs XX_failed_runs

qiime fondue get-sequences \
  --i-accession-ids ncbi-accession-i-ds-137.qza \
  --p- \
  --p-retries 5 \
  --p-n-jobs 1 \
  --p-log-level DEBUG \
  --p-no-restricted-access \
  --o-paired-reads paired-reads-137.qza \
  --o-single-reads XX_single_reads \
  --o-failed-runs XX_failed_runs

qiime fondue get-sequences \
  --i-accession-ids ncbi-accession-i-ds-138.qza \
  --p- \
  --p-retries 5 \
  --p-n-jobs 1 \
  --p-log-level DEBUG \
```

```
  --p-no-restricted-access \
  --o-paired-reads paired-reads-138.qza \
  --o-single-reads XX_single_reads \
  --o-failed-runs XX_failed_runs

qiime fondue get-sequences \
  --i-accession-ids ncbi-accession-i-ds-139.qza \
  --p- \
  --p-retries 5 \
  --p-n-jobs 1 \
  --p-log-level DEBUG \
  --p-no-restricted-access \
  --o-paired-reads paired-reads-139.qza \
  --o-single-reads XX_single_reads \
  --o-failed-runs XX_failed_runs

qiime fondue get-sequences \
  --i-accession-ids ncbi-accession-i-ds-140.qza \
  --p- \
  --p-retries 5 \
  --p-n-jobs 1 \
  --p-log-level DEBUG \
  --p-no-restricted-access \
  --o-paired-reads paired-reads-140.qza \
  --o-single-reads XX_single_reads \
  --o-failed-runs XX_failed_runs

qiime fondue get-sequences \
  --i-accession-ids ncbi-accession-i-ds-141.qza \
  --p- \
  --p-retries 5 \
  --p-n-jobs 1 \
  --p-log-level DEBUG \
  --p-no-restricted-access \
  --o-paired-reads paired-reads-141.qza \
  --o-single-reads XX_single_reads \
  --o-failed-runs XX_failed_runs

qiime fondue get-sequences \
  --i-accession-ids ncbi-accession-i-ds-142.qza \
  --p- \
  --p-retries 5 \
  --p-n-jobs 1 \
  --p-log-level DEBUG \
  --p-no-restricted-access \
  --o-paired-reads paired-reads-142.qza \
  --o-single-reads XX_single_reads \
  --o-failed-runs XX_failed_runs

qiime fondue get-sequences \
  --i-accession-ids ncbi-accession-i-ds-143.qza \
  --p- \
```

```
  --p-retries 5 \
  --p-n-jobs 1 \
  --p-log-level DEBUG \
  --p-no-restricted-access \
  --o-paired-reads paired-reads-143.qza \
  --o-single-reads XX_single_reads \
  --o-failed-runs XX_failed_runs

qiime fondue get-sequences \
  --i-accession-ids ncbi-accession-i-ds-144.qza \
  --p- \
  --p-retries 5 \
  --p-n-jobs 1 \
  --p-log-level DEBUG \
  --p-no-restricted-access \
  --o-paired-reads paired-reads-144.qza \
  --o-single-reads XX_single_reads \
  --o-failed-runs XX_failed_runs

qiime fondue get-sequences \
  --i-accession-ids ncbi-accession-i-ds-145.qza \
  --p- \
  --p-retries 5 \
  --p-n-jobs 1 \
  --p-log-level DEBUG \
  --p-no-restricted-access \
  --o-paired-reads paired-reads-145.qza \
  --o-single-reads XX_single_reads \
  --o-failed-runs XX_failed_runs

qiime fondue get-sequences \
  --i-accession-ids ncbi-accession-i-ds-146.qza \
  --p- \
  --p-retries 5 \
  --p-n-jobs 1 \
  --p-log-level DEBUG \
  --p-no-restricted-access \
  --o-paired-reads paired-reads-146.qza \
  --o-single-reads XX_single_reads \
  --o-failed-runs XX_failed_runs

qiime fondue get-sequences \
  --i-accession-ids ncbi-accession-i-ds-147.qza \
  --p- \
  --p-retries 5 \
  --p-n-jobs 1 \
  --p-log-level DEBUG \
  --p-no-restricted-access \
  --o-paired-reads paired-reads-147.qza \
  --o-single-reads XX_single_reads \
  --o-failed-runs XX_failed_runs
```

```
qiime fondue get-sequences \
  --i-accession-ids ncbi-accession-i-ds-148.qza \
  --p- \
  --p-retries 5 \
  --p-n-jobs 1 \
  --p-log-level DEBUG \
  --p-no-restricted-access \
  --o-paired-reads paired-reads-148.qza \
  --o-single-reads XX_single_reads \
  --o-failed-runs XX_failed_runs

qiime fondue get-sequences \
  --i-accession-ids ncbi-accession-i-ds-149.qza \
  --p- \
  --p-retries 5 \
  --p-n-jobs 1 \
  --p-log-level DEBUG \
  --p-no-restricted-access \
  --o-paired-reads paired-reads-149.qza \
  --o-single-reads XX_single_reads \
  --o-failed-runs XX_failed_runs

qiime fondue get-sequences \
  --i-accession-ids ncbi-accession-i-ds-150.qza \
  --p- \
  --p-retries 5 \
  --p-n-jobs 1 \
  --p-log-level DEBUG \
  --p-no-restricted-access \
  --o-paired-reads paired-reads-150.qza \
  --o-single-reads XX_single_reads \
  --o-failed-runs XX_failed_runs

qiime fondue get-sequences \
  --i-accession-ids ncbi-accession-i-ds-151.qza \
  --p- \
  --p-retries 5 \
  --p-n-jobs 1 \
  --p-log-level DEBUG \
  --p-no-restricted-access \
  --o-paired-reads paired-reads-151.qza \
  --o-single-reads XX_single_reads \
  --o-failed-runs XX_failed_runs

qiime fondue get-sequences \
  --i-accession-ids ncbi-accession-i-ds-152.qza \
  --p- \
  --p-retries 5 \
  --p-n-jobs 1 \
  --p-log-level DEBUG \
  --p-no-restricted-access \
  --o-paired-reads paired-reads-152.qza \
```

```
  --o-single-reads XX_single_reads \
  --o-failed-runs XX_failed_runs

qiime fondue get-sequences \
  --i-accession-ids ncbi-accession-i-ds-153.qza \
  --p- \
  --p-retries 5 \
  --p-n-jobs 1 \
  --p-log-level DEBUG \
  --p-no-restricted-access \
  --o-paired-reads paired-reads-153.qza \
  --o-single-reads XX_single_reads \
  --o-failed-runs XX_failed_runs

qiime fondue get-sequences \
  --i-accession-ids ncbi-accession-i-ds-154.qza \
  --p- \
  --p-retries 5 \
  --p-n-jobs 1 \
  --p-log-level DEBUG \
  --p-no-restricted-access \
  --o-paired-reads paired-reads-154.qza \
  --o-single-reads XX_single_reads \
  --o-failed-runs XX_failed_runs

qiime fondue get-sequences \
  --i-accession-ids ncbi-accession-i-ds-155.qza \
  --p- \
  --p-retries 5 \
  --p-n-jobs 1 \
  --p-log-level DEBUG \
  --p-no-restricted-access \
  --o-paired-reads paired-reads-155.qza \
  --o-single-reads XX_single_reads \
  --o-failed-runs XX_failed_runs

qiime fondue get-sequences \
  --i-accession-ids ncbi-accession-i-ds-156.qza \
  --p- \
  --p-retries 5 \
  --p-n-jobs 1 \
  --p-log-level DEBUG \
  --p-no-restricted-access \
  --o-paired-reads paired-reads-156.qza \
  --o-single-reads XX_single_reads \
  --o-failed-runs XX_failed_runs

qiime fondue get-sequences \
  --i-accession-ids ncbi-accession-i-ds-157.qza \
  --p- \
  --p-retries 5 \
  --p-n-jobs 1 \
```

```
  --p-log-level DEBUG \
  --p-no-restricted-access \
  --o-paired-reads paired-reads-157.qza \
  --o-single-reads XX_single_reads \
  --o-failed-runs XX_failed_runs

qiime fondue get-sequences \
  --i-accession-ids ncbi-accession-i-ds-158.qza \
  --p- \
  --p-retries 5 \
  --p-n-jobs 1 \
  --p-log-level DEBUG \
  --p-no-restricted-access \
  --o-paired-reads paired-reads-158.qza \
  --o-single-reads XX_single_reads \
  --o-failed-runs XX_failed_runs

qiime fondue get-sequences \
  --i-accession-ids ncbi-accession-i-ds-159.qza \
  --p- \
  --p-retries 5 \
  --p-n-jobs 1 \
  --p-log-level DEBUG \
  --p-no-restricted-access \
  --o-paired-reads paired-reads-159.qza \
  --o-single-reads XX_single_reads \
  --o-failed-runs XX_failed_runs

qiime fondue get-sequences \
  --i-accession-ids ncbi-accession-i-ds-160.qza \
  --p- \
  --p-retries 5 \
  --p-n-jobs 1 \
  --p-log-level DEBUG \
  --p-no-restricted-access \
  --o-paired-reads paired-reads-160.qza \
  --o-single-reads XX_single_reads \
  --o-failed-runs XX_failed_runs

qiime fondue get-sequences \
  --i-accession-ids ncbi-accession-i-ds-161.qza \
  --p- \
  --p-retries 5 \
  --p-n-jobs 1 \
  --p-log-level DEBUG \
  --p-no-restricted-access \
  --o-paired-reads paired-reads-161.qza \
  --o-single-reads XX_single_reads \
  --o-failed-runs XX_failed_runs

qiime fondue get-sequences \
  --i-accession-ids ncbi-accession-i-ds-162.qza \
```

```
  --p- \
  --p-retries 5 \
  --p-n-jobs 1 \
  --p-log-level DEBUG \
  --p-no-restricted-access \
  --o-paired-reads paired-reads-162.qza \
  --o-single-reads XX_single_reads \
  --o-failed-runs XX_failed_runs

qiime fondue get-sequences \
  --i-accession-ids ncbi-accession-i-ds-163.qza \
  --p- \
  --p-retries 5 \
  --p-n-jobs 1 \
  --p-log-level DEBUG \
  --p-no-restricted-access \
  --o-paired-reads paired-reads-163.qza \
  --o-single-reads XX_single_reads \
  --o-failed-runs XX_failed_runs

qiime fondue get-sequences \
  --i-accession-ids ncbi-accession-i-ds-164.qza \
  --p- \
  --p-retries 5 \
  --p-n-jobs 1 \
  --p-log-level DEBUG \
  --p-no-restricted-access \
  --o-paired-reads paired-reads-164.qza \
  --o-single-reads XX_single_reads \
  --o-failed-runs XX_failed_runs

qiime fondue get-sequences \
  --i-accession-ids ncbi-accession-i-ds-165.qza \
  --p- \
  --p-retries 5 \
  --p-n-jobs 1 \
  --p-log-level DEBUG \
  --p-no-restricted-access \
  --o-paired-reads paired-reads-165.qza \
  --o-single-reads XX_single_reads \
  --o-failed-runs XX_failed_runs

qiime fondue get-sequences \
  --i-accession-ids ncbi-accession-i-ds-166.qza \
  --p- \
  --p-retries 5 \
  --p-n-jobs 1 \
  --p-log-level DEBUG \
  --p-no-restricted-access \
  --o-paired-reads paired-reads-166.qza \
  --o-single-reads XX_single_reads \
  --o-failed-runs XX_failed_runs
```

```
qiime fondue get-sequences \
  --i-accession-ids ncbi-accession-i-ds-167.qza \
  --p- \
  --p-retries 5 \
  --p-n-jobs 1 \
  --p-log-level DEBUG \
  --p-no-restricted-access \
  --o-paired-reads paired-reads-167.qza \
  --o-single-reads XX_single_reads \
  --o-failed-runs XX_failed_runs

qiime fondue get-sequences \
  --i-accession-ids ncbi-accession-i-ds-168.qza \
  --p- \
  --p-retries 5 \
  --p-n-jobs 1 \
  --p-log-level DEBUG \
  --p-no-restricted-access \
  --o-paired-reads paired-reads-168.qza \
  --o-single-reads XX_single_reads \
  --o-failed-runs XX_failed_runs

qiime fondue get-sequences \
  --i-accession-ids ncbi-accession-i-ds-169.qza \
  --p- \
  --p-retries 5 \
  --p-n-jobs 1 \
  --p-log-level DEBUG \
  --p-no-restricted-access \
  --o-paired-reads paired-reads-169.qza \
  --o-single-reads XX_single_reads \
  --o-failed-runs XX_failed_runs

qiime fondue get-sequences \
  --i-accession-ids ncbi-accession-i-ds-170.qza \
  --p- \
  --p-retries 5 \
  --p-n-jobs 1 \
  --p-log-level DEBUG \
  --p-no-restricted-access \
  --o-paired-reads paired-reads-170.qza \
  --o-single-reads XX_single_reads \
  --o-failed-runs XX_failed_runs

qiime fondue get-sequences \
  --i-accession-ids ncbi-accession-i-ds-171.qza \
  --p- \
  --p-retries 5 \
  --p-n-jobs 1 \
  --p-log-level DEBUG \
  --p-no-restricted-access \
```

```
  --o-paired-reads paired-reads-171.qza \
  --o-single-reads XX_single_reads \
  --o-failed-runs XX_failed_runs

qiime fondue get-sequences \
  --i-accession-ids ncbi-accession-i-ds-172.qza \
  --p- \
  --p-retries 5 \
  --p-n-jobs 1 \
  --p-log-level DEBUG \
  --p-no-restricted-access \
  --o-paired-reads paired-reads-172.qza \
  --o-single-reads XX_single_reads \
  --o-failed-runs XX_failed_runs

qiime fondue get-sequences \
  --i-accession-ids ncbi-accession-i-ds-173.qza \
  --p- \
  --p-retries 5 \
  --p-n-jobs 1 \
  --p-log-level DEBUG \
  --p-no-restricted-access \
  --o-paired-reads paired-reads-173.qza \
  --o-single-reads XX_single_reads \
  --o-failed-runs XX_failed_runs

qiime fondue get-sequences \
  --i-accession-ids ncbi-accession-i-ds-174.qza \
  --p- \
  --p-retries 5 \
  --p-n-jobs 1 \
  --p-log-level DEBUG \
  --p-no-restricted-access \
  --o-paired-reads paired-reads-174.qza \
  --o-single-reads XX_single_reads \
  --o-failed-runs XX_failed_runs

qiime fondue get-sequences \
  --i-accession-ids ncbi-accession-i-ds-175.qza \
  --p- \
  --p-retries 5 \
  --p-n-jobs 1 \
  --p-log-level DEBUG \
  --p-no-restricted-access \
  --o-paired-reads paired-reads-175.qza \
  --o-single-reads XX_single_reads \
  --o-failed-runs XX_failed_runs

qiime fondue get-sequences \
  --i-accession-ids ncbi-accession-i-ds-176.qza \
  --p- \
  --p-retries 5 \
```

```
  --p-n-jobs 1 \
  --p-log-level DEBUG \
  --p-no-restricted-access \
  --o-paired-reads paired-reads-176.qza \
  --o-single-reads XX_single_reads \
  --o-failed-runs XX_failed_runs

qiime fondue get-sequences \
  --i-accession-ids ncbi-accession-i-ds-177.qza \
  --p- \
  --p-retries 5 \
  --p-n-jobs 1 \
  --p-log-level DEBUG \
  --p-no-restricted-access \
  --o-paired-reads paired-reads-177.qza \
  --o-single-reads XX_single_reads \
  --o-failed-runs XX_failed_runs

qiime fondue get-sequences \
  --i-accession-ids ncbi-accession-i-ds-178.qza \
  --p- \
  --p-retries 5 \
  --p-n-jobs 1 \
  --p-log-level DEBUG \
  --p-no-restricted-access \
  --o-paired-reads paired-reads-178.qza \
  --o-single-reads XX_single_reads \
  --o-failed-runs XX_failed_runs

qiime fondue get-sequences \
  --i-accession-ids ncbi-accession-i-ds-179.qza \
  --p- \
  --p-retries 5 \
  --p-n-jobs 1 \
  --p-log-level DEBUG \
  --p-no-restricted-access \
  --o-paired-reads paired-reads-179.qza \
  --o-single-reads XX_single_reads \
  --o-failed-runs XX_failed_runs

qiime fondue get-sequences \
  --i-accession-ids ncbi-accession-i-ds-180.qza \
  --p- \
  --p-retries 5 \
  --p-n-jobs 1 \
  --p-log-level DEBUG \
  --p-no-restricted-access \
  --o-paired-reads paired-reads-180.qza \
  --o-single-reads XX_single_reads \
  --o-failed-runs XX_failed_runs

qiime fondue get-sequences \
```

```
 --i-accession-ids ncbi-accession-i-ds-181.qza \
 --p- \
 --p-retries 5 \
 --p-n-jobs 1 \
 --p-log-level DEBUG \
 --p-no-restricted-access \
 --o-paired-reads paired-reads-181.qza \
 --o-single-reads XX_single_reads \
 --o-failed-runs XX_failed_runs

qiime fondue get-sequences \
 --i-accession-ids ncbi-accession-i-ds-182.qza \
 --p- \
 --p-retries 5 \
 --p-n-jobs 1 \
 --p-log-level DEBUG \
 --p-no-restricted-access \
 --o-paired-reads paired-reads-182.qza \
 --o-single-reads XX_single_reads \
 --o-failed-runs XX_failed_runs

qiime fondue get-sequences \
 --i-accession-ids ncbi-accession-i-ds-183.qza \
 --p- \
 --p-retries 5 \
 --p-n-jobs 1 \
 --p-log-level DEBUG \
 --p-no-restricted-access \
 --o-paired-reads paired-reads-183.qza \
 --o-single-reads XX_single_reads \
 --o-failed-runs XX_failed_runs

qiime fondue get-sequences \
 --i-accession-ids ncbi-accession-i-ds-184.qza \
 --p- \
 --p-retries 5 \
 --p-n-jobs 1 \
 --p-log-level DEBUG \
 --p-no-restricted-access \
 --o-paired-reads paired-reads-184.qza \
 --o-single-reads XX_single_reads \
 --o-failed-runs XX_failed_runs

qiime fondue get-sequences \
 --i-accession-ids ncbi-accession-i-ds-185.qza \
 --p- \
 --p-retries 5 \
 --p-n-jobs 1 \
 --p-log-level DEBUG \
 --p-no-restricted-access \
 --o-paired-reads paired-reads-185.qza \
 --o-single-reads XX_single_reads \
```

```
  --o-failed-runs XX_failed_runs

qiime fondue get-sequences \
  --i-accession-ids ncbi-accession-i-ds-186.qza \
  --p- \
  --p-retries 5 \
  --p-n-jobs 1 \
  --p-log-level DEBUG \
  --p-no-restricted-access \
  --o-paired-reads paired-reads-186.qza \
  --o-single-reads XX_single_reads \
  --o-failed-runs XX_failed_runs

qiime fondue get-sequences \
  --i-accession-ids ncbi-accession-i-ds-187.qza \
  --p- \
  --p-retries 5 \
  --p-n-jobs 1 \
  --p-log-level DEBUG \
  --p-no-restricted-access \
  --o-paired-reads paired-reads-187.qza \
  --o-single-reads XX_single_reads \
  --o-failed-runs XX_failed_runs

qiime fondue get-sequences \
  --i-accession-ids ncbi-accession-i-ds-188.qza \
  --p- \
  --p-retries 5 \
  --p-n-jobs 1 \
  --p-log-level DEBUG \
  --p-no-restricted-access \
  --o-paired-reads paired-reads-188.qza \
  --o-single-reads XX_single_reads \
  --o-failed-runs XX_failed_runs

qiime fondue get-sequences \
  --i-accession-ids ncbi-accession-i-ds-189.qza \
  --p- \
  --p-retries 5 \
  --p-n-jobs 1 \
  --p-log-level DEBUG \
  --p-no-restricted-access \
  --o-paired-reads paired-reads-189.qza \
  --o-single-reads XX_single_reads \
  --o-failed-runs XX_failed_runs

qiime fondue get-sequences \
  --i-accession-ids ncbi-accession-i-ds-190.qza \
  --p- \
  --p-retries 5 \
  --p-n-jobs 1 \
  --p-log-level DEBUG \
```

```
  --p-no-restricted-access \
  --o-paired-reads paired-reads-190.qza \
  --o-single-reads XX_single_reads \
  --o-failed-runs XX_failed_runs

qiime fondue get-sequences \
  --i-accession-ids ncbi-accession-i-ds-191.qza \
  --p- \
  --p-retries 5 \
  --p-n-jobs 1 \
  --p-log-level DEBUG \
  --p-no-restricted-access \
  --o-paired-reads paired-reads-191.qza \
  --o-single-reads XX_single_reads \
  --o-failed-runs XX_failed_runs

qiime fondue get-sequences \
  --i-accession-ids ncbi-accession-i-ds-192.qza \
  --p- \
  --p-retries 5 \
  --p-n-jobs 1 \
  --p-log-level DEBUG \
  --p-no-restricted-access \
  --o-paired-reads paired-reads-192.qza \
  --o-single-reads XX_single_reads \
  --o-failed-runs XX_failed_runs

qiime fondue get-sequences \
  --i-accession-ids ncbi-accession-i-ds-193.qza \
  --p- \
  --p-retries 5 \
  --p-n-jobs 1 \
  --p-log-level DEBUG \
  --p-no-restricted-access \
  --o-paired-reads paired-reads-193.qza \
  --o-single-reads XX_single_reads \
  --o-failed-runs XX_failed_runs

qiime fondue get-sequences \
  --i-accession-ids ncbi-accession-i-ds-194.qza \
  --p- \
  --p-retries 5 \
  --p-n-jobs 1 \
  --p-log-level DEBUG \
  --p-no-restricted-access \
  --o-paired-reads paired-reads-194.qza \
  --o-single-reads XX_single_reads \
  --o-failed-runs XX_failed_runs

qiime fondue get-sequences \
  --i-accession-ids ncbi-accession-i-ds-195.qza \
  --p- \
```

```
  --p-retries 5 \
  --p-n-jobs 1 \
  --p-log-level DEBUG \
  --p-no-restricted-access \
  --o-paired-reads paired-reads-195.qza \
  --o-single-reads XX_single_reads \
  --o-failed-runs XX_failed_runs

qiime fondue get-sequences \
  --i-accession-ids ncbi-accession-i-ds-196.qza \
  --p- \
  --p-retries 5 \
  --p-n-jobs 1 \
  --p-log-level DEBUG \
  --p-no-restricted-access \
  --o-paired-reads paired-reads-196.qza \
  --o-single-reads XX_single_reads \
  --o-failed-runs XX_failed_runs

qiime fondue get-sequences \
  --i-accession-ids ncbi-accession-i-ds-197.qza \
  --p- \
  --p-retries 5 \
  --p-n-jobs 1 \
  --p-log-level DEBUG \
  --p-no-restricted-access \
  --o-paired-reads paired-reads-197.qza \
  --o-single-reads XX_single_reads \
  --o-failed-runs XX_failed_runs

qiime fondue get-sequences \
  --i-accession-ids ncbi-accession-i-ds-198.qza \
  --p- \
  --p-retries 5 \
  --p-n-jobs 1 \
  --p-log-level DEBUG \
  --p-no-restricted-access \
  --o-paired-reads paired-reads-198.qza \
  --o-single-reads XX_single_reads \
  --o-failed-runs XX_failed_runs

qiime fondue get-sequences \
  --i-accession-ids ncbi-accession-i-ds-199.qza \
  --p- \
  --p-retries 5 \
  --p-n-jobs 1 \
  --p-log-level DEBUG \
  --p-no-restricted-access \
  --o-paired-reads paired-reads-199.qza \
  --o-single-reads XX_single_reads \
  --o-failed-runs XX_failed_runs
```

```
qiime fondue get-sequences \
  --i-accession-ids ncbi-accession-i-ds-200.qza \
  --p- \
  --p-retries 5 \
  --p-n-jobs 1 \
  --p-log-level DEBUG \
  --p-no-restricted-access \
  --o-paired-reads paired-reads-200.qza \
  --o-single-reads XX_single_reads \
  --o-failed-runs XX_failed_runs

qiime fondue get-sequences \
  --i-accession-ids ncbi-accession-i-ds-201.qza \
  --p- \
  --p-retries 5 \
  --p-n-jobs 1 \
  --p-log-level DEBUG \
  --p-no-restricted-access \
  --o-paired-reads paired-reads-201.qza \
  --o-single-reads XX_single_reads \
  --o-failed-runs XX_failed_runs

qiime fondue get-sequences \
  --i-accession-ids ncbi-accession-i-ds-202.qza \
  --p- \
  --p-retries 5 \
  --p-n-jobs 1 \
  --p-log-level DEBUG \
  --p-no-restricted-access \
  --o-paired-reads paired-reads-202.qza \
  --o-single-reads XX_single_reads \
  --o-failed-runs XX_failed_runs

qiime fondue get-sequences \
  --i-accession-ids ncbi-accession-i-ds-203.qza \
  --p- \
  --p-retries 5 \
  --p-n-jobs 1 \
  --p-log-level DEBUG \
  --p-no-restricted-access \
  --o-paired-reads paired-reads-203.qza \
  --o-single-reads XX_single_reads \
  --o-failed-runs XX_failed_runs

qiime fondue get-sequences \
  --i-accession-ids ncbi-accession-i-ds-204.qza \
  --p- \
  --p-retries 5 \
  --p-n-jobs 1 \
  --p-log-level DEBUG \
  --p-no-restricted-access \
  --o-paired-reads paired-reads-204.qza \
```

```
  --o-single-reads XX_single_reads \
  --o-failed-runs XX_failed_runs

qiime fondue get-sequences \
  --i-accession-ids ncbi-accession-i-ds-205.qza \
  --p- \
  --p-retries 5 \
  --p-n-jobs 1 \
  --p-log-level DEBUG \
  --p-no-restricted-access \
  --o-paired-reads paired-reads-205.qza \
  --o-single-reads XX_single_reads \
  --o-failed-runs XX_failed_runs

qiime fondue get-sequences \
  --i-accession-ids ncbi-accession-i-ds-206.qza \
  --p- \
  --p-retries 5 \
  --p-n-jobs 1 \
  --p-log-level DEBUG \
  --p-no-restricted-access \
  --o-paired-reads paired-reads-206.qza \
  --o-single-reads XX_single_reads \
  --o-failed-runs XX_failed_runs

qiime fondue get-sequences \
  --i-accession-ids ncbi-accession-i-ds-207.qza \
  --p- \
  --p-retries 5 \
  --p-n-jobs 1 \
  --p-log-level DEBUG \
  --p-no-restricted-access \
  --o-paired-reads paired-reads-207.qza \
  --o-single-reads XX_single_reads \
  --o-failed-runs XX_failed_runs

qiime fondue get-sequences \
  --i-accession-ids ncbi-accession-i-ds-208.qza \
  --p- \
  --p-retries 5 \
  --p-n-jobs 1 \
  --p-log-level DEBUG \
  --p-no-restricted-access \
  --o-paired-reads paired-reads-208.qza \
  --o-single-reads XX_single_reads \
  --o-failed-runs XX_failed_runs

qiime fondue get-sequences \
  --i-accession-ids ncbi-accession-i-ds-209.qza \
  --p- \
  --p-retries 5 \
  --p-n-jobs 1 \
```

```
  --p-log-level DEBUG \
  --p-no-restricted-access \
  --o-paired-reads paired-reads-209.qza \
  --o-single-reads XX_single_reads \
  --o-failed-runs XX_failed_runs

qiime fondue get-sequences \
  --i-accession-ids ncbi-accession-i-ds-210.qza \
  --p- \
  --p-retries 5 \
  --p-n-jobs 1 \
  --p-log-level DEBUG \
  --p-no-restricted-access \
  --o-paired-reads paired-reads-210.qza \
  --o-single-reads XX_single_reads \
  --o-failed-runs XX_failed_runs

qiime fondue get-sequences \
  --i-accession-ids ncbi-accession-i-ds-211.qza \
  --p- \
  --p-retries 5 \
  --p-n-jobs 1 \
  --p-log-level DEBUG \
  --p-no-restricted-access \
  --o-paired-reads paired-reads-211.qza \
  --o-single-reads XX_single_reads \
  --o-failed-runs XX_failed_runs

qiime fondue get-sequences \
  --i-accession-ids ncbi-accession-i-ds-212.qza \
  --p- \
  --p-retries 5 \
  --p-n-jobs 1 \
  --p-log-level DEBUG \
  --p-no-restricted-access \
  --o-paired-reads paired-reads-212.qza \
  --o-single-reads XX_single_reads \
  --o-failed-runs XX_failed_runs

qiime fondue get-sequences \
  --i-accession-ids ncbi-accession-i-ds-213.qza \
  --p- \
  --p-retries 5 \
  --p-n-jobs 1 \
  --p-log-level DEBUG \
  --p-no-restricted-access \
  --o-paired-reads paired-reads-213.qza \
  --o-single-reads XX_single_reads \
  --o-failed-runs XX_failed_runs

qiime fondue get-sequences \
  --i-accession-ids ncbi-accession-i-ds-214.qza \
```

```
  --p- \
  --p-retries 5 \
  --p-n-jobs 1 \
  --p-log-level DEBUG \
  --p-no-restricted-access \
  --o-paired-reads paired-reads-214.qza \
  --o-single-reads XX_single_reads \
  --o-failed-runs XX_failed_runs

qiime fondue get-sequences \
  --i-accession-ids ncbi-accession-i-ds-215.qza \
  --p- \
  --p-retries 5 \
  --p-n-jobs 1 \
  --p-log-level DEBUG \
  --p-no-restricted-access \
  --o-paired-reads paired-reads-215.qza \
  --o-single-reads XX_single_reads \
  --o-failed-runs XX_failed_runs

qiime fondue get-sequences \
  --i-accession-ids ncbi-accession-i-ds-216.qza \
  --p- \
  --p-retries 5 \
  --p-n-jobs 1 \
  --p-log-level DEBUG \
  --p-no-restricted-access \
  --o-paired-reads paired-reads-216.qza \
  --o-single-reads XX_single_reads \
  --o-failed-runs XX_failed_runs

qiime fondue get-sequences \
  --i-accession-ids ncbi-accession-i-ds-217.qza \
  --p- \
  --p-retries 5 \
  --p-n-jobs 1 \
  --p-log-level DEBUG \
  --p-no-restricted-access \
  --o-paired-reads paired-reads-217.qza \
  --o-single-reads XX_single_reads \
  --o-failed-runs XX_failed_runs

qiime fondue get-sequences \
  --i-accession-ids ncbi-accession-i-ds-218.qza \
  --p- \
  --p-retries 5 \
  --p-n-jobs 1 \
  --p-log-level DEBUG \
  --p-no-restricted-access \
  --o-paired-reads paired-reads-218.qza \
  --o-single-reads XX_single_reads \
  --o-failed-runs XX_failed_runs
```

```
qiime fondue get-sequences \
  --i-accession-ids ncbi-accession-i-ds-219.qza \
  --p- \
  --p-retries 5 \
  --p-n-jobs 1 \
  --p-log-level DEBUG \
  --p-no-restricted-access \
  --o-paired-reads paired-reads-219.qza \
  --o-single-reads XX_single_reads \
  --o-failed-runs XX_failed_runs

qiime fondue get-sequences \
  --i-accession-ids ncbi-accession-i-ds-220.qza \
  --p- \
  --p-retries 5 \
  --p-n-jobs 1 \
  --p-log-level DEBUG \
  --p-no-restricted-access \
  --o-paired-reads paired-reads-220.qza \
  --o-single-reads XX_single_reads \
  --o-failed-runs XX_failed_runs

qiime fondue get-sequences \
  --i-accession-ids ncbi-accession-i-ds-221.qza \
  --p- \
  --p-retries 5 \
  --p-n-jobs 1 \
  --p-log-level DEBUG \
  --p-no-restricted-access \
  --o-paired-reads paired-reads-221.qza \
  --o-single-reads XX_single_reads \
  --o-failed-runs XX_failed_runs

qiime fondue get-sequences \
  --i-accession-ids ncbi-accession-i-ds-222.qza \
  --p- \
  --p-retries 5 \
  --p-n-jobs 1 \
  --p-log-level DEBUG \
  --p-no-restricted-access \
  --o-paired-reads paired-reads-222.qza \
  --o-single-reads XX_single_reads \
  --o-failed-runs XX_failed_runs

qiime fondue get-sequences \
  --i-accession-ids ncbi-accession-i-ds-223.qza \
  --p- \
  --p-retries 5 \
  --p-n-jobs 1 \
  --p-log-level DEBUG \
  --p-no-restricted-access \
```

```
  --o-paired-reads paired-reads-223.qza \
  --o-single-reads XX_single_reads \
  --o-failed-runs XX_failed_runs

qiime fondue get-sequences \
  --i-accession-ids ncbi-accession-i-ds-224.qza \
  --p- \
  --p-retries 5 \
  --p-n-jobs 1 \
  --p-log-level DEBUG \
  --p-no-restricted-access \
  --o-paired-reads paired-reads-224.qza \
  --o-single-reads XX_single_reads \
  --o-failed-runs XX_failed_runs

qiime fondue get-sequences \
  --i-accession-ids ncbi-accession-i-ds-225.qza \
  --p- \
  --p-retries 5 \
  --p-n-jobs 1 \
  --p-log-level DEBUG \
  --p-no-restricted-access \
  --o-paired-reads paired-reads-225.qza \
  --o-single-reads XX_single_reads \
  --o-failed-runs XX_failed_runs

qiime fondue get-sequences \
  --i-accession-ids ncbi-accession-i-ds-226.qza \
  --p- \
  --p-retries 5 \
  --p-n-jobs 1 \
  --p-log-level DEBUG \
  --p-no-restricted-access \
  --o-paired-reads paired-reads-226.qza \
  --o-single-reads XX_single_reads \
  --o-failed-runs XX_failed_runs

qiime fondue get-sequences \
  --i-accession-ids ncbi-accession-i-ds-227.qza \
  --p- \
  --p-retries 5 \
  --p-n-jobs 1 \
  --p-log-level DEBUG \
  --p-no-restricted-access \
  --o-paired-reads paired-reads-227.qza \
  --o-single-reads XX_single_reads \
  --o-failed-runs XX_failed_runs

qiime fondue get-sequences \
  --i-accession-ids ncbi-accession-i-ds-228.qza \
  --p- \
  --p-retries 5 \
```

```
  --p-n-jobs 1 \
  --p-log-level DEBUG \
  --p-no-restricted-access \
  --o-paired-reads paired-reads-228.qza \
  --o-single-reads XX_single_reads \
  --o-failed-runs XX_failed_runs

qiime fondue get-sequences \
  --i-accession-ids ncbi-accession-i-ds-229.qza \
  --p- \
  --p-retries 5 \
  --p-n-jobs 1 \
  --p-log-level DEBUG \
  --p-no-restricted-access \
  --o-paired-reads paired-reads-229.qza \
  --o-single-reads XX_single_reads \
  --o-failed-runs XX_failed_runs

qiime fondue get-sequences \
  --i-accession-ids ncbi-accession-i-ds-230.qza \
  --p- \
  --p-retries 5 \
  --p-n-jobs 1 \
  --p-log-level DEBUG \
  --p-no-restricted-access \
  --o-paired-reads paired-reads-230.qza \
  --o-single-reads XX_single_reads \
  --o-failed-runs XX_failed_runs

qiime fondue get-sequences \
  --i-accession-ids ncbi-accession-i-ds-231.qza \
  --p- \
  --p-retries 5 \
  --p-n-jobs 1 \
  --p-log-level DEBUG \
  --p-no-restricted-access \
  --o-paired-reads paired-reads-231.qza \
  --o-single-reads XX_single_reads \
  --o-failed-runs XX_failed_runs

qiime fondue get-sequences \
  --i-accession-ids ncbi-accession-i-ds-232.qza \
  --p- \
  --p-retries 5 \
  --p-n-jobs 1 \
  --p-log-level DEBUG \
  --p-no-restricted-access \
  --o-paired-reads paired-reads-232.qza \
  --o-single-reads XX_single_reads \
  --o-failed-runs XX_failed_runs

qiime fondue get-sequences \
```

```
  --i-accession-ids ncbi-accession-i-ds-233.qza \
  --p- \
  --p-retries 5 \
  --p-n-jobs 1 \
  --p-log-level DEBUG \
  --p-no-restricted-access \
  --o-paired-reads paired-reads-233.qza \
  --o-single-reads XX_single_reads \
  --o-failed-runs XX_failed_runs

qiime fondue get-sequences \
  --i-accession-ids ncbi-accession-i-ds-234.qza \
  --p- \
  --p-retries 5 \
  --p-n-jobs 1 \
  --p-log-level DEBUG \
  --p-no-restricted-access \
  --o-paired-reads paired-reads-234.qza \
  --o-single-reads XX_single_reads \
  --o-failed-runs XX_failed_runs

qiime fondue get-sequences \
  --i-accession-ids ncbi-accession-i-ds-235.qza \
  --p- \
  --p-retries 5 \
  --p-n-jobs 1 \
  --p-log-level DEBUG \
  --p-no-restricted-access \
  --o-paired-reads paired-reads-235.qza \
  --o-single-reads XX_single_reads \
  --o-failed-runs XX_failed_runs

qiime fondue get-sequences \
  --i-accession-ids ncbi-accession-i-ds-236.qza \
  --p- \
  --p-retries 5 \
  --p-n-jobs 1 \
  --p-log-level DEBUG \
  --p-no-restricted-access \
  --o-paired-reads paired-reads-236.qza \
  --o-single-reads XX_single_reads \
  --o-failed-runs XX_failed_runs

qiime fondue get-sequences \
  --i-accession-ids ncbi-accession-i-ds-237.qza \
  --p- \
  --p-retries 5 \
  --p-n-jobs 1 \
  --p-log-level DEBUG \
  --p-no-restricted-access \
  --o-paired-reads paired-reads-237.qza \
  --o-single-reads XX_single_reads \
```

```
  --o-failed-runs XX_failed_runs

qiime fondue get-sequences \
  --i-accession-ids ncbi-accession-i-ds-238.qza \
  --p- \
  --p-retries 5 \
  --p-n-jobs 1 \
  --p-log-level DEBUG \
  --p-no-restricted-access \
  --o-paired-reads paired-reads-238.qza \
  --o-single-reads XX_single_reads \
  --o-failed-runs XX_failed_runs

qiime fondue get-sequences \
  --i-accession-ids ncbi-accession-i-ds-239.qza \
  --p- \
  --p-retries 5 \
  --p-n-jobs 1 \
  --p-log-level DEBUG \
  --p-no-restricted-access \
  --o-paired-reads paired-reads-239.qza \
  --o-single-reads XX_single_reads \
  --o-failed-runs XX_failed_runs

qiime fondue get-sequences \
  --i-accession-ids ncbi-accession-i-ds-240.qza \
  --p- \
  --p-retries 5 \
  --p-n-jobs 1 \
  --p-log-level DEBUG \
  --p-no-restricted-access \
  --o-paired-reads paired-reads-240.qza \
  --o-single-reads XX_single_reads \
  --o-failed-runs XX_failed_runs

qiime fondue get-sequences \
  --i-accession-ids ncbi-accession-i-ds-241.qza \
  --p- \
  --p-retries 5 \
  --p-n-jobs 1 \
  --p-log-level DEBUG \
  --p-no-restricted-access \
  --o-paired-reads paired-reads-241.qza \
  --o-single-reads XX_single_reads \
  --o-failed-runs XX_failed_runs

qiime fondue get-sequences \
  --i-accession-ids ncbi-accession-i-ds-242.qza \
  --p- \
  --p-retries 5 \
  --p-n-jobs 1 \
  --p-log-level DEBUG \
```

```
  --p-no-restricted-access \
  --o-paired-reads paired-reads-242.qza \
  --o-single-reads XX_single_reads \
  --o-failed-runs XX_failed_runs

qiime fondue combine-seqs \
  --i-seqs paired-reads-105.qza paired-reads-136.qza paired-reads-14.qza paired-reads-89.qza paired-
reads-157.qza paired-reads-108.qza paired-reads-111.qza paired-reads-160.qza paired-reads-73.qza
paired-reads-228.qza paired-reads-119.qza paired-reads-242.qza paired-reads-239.qza paired-reads-
140.qza paired-reads-51.qza paired-reads-197.qza paired-reads-33.qza paired-reads-201.qza paired-
reads-142.qza paired-reads-18.qza paired-reads-128.qza paired-reads-235.qza paired-reads-123.qza
paired-reads-178.qza paired-reads-22.qza paired-reads-39.qza paired-reads-37.qza paired-reads-
96.qza paired-reads-206.qza paired-reads-170.qza paired-reads-200.qza paired-reads-158.qza
paired-reads-227.qza paired-reads-196.qza paired-reads-62.qza paired-reads-192.qza paired-reads-
26.qza paired-reads-186.qza paired-reads-210.qza paired-reads-163.qza paired-reads-9.qza paired-
reads-187.qza paired-reads-95.qza paired-reads-23.qza paired-reads-120.qza paired-reads-79.qza
paired-reads-240.qza paired-reads-180.qza paired-reads-231.qza paired-reads-100.qza paired-reads-
194.qza paired-reads-70.qza paired-reads-106.qza paired-reads-74.qza paired-reads-98.qza paired-
reads-190.qza paired-reads-173.qza paired-reads-50.qza paired-reads-0.qza paired-reads-30.qza
paired-reads-58.qza paired-reads-202.qza paired-reads-67.qza paired-reads-110.qza paired-reads-
185.qza paired-reads-36.qza paired-reads-85.qza paired-reads-11.qza paired-reads-59.qza paired-
reads-152.qza paired-reads-159.qza paired-reads-218.qza paired-reads-47.qza paired-reads-94.qza
paired-reads-53.qza paired-reads-226.qza paired-reads-99.qza paired-reads-174.qza paired-reads-
57.qza paired-reads-131.qza paired-reads-49.qza paired-reads-75.qza paired-reads-129.qza paired-
reads-208.qza paired-reads-48.qza paired-reads-109.qza paired-reads-88.qza paired-reads-86.qza
paired-reads-16.qza paired-reads-156.qza paired-reads-146.qza paired-reads-121.qza paired-reads-
84.qza paired-reads-52.qza paired-reads-219.qza paired-reads-177.qza paired-reads-115.qza paired-
reads-225.qza paired-reads-199.qza paired-reads-8.qza paired-reads-45.qza paired-reads-222.qza
paired-reads-4.qza paired-reads-203.qza paired-reads-232.qza paired-reads-221.qza paired-reads-
189.qza paired-reads-41.qza paired-reads-25.qza paired-reads-167.qza paired-reads-24.qza paired-
reads-72.qza paired-reads-151.qza paired-reads-172.qza paired-reads-77.qza paired-reads-15.qza
paired-reads-28.qza paired-reads-141.qza paired-reads-21.qza paired-reads-132.qza paired-reads-
233.qza paired-reads-213.qza paired-reads-43.qza paired-reads-198.qza paired-reads-211.qza
paired-reads-144.qza paired-reads-147.qza paired-reads-55.qza paired-reads-161.qza paired-reads-
20.qza paired-reads-138.qza paired-reads-92.qza paired-reads-32.qza paired-reads-112.qza paired-
reads-46.qza paired-reads-193.qza paired-reads-216.qza paired-reads-42.qza paired-reads-124.qza
paired-reads-237.qza paired-reads-164.qza paired-reads-117.qza paired-reads-56.qza paired-reads-
54.qza paired-reads-149.qza paired-reads-130.qza paired-reads-103.qza paired-reads-125.qza paired-
reads-223.qza paired-reads-169.qza paired-reads-165.qza paired-reads-182.qza paired-reads-
234.qza paired-reads-150.qza paired-reads-153.qza paired-reads-44.qza paired-reads-1.qza paired-
reads-168.qza paired-reads-38.qza paired-reads-214.qza paired-reads-34.qza paired-reads-139.qza
paired-reads-27.qza paired-reads-102.qza paired-reads-118.qza paired-reads-80.qza paired-reads-
207.qza paired-reads-133.qza paired-reads-35.qza paired-reads-19.qza paired-reads-241.qza paired-
reads-179.qza paired-reads-63.qza paired-reads-69.qza paired-reads-127.qza paired-reads-17.qza
paired-reads-29.qza paired-reads-114.qza paired-reads-83.qza paired-reads-12.qza paired-reads-
65.qza paired-reads-3.qza paired-reads-188.qza paired-reads-116.qza paired-reads-154.qza paired-
reads-183.qza paired-reads-195.qza paired-reads-76.qza paired-reads-68.qza paired-reads-236.qza
paired-reads-2.qza paired-reads-40.qza paired-reads-209.qza paired-reads-212.qza paired-reads-
175.qza paired-reads-134.qza paired-reads-162.qza paired-reads-204.qza paired-reads-220.qza
paired-reads-91.qza paired-reads-229.qza paired-reads-64.qza paired-reads-217.qza paired-reads-
205.qza paired-reads-90.qza paired-reads-215.qza paired-reads-78.qza paired-reads-191.qza paired-
reads-184.qza paired-reads-122.qza paired-reads-135.qza paired-reads-60.qza paired-reads-126.qza
```

```
paired-reads-176.qza paired-reads-97.qza paired-reads-82.qza paired-reads-66.qza paired-reads-7.qza paired-reads-13.qza paired-reads-81.qza paired-reads-107.qza paired-reads-166.qza paired-reads-155.qza paired-reads-238.qza paired-reads-143.qza paired-reads-6.qza paired-reads-113.qza paired-reads-10.qza paired-reads-87.qza paired-reads-171.qza paired-reads-61.qza paired-reads-145.qza paired-reads-104.qza paired-reads-181.qza paired-reads-101.qza paired-reads-5.qza paired-reads-148.qza paired-reads-224.qza paired-reads-31.qza paired-reads-230.qza paired-reads-137.qza paired-reads-93.qza paired-reads-71.qza \
  --p-on-duplicates error \
  --o-combined-seqs combined-seqs-0.qza

qiime cutadapt trim-paired \
  --i-demultiplexed-sequences combined-seqs-0.qza \
  --p-cores 24 \
  --p-error-rate 0.1 \
  --p-indels \
  --p-times 1 \
  --p-overlap 3 \
  --p-no-match-read-wildcards \
  --p-match-adapter-wildcards \
  --p-minimum-length 90 \
  --p-no-discard-untrimmed \
  --p-quality-cutoff-5end 0 \
  --p-quality-cutoff-3end 0 \
  --p-quality-base 33 \
  --o-trimmed-sequences trimmed-sequences-0.qza

qiime moshpit classify-kraken2 \
  --i-seqs trimmed-sequences-0.qza \
  --i-kraken2-db kraken2-database-0.qza \
  --p-threads 72 \
  --p-confidence 0.2 \
  --p-minimum-base-quality 0 \
  --p-no-memory-mapping \
  --p-minimum-hit-groups 2 \
  --p-no-quick \
  --p-report-minimizer-data \
  --o-reports reports-0.qza \
  --o-hits XX_hits

qiime moshpit estimate-bracken \
  --i-kraken-reports reports-0.qza \
  --i-bracken-db bracken-database-0.qza \
  --p-threshold 0 \
  --p-read-len 100 \
  --p-level S \
  --o-table table-0.qza \
  --o-taxonomy taxonomy-0.qza \
  --o-reports XX_reports

qiime taxa filter-table \
  --i-table table-0.qza \
  --i-taxonomy taxonomy-0.qza \
```

```
  --p-exclude Unclassified,sapiens \
  --p-query-delimiter , \
  --p-mode contains \
  --o-filtered-table filtered-table-0.qza

qiime taxa filter-table \
  --i-table table-0.qza \
  --i-taxonomy taxonomy-0.qza \
  --p-exclude Unclassified \
  --p-query-delimiter , \
  --p-mode contains \
  --o-filtered-table filtered-table-1.qza

# Replay attempts to represent metadata inputs accurately, but metadata .tsv
# files are merged automatically by some interfaces, rendering distinctions
# between file inputs invisible in provenance. We output the recorded
# metadata to disk to enable visual inspection.

# The following command may have received additional metadata .tsv files. To
# confirm you have covered your metadata needs adequately, review the
# original metadata, saved at '/var/folders/7f/7nw_x13n5q965rss_qz6061m0000g
# q/T/tmpbnwxdnkd/provenance_replay/recorded_metadata/diversity_core_metrics
# _0'

qiime diversity core-metrics \
  --i-table filtered-table-0.qza \
  --p-sampling-depth 617000 \
  --m-metadata-file <your metadata filepath>.tsv \
  --p-no-with-replacement \
  --p-n-jobs 4 \
  --p-no-ignore-missing-samples \
  --o-bray-curtis-pcoa-results bray-curtis-pcoa-results-0.qza \
  --o-rarefied-table rarefied-table-0.qza \
  --o-observed-features-vector XX_observed_features_vector \
  --o-shannon-vector XX_shannon_vector \
  --o-evenness-vector XX_evenness_vector \
  --o-jaccard-distance-matrix XX_jaccard_distance_matrix \
  --o-bray-curtis-distance-matrix XX_bray_curtis_distance_matrix \
  --o-jaccard-pcoa-results XX_jaccard_pcoa_results \
  --o-jaccard-emperor XX_jaccard_emperor \
  --o-bray-curtis-emperor XX_bray_curtis_emperor

# The following command may have received additional metadata .tsv files. To
# confirm you have covered your metadata needs adequately, review the
# original metadata, saved at '/var/folders/7f/7nw_x13n5q965rss_qz6061m0000g
# q/T/tmpbnwxdnkd/provenance_replay/recorded_metadata/diversity_core_metrics
# _1'

qiime diversity core-metrics \
  --i-table filtered-table-1.qza \
  --p-sampling-depth 696000 \
  --m-metadata-file <your metadata filepath>.tsv \
```

```
 --p-no-with-replacement \
 --p-n-jobs 4 \
 --p-no-ignore-missing-samples \
 --o-shannon-vector shannon-vector-0.qza \
 --o-rarefied-table XX_rarefied_table \
 --o-observed-features-vector XX_observed_features_vector \
 --o-evenness-vector XX_evenness_vector \
 --o-jaccard-distance-matrix XX_jaccard_distance_matrix \
 --o-bray-curtis-distance-matrix XX_bray_curtis_distance_matrix \
 --o-jaccard-pcoa-results XX_jaccard_pcoa_results \
 --o-bray-curtis-pcoa-results XX_bray_curtis_pcoa_results \
 --o-jaccard-emperor XX_jaccard_emperor \
 --o-bray-curtis-emperor XX_bray_curtis_emperor

##################################################################################
# The following QIIME 2 Results were parsed to produce this script:
# 80a8f53e-f453-49de-be39-f3c0fda16c24        9e600572-900b-47ff-b099-30c37baa6c34
# ce29d3aa-2a2e-410b-b33e-47a2f7e5b16d        f5d5c1d9-53bb-4a95-b60b-313dc26758fc
##################################################################################
```

**Cocoa fermentation: provenance replay script**

```
#!/usr/bin/env bash
##########################################################################################
# Auto-generated by qiime2 v.2024.10.1 at 10:59:07 AM on 29 Nov, 2024
# This document is a representation of the scholarly work of the creator of the
# QIIME 2 Results provided as input to this software, and may be protected by
# intellectual property law. Please respect all copyright restrictions and
# licenses governing the use, modification, and redistribution of this work.

# For User Support, post to the QIIME2 Forum at https://forum.qiime2.org.

# Instructions for use:
# 1. Open this script in a text editor or IDE. Support for BASH
#    syntax highlighting can be helpful.
# 2. Search or scan visually for '<' or '>' characters to find places where
#    user input (e.g. a filepath or column name) is required. These must be
#    replaced with your own values. E.g. <column name> -> 'patient_id'.
#    Failure to remove '<' or '>' may result in `No such File ...` errors
# 3. Search for 'FIXME' comments in the script, and respond as directed.
# 4. Remove all 'FIXME' comments from the script completely. Failure to do so
#    may result in 'Missing Option' errors
# 5. Adjust the arguments to the commands below to suit your data and metadata.
#    If your data is not identical to that in the replayed analysis,
#    changes may be required. (e.g. sample ids or rarefaction depth)
# 6. Optional: replace any filenames in this script that begin with 'XX' with
#    unique file names to ensure they are preserved. QIIME 2 saves all outputs
#    from all actions in this script to disk regardless of whether those
#    outputs were in the original collection of replayed results. The filenames
#    of "un-replayed" artifacts are prefixed with 'XX' so they may be easily
#    located. These names are not guaranteed to be unique, so 'XX_table.qza'
#    may be overwritten by another 'XX_table.qza' later in the script.
# 7. Activate your replay conda environment, and confirm you have installed all
#    plugins used by the script.
# 8. Run this script with `bash <path to this script>`, or copy-paste commands
#    into the terminal for a more interactive analysis.
# 9. Optional: to delete all results not required to produce the figures and
#    data used to generate this script, navigate to the directory in which you
#    ran the script and `rm XX*.qz*`
##########################################################################################
## function to create result collections ##
construct_result_collection () {
	mkdir $rc_name
	touch $rc_name.order
	for key in "${keys[@]}"; do
		echo $key >> $rc_name.order
	done
	for i in "${!keys[@]}"; do
		ln -s ../"${names[i]}" $rc_name"${keys[i]}"$ext
	done
}
##

# This tells bash to -e exit immediately if a command fails
```

```
# and -x show all commands in stdout so you can track progress
set -e -x

qiime tools import \
  --type 'NCBIAccessionIDs' \
  --input-path <your data here> \
  --output-path ncbi-accession-i-ds-0.qza

qiime moshpit build-kraken-db \
  --p-collection pluspf \
  --p-threads 1 \
  --p-kmer-len 35 \
  --p-minimizer-len 31 \
  --p-minimizer-spaces 7 \
  --p-no-no-masking \
  --p-max-db-size 0 \
  --p-no-use-ftp \
  --p-load-factor 0.7 \
  --p-no-fast-build \
  --o-kraken2-database kraken2-database-0.qza \
  --o-bracken-database bracken-database-0.qza

qiime moshpit fetch-busco-db \
  --p-virus False \
  --p-prok True \
  --p-euk False \
  --o-busco-db busco-db-0.qza

qiime tools import \
  --type 'FeatureTable[Frequency]' \
  --input-path <your data here> \
  --output-path feature-table-frequency-0.qza

qiime moshpit fetch-kaiju-db \
  --p-database-type nr_euk \
  --o-database database-0.qza

qiime tools import \
  --type 'ReferenceDB[Eggnog]' \
  --input-path <your data here> \
  --output-path reference-db-eggnog-0.qza

qiime tools import \
  --type 'ReferenceDB[Diamond]' \
  --input-path <your data here> \
  --output-path reference-db-diamond-0.qza

qiime fondue get-all \
  --i-accession-ids ncbi-accession-i-ds-0.qza \
  --p- \
  --p-n-jobs 5 \
  --p-retries 5 \
```

```
  --p-log-level DEBUG \
  --o-paired-reads paired-reads-0.qza \
  --o-metadata XX_metadata \
  --o-single-reads XX_single_reads \
  --o-failed-runs XX_failed_runs

qiime moshpit classify-kraken2 \
  --i-seqs paired-reads-0.qza \
  --i-kraken2-db kraken2-database-0.qza \
  --p-threads 72 \
  --p-confidence 0.5 \
  --p-minimum-base-quality 0 \
  --p-no-memory-mapping \
  --p-minimum-hit-groups 2 \
  --p-no-quick \
  --p-report-minimizer-data \
  --o-reports reports-0.qza \
  --o-hits XX_hits

qiime assembly assemble-megahit \
  --i-seqs paired-reads-0.qza \
  --p-presets meta-sensitive \
  --p-min-count 2 \
  --p-k-list 21 29 39 59 79 99 119 141 \
  --p-no-no-mercy \
  --p-bubble-level 2 \
  --p-prune-level 2 \
  --p-prune-depth 2 \
  --p-disconnect-ratio 0.1 \
  --p-low-local-ratio 0.2 \
  --p-max-tip-len auto \
  --p-cleaning-rounds 5 \
  --p-no-no-local \
  --p-no-kmin-1pass \
  --p-memory 0.9 \
  --p-mem-flag 1 \
  --p-num-cpu-threads 24 \
  --p-no-no-hw-accel \
  --p-min-contig-len 200 \
  --p-coassemble False \
  --o-contigs contigs-0.qza

qiime moshpit classify-kaiju \
  --i-seqs paired-reads-0.qza \
  --i-db database-0.qza \
  --p-z 16 \
  --p-a greedy \
  --p-e 3 \
  --p-m 11 \
  --p-s 65 \
  --p-evalue 0.01 \
  --p-x \
```

```
  --p-r species \
  --p-c 0.1 \
  --p-no-exp \
  --p-no-u \
  --o-taxonomy taxonomy-0.qza \
  --o-abundances XX_abundances

qiime motus profile \
  --i-samples paired-reads-0.qza \
  --p-threads 4 \
  --p-min-alen 75 \
  --p-marker-gene-cutoff 3 \
  --p-mode insert.scaled_counts \
  --p-no-reference-genomes \
  --p-jobs 4 \
  --o-taxonomy taxonomy-1.qza \
  --o-table table-0.qza

qiime moshpit estimate-bracken \
  --i-kraken-reports reports-0.qza \
  --i-bracken-db bracken-database-0.qza \
  --p-threshold 5 \
  --p-read-len 150 \
  --p-level S \
  --o-taxonomy taxonomy-2.qza \
  --o-table table-1.qza \
  --o-reports XX_reports

qiime assembly index-contigs \
  --i-contigs contigs-0.qza \
  --p-no-large-index \
  --p-no-debug \
  --p-no-sanitized \
  --p-verbose \
  --p-no-noauto \
  --p-no-packed \
  --p-bmax auto \
  --p-bmaxdivn 4 \
  --p-dcv 1024 \
  --p-no-nodc \
  --p-offrate 5 \
  --p-ftabchars 10 \
  --p-threads 8 \
  --p-seed 100 \
  --o-index index-0.qza

qiime taxa filter-table \
  --i-table feature-table-frequency-0.qza \
  --i-taxonomy taxonomy-0.qza \
  --p-exclude unclassified,belong,cannot \
  --p-query-delimiter , \
  --p-mode contains \
```

```
  --o-filtered-table filtered-table-0.qza

qiime taxa filter-table \
  --i-table table-0.qza \
  --i-taxonomy taxonomy-1.qza \
  --p-exclude 'u; n; a; s; s; i; g; n; e; d' \
  --p-query-delimiter , \
  --p-mode contains \
  --o-filtered-table filtered-table-1.qza

qiime taxa filter-table \
  --i-table table-1.qza \
  --i-taxonomy taxonomy-2.qza \
  --p-exclude Unclassified \
  --p-query-delimiter , \
  --p-mode contains \
  --o-filtered-table filtered-table-2.qza

qiime assembly map-reads \
  --i-index index-0.qza \
  --i-reads paired-reads-0.qza \
  --p-skip 0 \
  --p-qupto unlimited \
  --p-trim5 0 \
  --p-trim3 0 \
  --p-trim-to untrimmed \
  --p-no-phred33 \
  --p-no-phred64 \
  --p-mode local \
  --p-sensitivity sensitive \
  --p-n 0 \
  --p-len 22 \
  --p-i S,1,1.15 \
  --p-n-ceil L,0,0.15 \
  --p-dpad 15 \
  --p-gbar 4 \
  --p-no-ignore-quals \
  --p-no-nofw \
  --p-no-norc \
  --p-no-no-1mm-upfront \
  --p-no-end-to-end \
  --p-no-local \
  --p-ma 2 \
  --p-mp 6 \
  --p-np 1 \
  --p-rdg 5,3 \
  --p-rfg 5,3 \
  --p-k off \
  --p-no-a \
  --p-d 15 \
  --p-r 2 \
  --p-minins 0 \
```

```
  --p-maxins 500 \
  --p-valid-mate-orientations fr \
  --p-no-no-mixed \
  --p-no-no-discordant \
  --p-no-dovetail \
  --p-no-no-contain \
  --p-no-no-overlap \
  --p-offrate off \
  --p-threads 12 \
  --p-no-reorder \
  --p-no-mm \
  --p-seed 100 \
  --p-no-non-deterministic \
  --o-alignment-map alignment-map-0.qza

# Replay attempts to represent metadata inputs accurately, but metadata .tsv
# files are merged automatically by some interfaces, rendering distinctions
# between file inputs invisible in provenance. We output the recorded
# metadata to disk to enable visual inspection.

# The following command may have received additional metadata .tsv files. To
# confirm you have covered your metadata needs adequately, review the
# original metadata, saved at '/var/folders/7f/7nw_x13n5q965rss_qz6061m0000g
# q/T/tmpqp6st2e1/provenance_replay/recorded_metadata/diversity_core_metrics
# _0'

qiime diversity core-metrics \
  --i-table filtered-table-0.qza \
  --p-sampling-depth 5298000 \
  --m-metadata-file <your metadata filepath>.tsv \
  --p-no-with-replacement \
  --p-n-jobs 6 \
  --p-no-ignore-missing-samples \
  --o-shannon-vector shannon-vector-0.qza \
  --o-rarefied-table XX_rarefied_table \
  --o-observed-features-vector XX_observed_features_vector \
  --o-evenness-vector XX_evenness_vector \
  --o-jaccard-distance-matrix XX_jaccard_distance_matrix \
  --o-bray-curtis-distance-matrix XX_bray_curtis_distance_matrix \
  --o-jaccard-pcoa-results XX_jaccard_pcoa_results \
  --o-bray-curtis-pcoa-results XX_bray_curtis_pcoa_results \
  --o-jaccard-emperor XX_jaccard_emperor \
  --o-bray-curtis-emperor XX_bray_curtis_emperor

# The following command may have received additional metadata .tsv files. To
# confirm you have covered your metadata needs adequately, review the
# original metadata, saved at '/var/folders/7f/7nw_x13n5q965rss_qz6061m0000g
# q/T/tmpqp6st2e1/provenance_replay/recorded_metadata/diversity_core_metrics
# _1'

qiime diversity core-metrics \
  --i-table filtered-table-1.qza \
```

```
  --p-sampling-depth 1792 \
  --m-metadata-file <your metadata filepath>.tsv \
  --p-no-with-replacement \
  --p-n-jobs 6 \
  --p-no-ignore-missing-samples \
  --o-shannon-vector shannon-vector-1.qza \
  --o-rarefied-table XX_rarefied_table \
  --o-observed-features-vector XX_observed_features_vector \
  --o-evenness-vector XX_evenness_vector \
  --o-jaccard-distance-matrix XX_jaccard_distance_matrix \
  --o-bray-curtis-distance-matrix XX_bray_curtis_distance_matrix \
  --o-jaccard-pcoa-results XX_jaccard_pcoa_results \
  --o-bray-curtis-pcoa-results XX_bray_curtis_pcoa_results \
  --o-jaccard-emperor XX_jaccard_emperor \
  --o-bray-curtis-emperor XX_bray_curtis_emperor

# The following command may have received additional metadata .tsv files. To
# confirm you have covered your metadata needs adequately, review the
# original metadata, saved at '/var/folders/7f/7nw_x13n5q965rss_qz6061m0000g
# q/T/tmpqp6st2e1/provenance_replay/recorded_metadata/diversity_core_metrics
# _2'

qiime diversity core-metrics \
  --i-table filtered-table-2.qza \
  --p-sampling-depth 2320000 \
  --m-metadata-file <your metadata filepath>.tsv \
  --p-no-with-replacement \
  --p-n-jobs 6 \
  --p-no-ignore-missing-samples \
  --o-rarefied-table rarefied-table-0.qza \
  --o-shannon-vector shannon-vector-2.qza \
  --o-observed-features-vector XX_observed_features_vector \
  --o-evenness-vector XX_evenness_vector \
  --o-jaccard-distance-matrix XX_jaccard_distance_matrix \
  --o-bray-curtis-distance-matrix XX_bray_curtis_distance_matrix \
  --o-jaccard-pcoa-results XX_jaccard_pcoa_results \
  --o-bray-curtis-pcoa-results XX_bray_curtis_pcoa_results \
  --o-jaccard-emperor XX_jaccard_emperor \
  --o-bray-curtis-emperor XX_bray_curtis_emperor

qiime moshpit bin-contigs-metabat \
  --i-contigs contigs-0.qza \
  --i-alignment-maps alignment-map-0.qza \
  --p-num-threads 64 \
  --p-seed 100 \
  --p-verbose \
  --o-mags mags-0.qza \
  --o-contig-map XX_contig_map \
  --o-unbinned-contigs XX_unbinned_contigs

qiime moshpit evaluate-busco \
  --i-bins mags-0.qza \
```

```
  --i-busco-db busco-db-0.qza \
  --p-mode genome \
  --p-lineage-dataset bacteria_odb10 \
  --p-no-augustus \
  --p-auto-lineage False \
  --p-auto-lineage-euk False \
  --p-auto-lineage-prok False \
  --p-cpu 16 \
  --p-contig-break 10 \
  --p-evalue 0.001 \
  --p-no-force \
  --p-limit 3 \
  --p-no-long \
  --p-no-miniprot \
  --p-no-scaffold-composition \
  --o-results-table results-table-0.qza \
  --o-visualization XX_visualization

# The following command may have received additional metadata .tsv files. To
# confirm you have covered your metadata needs adequately, review the
# original metadata, saved at '/var/folders/7f/7nw_x13n5q965rss_qz6061m0000g
# q/T/tmpqp6st2e1/provenance_replay/recorded_metadata/moshpit_filter_mags_0'

qiime moshpit filter-mags \
  --i-mags mags-0.qza \
  --m-metadata-file results-table-0.qza \
  --p-where 'complete>50' \
  --p-no-exclude-ids \
  --p-on mag \
  --o-filtered-mags filtered-mags-0.qza

# The following command may have received additional metadata .tsv files. To
# confirm you have covered your metadata needs adequately, review the
# original metadata, saved at '/var/folders/7f/7nw_x13n5q965rss_qz6061m0000g
# q/T/tmpqp6st2e1/provenance_replay/recorded_metadata/moshpit_filter_mags_1'

qiime moshpit filter-mags \
  --i-mags mags-0.qza \
  --m-metadata-file results-table-0.qza \
  --p-where 'complete>90' \
  --p-no-exclude-ids \
  --p-on mag \
  --o-filtered-mags filtered-mags-1.qza

qiime sourmash compute \
  --i-sequence-file filtered-mags-0.qza \
  --p-ksizes 35 \
  --p-scaled 10 \
  --p-track-abundance \
  --o-min-hash-signature min-hash-signature-0.qza

qiime sourmash compute \
```

```
  --i-sequence-file filtered-mags-1.qza \
  --p-ksizes 35 \
  --p-scaled 10 \
  --p-track-abundance \
  --o-min-hash-signature min-hash-signature-1.qza

qiime sourmash compare \
  --i-min-hash-signature min-hash-signature-0.qza \
  --p-ksize 35 \
  --p-ignore-abundance \
  --o-compare-output compare-output-0.qza

qiime sourmash compare \
  --i-min-hash-signature min-hash-signature-1.qza \
  --p-ksize 35 \
  --p-ignore-abundance \
  --o-compare-output compare-output-1.qza

qiime moshpit dereplicate-mags \
  --i-mags filtered-mags-0.qza \
  --i-distance-matrix compare-output-0.qza \
  --p-threshold 0.99 \
  --o-dereplicated-mags dereplicated-mags-0.qza \
  --o-feature-table XX_feature_table

qiime moshpit dereplicate-mags \
  --i-mags filtered-mags-1.qza \
  --i-distance-matrix compare-output-1.qza \
  --p-threshold 0.99 \
  --o-dereplicated-mags dereplicated-mags-1.qza \
  --o-feature-table XX_feature_table

qiime moshpit get-feature-lengths \
  --i-features dereplicated-mags-0.qza \
  --o-lengths lengths-0.qza

qiime assembly index-derep-mags \
  --i-mags dereplicated-mags-0.qza \
  --p-no-large-index \
  --p-no-debug \
  --p-no-sanitized \
  --p-verbose \
  --p-no-noauto \
  --p-no-packed \
  --p-bmax auto \
  --p-bmaxdivn 4 \
  --p-dcv 1024 \
  --p-no-nodc \
  --p-offrate 5 \
  --p-ftabchars 10 \
  --p-threads 8 \
  --p-seed 100 \
```

```
  --o-index index-1.qza

qiime moshpit classify-kraken2 \
  --i-seqs dereplicated-mags-0.qza \
  --i-kraken2-db kraken2-database-0.qza \
  --p-threads 72 \
  --p-confidence 0.5 \
  --p-minimum-base-quality 0 \
  --p-no-memory-mapping \
  --p-minimum-hit-groups 2 \
  --p-no-quick \
  --p-report-minimizer-data \
  --o-hits hits-0.qza \
  --o-reports reports-1.qza

qiime assembly index-derep-mags \
  --i-mags dereplicated-mags-1.qza \
  --p-no-large-index \
  --p-no-debug \
  --p-no-sanitized \
  --p-verbose \
  --p-no-noauto \
  --p-no-packed \
  --p-bmax auto \
  --p-bmaxdivn 4 \
  --p-dcv 1024 \
  --p-no-nodc \
  --p-offrate 5 \
  --p-ftabchars 10 \
  --p-threads 8 \
  --p-seed 100 \
  --o-index index-2.qza

qiime moshpit eggnog-diamond-search \
  --i-sequences dereplicated-mags-1.qza \
  --i-diamond-db reference-db-diamond-0.qza \
  --p-num-cpus 16 \
  --p-db-in-memory \
  --o-eggnog-hits eggnog-hits-0.qza \
  --o-table XX_table

qiime moshpit get-feature-lengths \
  --i-features dereplicated-mags-1.qza \
  --o-lengths lengths-1.qza

qiime assembly map-reads \
  --i-index index-1.qza \
  --i-reads paired-reads-0.qza \
  --p-skip 0 \
  --p-qupto unlimited \
  --p-trim5 0 \
  --p-trim3 0 \
```

```
  --p-trim-to untrimmed \
  --p-no-phred33 \
  --p-no-phred64 \
  --p-mode local \
  --p-sensitivity sensitive \
  --p-n 0 \
  --p-len 22 \
  --p-i S,1,1.15 \
  --p-n-ceil L,0,0.15 \
  --p-dpad 15 \
  --p-gbar 4 \
  --p-no-ignore-quals \
  --p-no-nofw \
  --p-no-norc \
  --p-no-no-1mm-upfront \
  --p-no-end-to-end \
  --p-no-local \
  --p-ma 2 \
  --p-mp 6 \
  --p-np 1 \
  --p-rdg 5,3 \
  --p-rfg 5,3 \
  --p-k off \
  --p-no-a \
  --p-d 15 \
  --p-r 2 \
  --p-minins 0 \
  --p-maxins 500 \
  --p-valid-mate-orientations fr \
  --p-no-no-mixed \
  --p-no-no-discordant \
  --p-no-dovetail \
  --p-no-no-contain \
  --p-no-no-overlap \
  --p-offrate off \
  --p-threads 12 \
  --p-no-reorder \
  --p-no-mm \
  --p-seed 100 \
  --p-no-non-deterministic \
  --o-alignment-map alignment-map-1.qza

qiime moshpit kraken2-to-mag-features \
  --i-reports reports-1.qza \
  --i-hits hits-0.qza \
  --p-coverage-threshold 0.1 \
  --o-taxonomy taxonomy-3.qza

qiime assembly map-reads \
  --i-index index-2.qza \
  --i-reads paired-reads-0.qza \
  --p-skip 0 \
```

```
  --p-qupto unlimited \
  --p-trim5 0 \
  --p-trim3 0 \
  --p-trim-to untrimmed \
  --p-no-phred33 \
  --p-no-phred64 \
  --p-mode local \
  --p-sensitivity sensitive \
  --p-n 0 \
  --p-len 22 \
  --p-i S,1,1.15 \
  --p-n-ceil L,0,0.15 \
  --p-dpad 15 \
  --p-gbar 4 \
  --p-no-ignore-quals \
  --p-no-nofw \
  --p-no-norc \
  --p-no-no-1mm-upfront \
  --p-no-end-to-end \
  --p-no-local \
  --p-ma 2 \
  --p-mp 6 \
  --p-np 1 \
  --p-rdg 5,3 \
  --p-rfg 5,3 \
  --p-k off \
  --p-no-a \
  --p-d 15 \
  --p-r 2 \
  --p-minins 0 \
  --p-maxins 500 \
  --p-valid-mate-orientations fr \
  --p-no-no-mixed \
  --p-no-no-discordant \
  --p-no-dovetail \
  --p-no-no-contain \
  --p-no-no-overlap \
  --p-offrate off \
  --p-threads 12 \
  --p-no-reorder \
  --p-no-mm \
  --p-seed 100 \
  --p-no-non-deterministic \
  --o-alignment-map alignment-map-2.qza

qiime moshpit eggnog-annotate \
  --i-eggnog-hits eggnog-hits-0.qza \
  --i-eggnog-db reference-db-eggnog-0.qza \
  --p-db-in-memory \
  --p-num-cpus 16 \
  --o-ortholog-annotations ortholog-annotations-0.qza
```

```
qiime moshpit estimate-mag-abundance \
  --i-maps alignment-map-1.qza \
  --i-mag-lengths lengths-0.qza \
  --p-metric rpkm \
  --p-min-mapq 42 \
  --p-min-query-len 0 \
  --p-min-base-quality 20 \
  --p-min-read-len 0 \
  --p-threads 24 \
  --o-abundances abundances-0.qza

qiime moshpit estimate-mag-abundance \
  --i-maps alignment-map-2.qza \
  --i-mag-lengths lengths-1.qza \
  --p-metric rpkm \
  --p-min-mapq 42 \
  --p-min-query-len 0 \
  --p-min-base-quality 20 \
  --p-min-read-len 0 \
  --p-threads 24 \
  --o-abundances abundances-1.qza

qiime moshpit extract-annotations \
  --i-ortholog-annotations ortholog-annotations-0.qza \
  --p-annotation caz \
  --p-max-evalue 0.0001 \
  --p-min-score 0.0 \
  --o-annotation-frequency annotation-frequency-0.qza

# The following command may have received additional metadata .tsv files. To
# confirm you have covered your metadata needs adequately, review the
# original metadata, saved at '/var/folders/7f/7nw_x13n5q965rss_qz6061m0000g
# q/T/tmpqp6st2e1/provenance_replay/recorded_metadata/diversity_core_metrics
# _3'

qiime diversity core-metrics \
  --i-table abundances-0.qza \
  --p-sampling-depth 210 \
  --m-metadata-file <your metadata filepath>.tsv \
  --p-no-with-replacement \
  --p-n-jobs 6 \
  --p-no-ignore-missing-samples \
  --o-shannon-vector shannon-vector-3.qza \
  --o-rarefied-table rarefied-table-1.qza \
  --o-observed-features-vector XX_observed_features_vector \
  --o-evenness-vector XX_evenness_vector \
  --o-jaccard-distance-matrix XX_jaccard_distance_matrix \
  --o-bray-curtis-distance-matrix XX_bray_curtis_distance_matrix \
  --o-jaccard-pcoa-results XX_jaccard_pcoa_results \
  --o-bray-curtis-pcoa-results XX_bray_curtis_pcoa_results \
  --o-jaccard-emperor XX_jaccard_emperor \
  --o-bray-curtis-emperor XX_bray_curtis_emperor
```

```
qiime moshpit multiply-tables \
  --i-table1 abundances-1.qza \
  --i-table2 annotation-frequency-0.qza \
  --o-result-table result-table-0.qza

# The following command may have received additional metadata .tsv files. To
# confirm you have covered your metadata needs adequately, review the
# original metadata, saved at '/var/folders/7f/7nw_x13n5q965rss_qz6061m0000g
# q/T/tmpqp6st2e1/provenance_replay/recorded_metadata/diversity_core_metrics
# _4'

qiime diversity core-metrics \
  --i-table result-table-0.qza \
  --p-sampling-depth 8000 \
  --m-metadata-file <your metadata filepath>.tsv \
  --p-no-with-replacement \
  --p-n-jobs 6 \
  --p-no-ignore-missing-samples \
  --o-rarefied-table rarefied-table-2.qza \
  --o-shannon-vector shannon-vector-4.qza \
  --o-observed-features-vector XX_observed_features_vector \
  --o-evenness-vector XX_evenness_vector \
  --o-jaccard-distance-matrix XX_jaccard_distance_matrix \
  --o-bray-curtis-distance-matrix XX_bray_curtis_distance_matrix \
  --o-jaccard-pcoa-results XX_jaccard_pcoa_results \
  --o-bray-curtis-pcoa-results XX_bray_curtis_pcoa_results \
  --o-jaccard-emperor XX_jaccard_emperor \
  --o-bray-curtis-emperor XX_bray_curtis_emperor

###############################################################################
# The following QIIME 2 Results were parsed to produce this script:
# 137046cb-fe18-4d0b-babc-de209fb49e65      2b5122a9-0396-4ab7-966b-f66fa8a00705
# 5a0bf237-2368-4fe7-9901-50d9d6d92dfb      6cc70426-844c-47ec-82ee-9be5a2d90cda
# 7f0e9ecb-01f7-4a4c-b38f-0bee505d5f76      af746e01-9abf-4d0f-89bb-8abe68b38705
# b6ac8052-ee05-4029-b871-6b2202641a86      bf3224d3-1051-47f4-b79f-c1997f22e1b6
# ca4c6fe6-66a7-41ca-b434-bab6f1cc6d18      e2c284d9-3b8b-4de8-8c19-dc090a0a07e0
###############################################################################
```
